# Estimation and Validation of Individualized Dynamic Brain Models with Resting State fMRI

**DOI:** 10.1101/678243

**Authors:** Matthew F. Singh, Todd S. Braver, Michael W. Cole, ShiNung Ching

## Abstract

A key challenge for neuroscience is to develop generative, causal models of the human nervous system in an individualized, data-driven manner. Previous initiatives have either constructed biologically-plausible models that are not constrained by individual-level human brain activity or used data-driven statistical characterizations of individuals that are not mechanistic. We aim to bridge this gap through the development of a new modeling approach termed Mesoscale Individualized Neurodynamic (MINDy) modeling, wherein we fit nonlinear dynamical systems models directly to human brain imaging data. The MINDy framework is able to produce these data-driven network models for hundreds to thousands of interacting brain regions in just 1-3 minutes per subject. We demonstrate that the models are valid, reliable, and robust. We show that MINDy models are predictive of individualized patterns of resting-state brain dynamical activity. Furthermore, MINDy is better able to uncover the mechanisms underlying individual differences in resting state activity than functional connectivity methods.

## 1. Introduction

To understand human brain function, it is necessary to understand the spatial and temporal computations that govern how its components interact. This understanding can take multiple levels, ranging from statistical descriptions of correlations between brain regions to generative models, which provide a formal mathematical description of how brain activity evolves in time. However, efforts have taken quite different approaches based upon what data is available in human vs. nonhuman subjects. Several international neuroscience initiatives have relied upon nonhuman subjects to collect vast amounts of anatomical and electrophysiological data at the cellular scale ([1], [2], [3]). Generative models are then formed by integrating these cellular-level observations with known neuronal biophysics at the spatial scale of individual neurons or small populations ([1],[2]).

In contrast, another set of large initiatives has instead focused on modeling individual human brain function using an approach often referred to as “connectomics” (e.g., Human Connectome Project, [4]). This approach relies on descriptive statistics, typically correlation between fluctuating activity signals in brain regions assessed during the resting state (“resting state functional connectivity” or rsFC; [5]). As a result, it is sometimes difficult to make mechanistic inferences based upon functional connectivity correlations ([6]). Moreover, neural processes are notoriously nonlinear and inherently dynamic, meaning that stationary descriptions, such as correlation/functional connectivity, may be unable to fully capture brain mechanisms. Nevertheless, rsFC remains the dominant framework for describing connectivity patterns in individual human brains.

Despite the promise of human connectomics, there have been only a few attempts to equip human fMRI studies with the sorts of generative neural population models that have powered insights into non-human nervous systems. Notable advances have occurred in direct-parameterization approaches, with methods being developed to identify directed, causal influences between brain regions (e.g. [7]). Conversely, neural mass modeling approaches have also been extended to study human brain activity in a generative fashion ([8]), and these have provided new insights into the computational mechanisms underlying fMRI and MEG/EEG activity dynamics ([9],[10],[11],[12],[13]). However, unlike (linear) data-driven approaches (e.g. Dynamic Causal Modeling; [14],[7]), neural mass models have been limited to replicating higher-level statistical summaries, such as functional connectivity, rather than predicting the actual time-series. This fact may not be relevant for some applications in which statistical descriptions will suffice. However, there remain many applications in basic neuroscience, neural medicine, and neural engineering for which more precise descriptions could be profitably leveraged.

Unfortunately, current approaches of both types have important limitations. In particular, the existing approaches to directly parameterize models (e.g. DCM) are subject to potential misinferences due to assumptions of linearity ([15]), and, in some cases, limitation to a relatively small number of brain regions ([16],[17],[18]). This number has increased dramatically in recent years by assuming a fixed hemodynamic response function ([19]), but remains well below modern brain parcellations, which feature several hundred regions (e.g. [20], [21], See Discussion). Likewise, with current neural mass modeling approaches, their ability to quantitatively recreate key features of individual-level functional connectivity has also been limited ([22],[12],[13]). This may be because the most common approach is to parameterize connectivity from estimates of white matter integrity from diffusion imaging, which also can lead to potential misinference, since these connectivity estimates are constrained to be symmetric and positive ([23]). Efforts have been made to personalize these models by using individualized diffusion imaging data rather than group-average and/or tuning a small number of free-parameters to better approximate each subject’s summary statistics (e.g. [10],[11][12]). However, again, these models are not directly inferred from the brain activity time-series, which could limit their ability to accurately simulate the dynamical features of these time-series. Indeed, up until this point, it has not been shown that individual-level brain models can be directly parameterized and fit from fMRI while retaining sufficient complexity to capture – and predict – whole-brain activity. This limitation is critical because in order to accurately characterize individual variation in humans – which is the goal of personalized neuroscience and precision medicine initiatives ([24],[25],[26]) – individualized whole-brain models are required.

In the current work, we aim to fill this gap, by advancing high-resolution characterization of the human connectome through the parameterization of nonlinear dynamical systems models that go beyond statistical correlation matrices. The models consist of hundreds of interacting neural populations, each of which is modeled as an abstracted neural mass model evolving over time-scales commensurate with fMRI. Most critically, the models are optimized to capture brain activity dynamics at the level of individual human subjects. We present a computationally efficient algorithm to rapidly fit these models directly from human resting-state fMRI. The algorithm extends data-driven techniques towards the estimation of biologically interpretable models, and conversely enables the parameterization of dynamical neural models in a data-driven, individualized fashion with relatively few priors on the dynamics within and between brain regions. Our approach represents a significant departure and alternative approach to that of previous modeling efforts, in that every parameter in our model is individually estimated without consideration of prior anatomical constraints or long-term summary statistics.

We describe our efforts to develop and validate these models, demonstrating that they successfully characterize whole-brain activity dynamics at the individual level, and as such can be used as a powerful alternative to rsFC, and even to more closely related modeling approaches, such as DCM. Because of this goal, we term our modeling approach MINDy: Mesoscale Individualized Neural Dynamics. In the sections below, we introduce the MINDy modeling framework, highlighting its most innovative and powerful features, and presenting results that validate its utility as an analytic tool for investigating the neural mechanisms and individual differences present in fMRI data.

## 2. Methods

### 2.1. Nature of Interpretations from the Model

The key premise of our approach is an expansion of the architectural description of brain networks from a simple connectivity matrix, to an interpretable dynamical model:

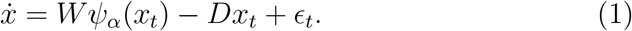

This model, which resembles a neural mass model ([31], [27], [32], [8]) describes the evolution of brain activity at each anatomical location (each element of the vector *x*_*t*_). Unlike true neural-mass models, we model abstracted brain activity commensurate with the fMRI timescale, rather than the evolution of population firing rate over milliseconds. Our model is similar, however, in that it is described by three components: a weight matrix (*W*) which identifies pathways of causal influence between neural populations, a parameterized sigmoidal transfer function (*ψ*) which describes the relation between the local activity of a population and its output to other brain regions (Eq. 3, Fig. 1 A, [33]), and a diagonal decay matrix (*D*) which describes how quickly a given neural population will return to its baseline state after being excited (i.e. the time-constant; Fig. 1 A). Process noise is denoted *ϵ*_*t*_ and is assumed to be uncorrelated between parcels. The additional parameters (*α* and *D*) reflect regional variation in intrinsic dynamics (*D*) and efferent signaling (*α*); critically, as described below, these parameters also show consistent anatomical distributions. These properties vary with brain network and are consistent even at the finer within-network scale (Fig. 4 A,B). Thus, our model, like a neural mass model, parameterizes both the interactions between brain regions and the processes that are local to each brain region that make it distinct.

**Fig 1.**
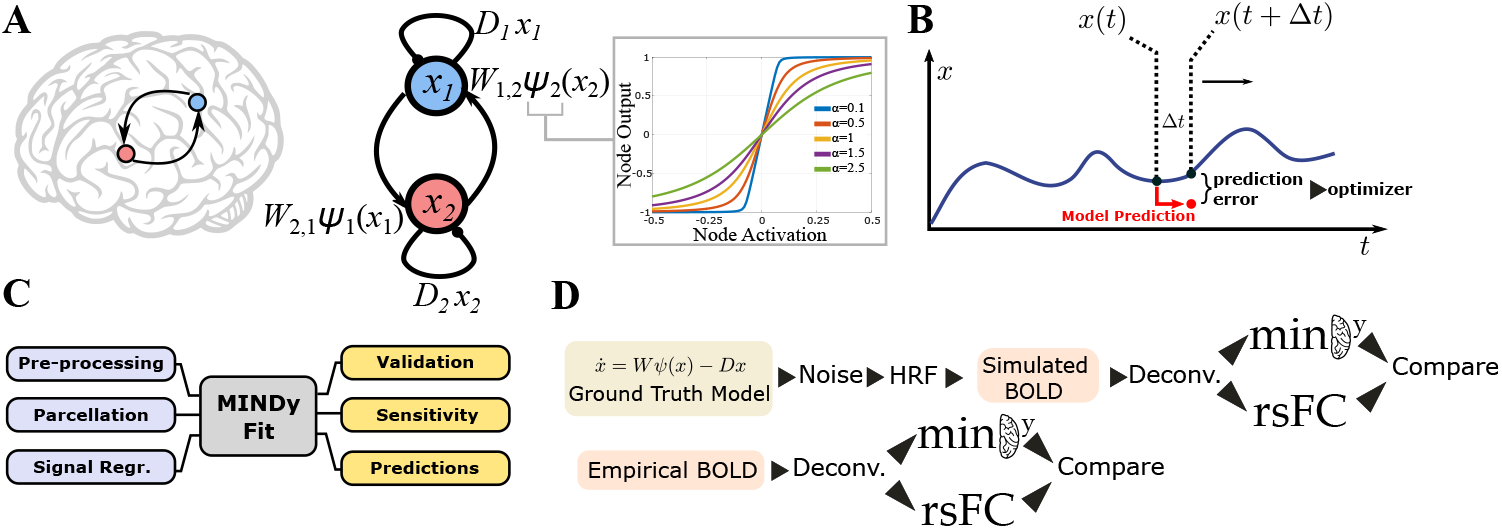
Overview of Methods Employed. A) The MINDy model consists of coupled 1-dimensional neural-mass models (Hopfield form [27]). The shape of the transfer-function for each brain region is parameterized by a curvature parameter *α*. B) Model goodness-of-fit was measured through one-step prediction of the empirical time-series. C) Overview of data processing and analyses: data was processed according to Siegel and colleagues ([28]) and parcellated. Reported analyses fall into three categories: validation, sensitivity to nuissance parameters, and predictions of brain activity patterns. D) In both simulations and empirical analyses the BOLD signal was Wiener-deconvolved ([29]) with a canonical HRF function (see Methods; [30]) before being analyzed with either MINDy or rsFC.

It is important to recognize that this model is a phenomenological model in the sense that the state variables are more abstract than encountered in traditional mean-field models which combine biophysical first-principles and phenomenological approximations (e.g. the sigmoidal nonlinearity). Thus, inferences gained from the model are bounded by the inherent limitations of fMRI data (e.g. low temporal resolution and the indirectness of BOLD). The parametric form that we have chosen leads itself to interpretability. However, we stress that interpretability should not be confused with biophysical equivalence. As described in SI, there are likely many biophysical processes (including non-neuronal) contributing to each estimated parameter (5.1).

### 2.2. Robust Estimation of Individualized Neural Model Parameters

While theoretical neural mass models operate in continuous-time, fMRI experiments have limiting temporal sampling rates. Therefore, we approximate the continuous time neural model by fitting a discrete-time analogue for temporal resolution Δ*t* (e.g. the sampling TR; Fig. 1 B):

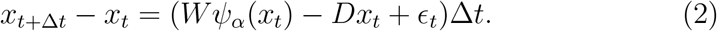

Parameter estimation in the MINDy algorithm contains three main ingredients, which ensure that estimates are robust, reliable, and valid. First, the transfer functions of neural mass models are allowed to vary by brain region through the scalar parameter *α*:

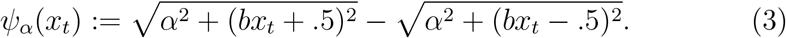

Each brain region has its own *α* parameter, fit on a subject-wise basis, while *b* is a fixed global hyperparameter (*b* = 20/3 for the current case). The use of a parameterized sigmoid allows for additional anatomical heterogeneity in region-wise dynamics. This form of transfer function is general enough to capture conventional choices (see SI Sec. 5.2 for a derivation of the function and its relation to conventional transfer functions). Secondly, we make use of recent advances in optimization to ensure that the fitting procedure (SI Fig. 9 C) is robust. By using Nesterov-Accelerated Adaptive Moment Estimation (NADAM, [34]) we achieve the speed advantage of stochastic gradient descent (SGD) algorithms, while at the same time preventing both over-fitting and under-fitting (see SI Sec. 5.3 for discussion). This approach leads to a very reasonable time duration for estimation (approximately one minute on a standard laptop; see SI Sec. 5.8 for a comparison with spDCM).

Lastly, we constrain the problem by decomposing the large matrix of connection weights (*W*) into two simultaneously fit components: a sparse component *W*_*S*_ and a low-dimensional component 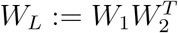 in which both *W*_1_ and *W*_2_ are *n × k* rectangular matrices with *n* being the number of neural masses (brain parcels) and *k < n* being a global constant that determines the maximum rank of *W*_*L*_. This decomposition is advantageous for concisely representing the interactions of structured networks and is the most important element of the fitting process. Sparseness criteria were achieved through *L*_1_ regularization ([35]) with the resultant fitting objective being to minimize:

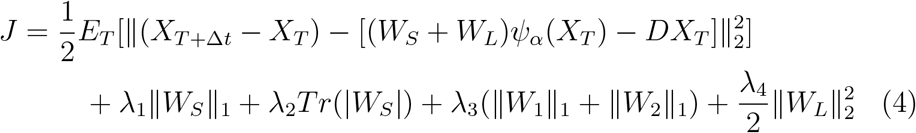

The notation *E*_*T*_ denotes the mean over all temporal samples considered (the “minibatch” of each iteration) so the first term simply corresponds to the mean square error of predictions. Each of the remaining penalty terms have a global regularization constant (*λ*_*i*_) that is shared across all subjects. This regularization scheme was adopted in order to reduce the dimensionality of the parameter estimation problem, while at the same time, attempting to reflect the consistently observed community-structure of brain connectivity measures. Under this view, brain connectivity patterns can be described in terms of communities (sub-networks) linked together by highly connected hubs. We envision the sparse component of connectivity to mimic the communication between connectivity hubs. By contrast the low-rank component is meant to account for the propagation of signals from hubs to their corresponding subnetworks and vice-versa.

We employ this two-component weight formulation as a heuristic that facilitates high-dimensional model fitting. In most analyses we only analyze the composite weight matrix than its components. However, preliminary results indicate that properties of this decomposition, namely the ratio of sparse vs. low-rank components, may be a marker of individual differences (see SI Sec. 5.6). Interestingly, recent work by Mastrogiuseppe and Ostojic ([36]) has also considered models in which connectivity is the sum of two terms: one low-rank and one random. The authors found that these structures produced low-dimensional dynamics which could be predicted based upon network structure and exogeneous (task) input. Such analyses may be relevent for understanding the role of connectivity in MINDy. Bayesian and algebraic interpretations of this penalty function are presented in SI Sec. 5.5. We also discuss the well-posedness of this problem (SI Sec. 5.5).

Throughout, we use the term “weights” to refer to the matrix *W* in estimated dynamic neural models. This is to differentiate the model connectivity parameter from the term “resting-state functional connectivity” (rsFC), which instead refers to the correlation matrix of BOLD time-series, rather than the mechanistic concept that it is often assumed to measure (i.e. direct and indirect interactions between brain regions). We reserve the term “effective connectivity” to indicate a causal, monotone relationship in activity between brain regions that evolves over no more than 2s (the typical fMRI sampling rate). Thus, both the fit model weights and the rsFC are ways to approximate the effective connectivity, even though rsFC may not support reverse inferences regarding directedness and causality.

### 2.3. Study Design

The objective of the current study was to rigorously validate a new approach for data-driven whole-brain modeling (MINDy). The study design consisted of both numerical simulations to validate the accuracy of models with respect to a known ground-truth, as well as empirical analyses of HCP resting-state data. The latter analyses were designed to test whether MINDy adds additional value in-practice and to quantify its performance in the presence of known experimental confounds (e.g. motion).

### 2.4. Empirical Dataset

#### 2.4.1. HCP Resting-State Scans

Data consisted of resting state scans from 53 subjects in the Human Connectome Project (HCP) young adult cohort, 900 subject release (for acquisition and minimal preprocessing details, see [37]; WU-Minn Consortium). Each subject underwent two scanning sessions on separate days. Each scan session included two 15-minute resting-state runs (two scans × two days) for a total resting state scan time of 60 minutes (4800 TRs). The two runs for each session corresponded to acquisitions that had left-right and right-left phase-encoding directions (i.e., balanced to account for potential asymmetries in signal loss and distortion). The TR was 720ms and scanning was performed at 3T. The subjects were selected by starting with an initial pool of the first 150 subjects and then excluding subjects who had at least one run in which more than 1/3 of frames were censored (i.e. 400 bad frames out of 1200).

Although this criterion greatly decreased the number of usable subjects from the initial pool of 150 to 53 (attrition=65%), it should be noted that it is likely to be overly conservative. We employed such a strongly conservative criterion for this first-stage validation effort to provide the cleanest data from which to test the model. Likewise, we had the luxury of drawing upon a very large-sample dataset. In contrast, we believe that the exclusion criteria will not need to be as conservative in a research setting for which model cross-validation is not performed on every subject (i.e., it is probably overly stringent to require that all four sessions be clean, since we only used two sessions at a time). In particular, the use of cross-validation required that two models be fit for every subject using disjoint data so that the validation required twice as much data as would normally be required. Moreover, we required that the data be uniformly clean so that we could parametrically vary the amount of data used (i.e. criteria were in terms of absolute cleanness for each scanning session rather than number of clean frames). However, there is no reason why the models could not be fit to clean segments of scanning sessions.

#### 2.4.2. Preprocessing

Data were preprocessed through the rsFC pipeline proposed by Siegel and colleagues ([28]; SI Fig. 9 A). The first stage of this pipeline is the HCP minimal pre-processing pipeline (see [37]) with FSL’s ICA-FIX correction ([38],[39]). We then applied one of three second-stage pipelines developed by Siegel and colleagues ([28]; Sec. 3.7.3), to test the effects of including various additional preprocessing steps. In all three pipelines, drift was mitigated by detrending data. The pipelines also all included motion scrubbing, using both Framewise Displacement (FD) and the temporal derivative of variation (DVARS). Frames that exceeded the cutoffs for FD (.2mm) or DVARS (5% above median) were replaced via linear interpolation ([40]). Respiratory artifact was mitigated with a 40^*th*^-order .06-.14 Hz band-stop filter applied to FD and DVARS for all pipelines ([28]).

The three second-stage pipeline variants differed however, in the number of regressors included to remove nuisance signals. The first variant mainly corrected frame-to-frame motion artifact, which has been found to induce systematic errors in functional connectivity studies, i.e. generating spurious short-distance correlations while diminishing long distance ones ([41]). In addition to data scrubbing, motion correction was performed using the 12 HCP motion regressors and their temporal derivatives. The second, more extensive pipeline variant, known as CompCor, also removed cardiac and respiratory signals, by additionally regressing out principal components of the white matter and cerebrospinal fluid signals ([42]). Lastly, the third pipeline variant also added global signal regression (GSR; [43]), in which the mean signals from white matter, cerebrospinal fluid, and grey matter are also included as regressors. As the variables included are cumulative, these three pipelines form a representative hierarchy of preprocessing approaches, that optionally includes CompCor or CompCor+GSR in addition to motion scrubbing. For most analyses we used the full (third) pipeline, but we also compared the effects of pipeline choice (Sec. 3.7.3).

After the second-stage preprocessing pipelines, we deconvolved the parcellated data (see below) with the generic SPM hemodynamic kernel ([30]) using the Wiener deconvolution ([29]). For the Weiner deconvolution, we used noise-power to signal-power parameter .02. The value of this parameter dictates the degree of temporal filtering during the deconvolution with smaller values being more parsimonious (less additional filtering). We then smoothed by convolving with the [.5 .5] kernel (2 point moving average) and z-scored the result. To test the robustness of the fitting procedure, we compared the effect of the second-stage preprocessing pipelines for some analyses. Based upon these results, we chose the third variant pipeline (GSR+CompCor+motion) for all other analyses. For all empirical rsFC analyses we use the deconvolved data to prevent bias from the deconvolution procedure in comparing MINDy and rsFC. As described further below, we also tested the effect of mismatches between “true” and canonical HRF models (Sec. 3.7.4, 3.7.5).

We defined derivatives in terms of finite differences. Since HCP employed unusually fast scanner TRs, we temporally downsampled the estimated derivatives for calculating goodness-of-fit in non-simulation analyses to represent the anticipated benefits to typical fMRI protocols and improve SNR: *dX*(*t*) = (*X*(*t* + 2) − *X*(*t*))/2.

#### 2.4.3. Parcellation Atlases

In the present framework we define whole-brain models in terms of connected neural populations. Thus, the approach demands that the neural populations be defined a-priori. For the present case of fMRI data, we define these populations to be anatomical brain regions corresponding to subcortical structures and cortical parcels. For subcortical regions, we follow the HCP protocol in considering 19 subcortical regions as defined by FreeSurfer ([44])). For cortical parcels, we generally employed the gradient-weighted Markov Random Field (gwMRF) parcellation with 200 parcels per hemisphere ([21]) and organized according to the 17 cortical networks described in [45]. The gwMRF parcellation is optimized to align with both resting-state and task fMRI, and has been found to demonstrate improved homogeneity within parcels relative to alternative parcellation techniques. However, for anatomical analyses we compared with an additional atlases (SI Fig. 11 C,G) to ensure generality : the MMP atlas ([20]) which was also derived from a combination of rest and task-based data. The MMP (Multi-Modal Parcellation) atlas is symmetric with 180 parcels per hemisphere.

### 2.5. MINDy Fitting Procedure

MINDy models were fit by applying the iterative NADAM algorithm ([34]) to optimize the MINDy cost-function (Eq. 4; see SI Sec. 5.12). This algorithm belongs to the family of stochastic gradient-descent techniques and we provide further detail/discussion regarding NADAM in SI Sec. 5.3. To ensure algorithmic stability, we used two transformations (one each for the curvature and decay parameters) which are detailed in SI Sec. 5.12. The gradient equations for each parameter in detailed in SI Tab. 11.

#### 2.5.1. Compensating for Regularization Bias

In order to retrieve parsimonious weight matrices and reduce overfitting, we employed regularization to each weight matrix (both the sparse and the low-rank matrices) during the fitting process. One consequence of regularization, however, is that the fitted weights may be unnecessarily small as weight magnitudes are penalized. After fitting, we therefore performed a global rescaling of weight and decay contributions for each model using robust regression ([46]) as implemented by MATLAB2018a. Specifically, we fit two scalar parameters: *p*_*W*_, *p*_*D*_ in regressing *dX*(*t*) = *p*_*W*_ *Wψ*(*X*(*t*)) − *p*_*D*_*Dx* collapsed across all parcels. Here *p*_*W*_ and *p*_*D*_ represent global rescaling coefficients for the weights and decay, respectively. As this compensating step only used global rescaling for *W* and *D*, it had no effect upon the relative values for each parcel, only the total magnitude of the *W* and *D* components. Since only two values are estimated, this step does not reintroduce overfitting. Although we performed this step using robust regression, we obtained identical results using conventional linear regression. The choice of robust regression was made as a safeguard for high leverage points as might occur due to motion artifact. However, results indicate that conventional regression may suffice for sufficiently clean data.

#### 2.5.2. Selecting Hyperparameters and Initialization

The proposed fitting procedure requires two sets of hyperparameters: the four regularization terms specific to our procedure and the four NADAM parameters ([34]). By “hyperparameters” we refer to free constants within an algorithm which distinguishes them from the “parameters” of an individualized model.

Hyperparameters were hand-selected for model goodness-of-fit and reliability, based upon prior numerical exploration with a subset of 10 subjects who did not belong to the “data source” subjects. Thus, these subjects were not included in any further analyses so the hyperparameter selection procedure did not artificially inflate model performance. The selection criteria were to maximize cross-validated goodness-of-fit under the constraint that test-retest correlations were greater than .7 for all parameters. Regularization values were sampled with resolution .005. The chosen set of hyperparameters was then constant for all test subjects. Hyperparameter values and discussion are included in SI (Tables 12, 13). The initialization distributions for the algorithm were similarly selected using the same subjects and are included in the SI (Table 12). We explored the effect of hyperparameter choices on the sparsity of MINDy relative to rsFC and found that for any choice of regularization hyperparameter (even 0), the group-average MINDy weights are sparser than rsFC (SI Sec. 5.7).

### 2.6. Ground-Truth Simulations

#### 2.6.1. Realistic Whole-Brain Simulations

For the analyses of sensitivity and individual differences we generated new, synthetic individuals by randomly sampling neural mass model parameters from the parameter distributions estimated from the full dataset (i.e. N=53 participants). The decay and curvature parameters (*α, D*) were independently sampled for each parcel from that parcel’s population distribution. The weight matrices, however, were sampled as a whole rather than sampling each individual connection as we found that the latter approach led to pathological behavior in simulations. For the robustness analyses, ground truth models were drawn from those fit to experimental sessions. The ground-truth models were simulated as stochastic differential equations (*dX* = *f*(*X*)*dt* + *σ*_*W*_ *dW*_*t*_) with *f*(*X*) the deterministic neural mass model and units time measured in terms of the fMRI TR. Models were Euler-Maruyama integrated with *dt* = 1/4 and *σ*_*W*_ = .45 in units TR (720*ms*) to generate simulated neural activity time-series. Neural-activity was then downsampled to 1 TR resolution (as opposed to the simulation’s time-step of dt × TR) and convolved with the SPM-style HRF kernel ([30]; SI Fig. 20 C):

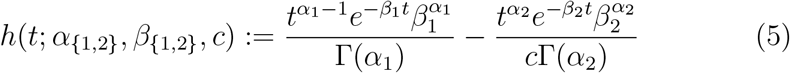

Here Γ is the gamma function (equal to factorial for integer values). The parameters describe two gamma-distributions (one *α, β* pair per distribution) and a mixing coefficient (*c*) to generate a double-gamma distribution. Parameters were set to their default values (*α*_1_ = 6, *α*_2_ = 16, *β*_1_ = 1, *β*_2_ = 1, *c* = 1/6) except for the simulation featuring HRF variability. In this case, random perturbations were added to each parameter and were drawn from the normal distribution with mean zero and SD as indicated. The final simulated BOLD signal was then generated by adding white, gaussian noise with the indicated SD (Fig. 1 D).

#### 2.6.2. Randomized Network Simulations

Although some ground-truth simulations leveraged the empirical MINDy distributions to maximize realism (Sec. 3.2.1, 3.2.2,3.7.1,3.7.4), others used randomly generated networks of Hopfield or neural mass models (Sec. 3.7.5, 3.8, SI Sec. 5.8). The latter ground-truth simulations prevent circularity (i.e. using MINDy distributions to test MINDy) by drawing parameters from random hyperdistributions independent of previous analyses. These distributions were designed to possess complex network structures by superimposing three simpler network structures: community-structure (*M*_1_), sparse structure (*M*_2_), and low rank structure (*M*_3_). These distributions are characterized by standard-deviation parameters *σ*_1_ and *σ*_2_. An asymmetry parameter *σ*_*a*_ characterizes the degree to which the resultant network is asymmetric. Each standard-deviation parameters was randomly sampled for each ground-truth model from normal distributions: *σ*_1_, *σ*_*a*_ ~ *N* (4, .05^2^) and *σ*_2_ ~ *N* (3, .05^2^). Connectivity matrices were then randomly parameterized as follows:

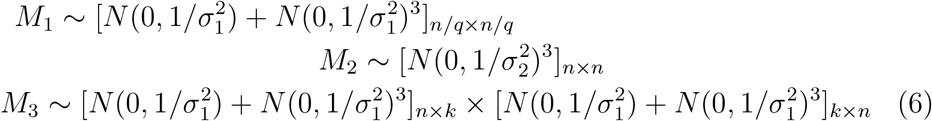

Here, the bracket outside each matrix denotes its size with *n* = 40 denoting the total number of nodes, *q* denoting the number of nodes per community (randomly set to either 1 or 2 with equal probability), and *k* = 5 denoting the rank of the low-rank component. We denote the Kronecker product ⊗ and use it to copy the community level matrix (*M*_1_) among each node belonging to the community: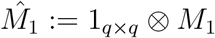. The three component matrices are then combined as follows:

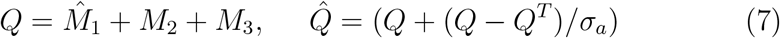

The final matrix *C* is formed by censoring elements of 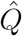 whose absolute value is below 1/4 the standard deviation of 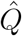. This same technique was used to randomly generate networks of Hopfield models with homogeneous, heterogeneous, or nonlinear hemodynamic effects and realistically-paramaterized neural mass models with nonlinear hemodynamics.

#### 2.6.3. Hopfield Network Simulations

We employed two cases of non-MINDy ground truths: Hopfield networks and neural-mass models (Sec. 3.7.5,3.8). Continuous, asymmetric Hopfield models are similar in form to the MINDy model, but use a tanh transfer function:

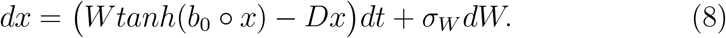

Here, the slope vector 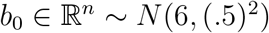 and diagonal elements of the decay matrix *D* drawn from *N* (.4, (.1)^2^) (non-diagonal elements are zero). As elsewhere, the symbol denotes the Hadamard product (element-wise multiplication). Models were simulated via Euler-Maruyamma integration with dt=.1s, *σ*_*W*_ =.2, TR=.7s, and total simulation length t=10,000. We considered the case in which no hemodynamics are present, in which case MINDy is fed *x*(*t*) downsampled according to TR, and the case in which *x*(*t*) is convolved with spatially heterogeneous hemodynamics and deconvolved with the canonical HRF before being fit by MINDy. In the latter case, the HRF function was parameterized as before, but with the ground-truth *α*_1_ parameter for each brain region drawn from *N* (6, (.25)^2^) and the *β*_1_ parameter drawn from *N* (1, (.25/6)^2^). The simulated BOLD was produced by convolving the simulated time-series with the ground-truth HRF before temporal downsampling. In both cases, initial conditions for each node were independently drawn from *N* (0, 1) and the first 100 samples were dropped. Since the total number of nodes was approximately one-tenth of those used in the HCP data, we rescaled the dimension of the low-rank component by one-tenth (from 150 to 15). Similarly, we rescaled the regularization terms inversely proportio√nate to the effect of rescaling *W* by a factor of 10: (*λ*_1_, *λ*_3_ by 1/10, *λ*_2_ by 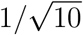 and *λ*_4_ by 1/10^2^). For simulations using the Balloon-Windkessel model of hemodynamics, *x*(*t*) was rescaled to the range of average synaptic gating via 5*S*(*t*) = 1 + *tanh*(*x*(*t*)/10). This transformation of *x* was then substituted into the nonlinear hemodynamic model (below) to generate simulated BOLD signal. In all cases, time series were z-scored, smoothed via nearest-neighbor ([.5 .5] kernel) and run through MINDy for 150,000 iterations (approximately 70 seconds) with the original batch size of 250.

#### 2.6.4. Neural Mass and Windkessel-Balloon Model Simulations

For our neural mass ground-truth simulations (Sec. 3.7.5, 3.8), we largely followed the approach of Wang and colleagues ([13]) in using single-population neural mass models (20 masses/simulation in Sec. 3.7.5 and 6 to 16 in Sec. 5.8) with Windkessel-Balloon model hemodynamics ([47])). Similar to the MINDy model, the neural mass model ([48]) contains a monotone nonlinearity 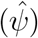 and linear decay 1/*τ*_*S*_:

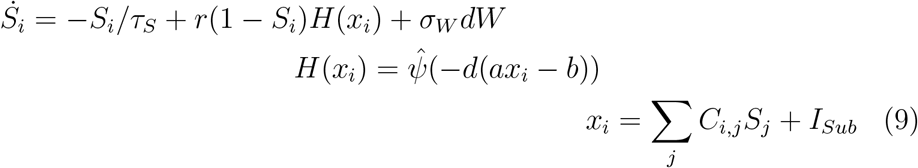

The variable *S* describes the average synaptic gating, while *H* describes the population firing-rate. We used the default parameter settings: *τ*_*S*_ = .1*s*, *a* = 270*n/C*, *b* = 108*Hz*, *d* = .154*s*, *r* = .641. Unlike Wang and colleagues ([13]), we used a logistic sigmoid transfer function for 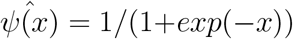 instead of the rectified linear transfer function: *x/*1−*exp*(−*x*), as the former is less prone to pathological behavior in random networks. Subcortical input was *I*_*sub*_ = 5. Connection weight matrices were randomly generated as described in the previous section, but with 1.5 added to all recurrent connections and the resultant matrix scaled by a factor of 100. Simulated neural activity is converted into BOLD signal through the Windkessel-Balloon model ([47]):

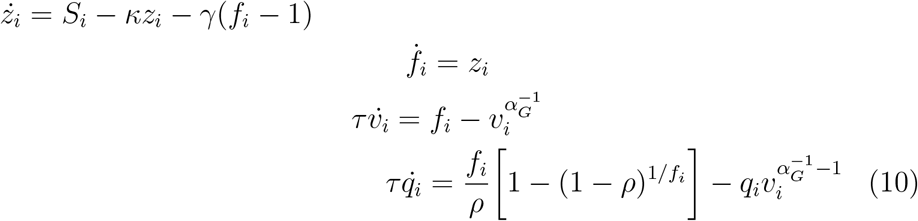

The variables *z*,*f*,*v*, and *q* model vasodilation, inflow, blood volume, and deoxyhemoglobin content, respectively. Parameters were: *ρ* = .34, *κ* = .65*s*^−1^, *γ* = .41*s*^−1^, *τ* = .98*s, α*_*G*_ = .32. The simulated BOLD signal at each TR is then modeled as:

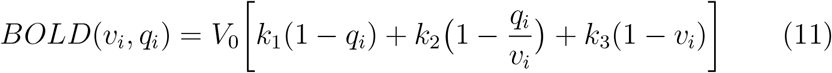

Resting blood volume fraction is denoted *V*_0_ = .02. Scanning parameters *k*1, *k*2, *k*3 were set to 3T values according to Demirtas and colleagues ([12]): *k*_1_ = 3.72, *k*_2_ = .53*k*_3_ = .53. Simulations were run with dt=25ms and *σ*_*W*_ = .005 for total length *t* = 40, 000. Sampling was performed every 29 time-steps (*TR* = 725*ms*) and the first 10% of samples were dropped. The resulting time-series were deconvolved with the canonical HRF assumed by MINDy and z-scored. MINDy hyperparameters were identical to the rate-model case and MINDy was run for 10,000 iterations (approximately 6 seconds) with batch size 250. Initial conditions for hemodynamic variables were randomly sampled from |*N* (0, 1)|. Initial conditions for the neural variable (*S*) were generated by first sampling *S*_0_ ∼ |*N* (0, 1)| and then performing the transformation *S*_0_/(1 + *S*_0_).

### 2.7. Simulations for DFC analysis

For analyses of dynamic functional connectivity, models were estimated for each subject (one per session) using the full HCP temporal resolution *dX*(*t*) = *X*(*t*+1) − *X*(*t*). These models were then used to generate simulated resting-state fMRI data, but with additional process noise added as would be expected in observed fMRI timeseries data. We used the same time-scale for simulation as in the validation models (dt=.5 TR). However, whereas the validation simulations employed process noise containing constant variance across parcels, we used a naive estimate of process noise for each parcel, that was based upon the residual error of model fits over subsequent time-steps. We avoided doing this in the validation stage so that ground-truth parameters could not be recovered simply by observing noise. The residual error covaried with the decay parameter across parcels at the group-level, but not at the individual level, despite individual differences in both noise and decay being reliable within parcel. We reintroduced parcel-based variation into the DFC simulations to obtain maximum realism. We considered both the case in which process noise was allowed to vary by parcel but not by individual within a test-retest group (e.g. using the mean noise across subjects for each session separately), as well as the case in which process noise was determined on a subject-wise basis. Results obtained with either method were near-identical for the DFC reliability analyses so we present results using the session-wise group-mean process noise (e.g. the mean process noise for each parcel averaged across all day 1 scans or all day 2 scans). Initial conditions were drawn from each subject’s observed data for that scanning session. Simulations were run for 2600 time steps (1300 TRs) using 15 different initial conditions per session and temporally downsampled back to the scanning TR. After simulation, we downsampled from the 400 parcel to the 100 cortical parcel variants of gwMRF ([21]) and removed subcortical ROIs in order to reduce computational complexity of subsequent DFC analyses.

### 2.8. DFC Analyses

DFC analyses consisted of the standard deviation and excursion ([49]) of the time-varying correlation between brain regions. To calculate time-varying correlations we used Dynamic Conditional Correlation (DCC; [50]). To avoid confusion with other references to “standard-deviation” we refer to this measure as “*σ*-DFC” as it pertains to time-varying correlations. Formally, *σ*-DFC is calculated by first estimating the time-varying covariance using DCC. Under this approach, the data, (*y*_*t*_) is modeled as a zero-mean stochastic process with auto-regressive covariance:

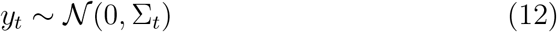

with time-varying covariance matrix Σ evolving according to the first-order autoregressive model:

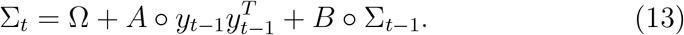

The matrices Ω, *A, B* are estimated in DCC using maximum-likelihood. We define the *σ*-DFC matrix as the standard deviation (over time) of the time-varying correlation matrix *Q*_*t*_:

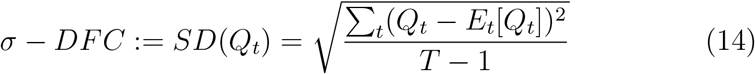

with *E*_*t*_[*Q*_*t*_] denoting the sample mean over time. To ensure numerical stability, we repeated the DCC algorithm 10 times per case (simulation or true data) and used the median estimated time-series for time-varying correlations. The excursion measure was calculated according to ([49]). Reliability was computed for each pair of region’s DFC statistics using Fisher’s ICC of group-demeaned DFC metrics between scanning session (ICC(2,1) in the Shrout and Fleiss convention [51]). Overall reliabilities collapsed across all regions were calculated using Image Intraclass Correlation ([52]).

### 2.9. Sensitivity Analyses

We conducted sensitivity analyses in Sec. 3.2.1 to test how the different mechanisms of ground-truth models (e.g. connections vs. decay) influence the estimates of “connectivity” in MINDy and rsFC. We were particularly interested in how each method responded to local heterogeneity (i.e. are MINDy/rsFC estimates of connection strength sensitive to local model parameters: decay and curvature). For each batch of the sensitivity analyses, we first simulated a resampled individual multiple times to generate a distribution of trial-to-trial variability (“within-subject”) in elements of MINDy’s weight matrix and the rsFC matrix. We then held the weights of the ground-truth neural mass model constant while resampling either the curvature (*α*) or decay (*D*) parameters and calculating MINDy weights and rsFC from simulations of the new model. Changes in the estimated connectivity (weights or rsFC) were deemed significant if they occurred with *p* < .05 for the corresponding “within-subject” distribution.

### 2.10. Statistical Analyses

Statistical testing was primarily within-subject between method/condition (e.g. paired t-tests). We used the conservative Bonferroni method for all multiple-comparison corrections. All reported p-values are calculated for two-tailed tests unless indicated otherwise. We use *p* ≈ 0 to denote *p*-values calculated as less than 10^−20^ for which precise numerical estimates may deteriorate.

## 3. Results

### 3.1. Overview of Results/Approach

The Results of the paper are structured as follows. The first section serves to relate MINDy parameter estimates to resting-state Functional Connectivity (and related partial correlation approaches) in terms of differentiating/identifying sources of individual variation. The “ground-truth” models for validation in this first set of analyses are drawn from the empirical distribution of MINDy parameters to ensure that the resultant simulated data is realistic. The second section directly addresses the potential for overfitting by testing whether MINDy models cross-validate and whether parameters are reliable. The third section demonstrates that MINDy parameters have distinct anatomical gradients consistent with previous, theoretical results ([12],[13]), and highly conserved individual variation (a feature not present in over-fit models). The fourth section demonstrates models’ predictive validity by reproducing individual differences in resting-state dynamics using the empirical models. In the fifth section, we demonstrate that the approach is robust to measurement noise, preprocessing pipelines, and hemodynamic confounds. This section uses three forms of “ground-truth” models. For initially testing robustness to noise and global hemodynamic variability, we again use parameters drawn from the empirical distribution to ensure maximum realism. In subsequent analyses, however, “ground-truth” parameter values are drawn from random hyper-distributions independent of the data and combined with more nuanced hemodynamics. This step tests model performance with more exotic “ground-truths” and prevents circularity. We also consider an additional case in which the simulated fMRI data is generated from randomly-parameterized neural-mass models (operating at the millisecond-scale) to provide insight into the relationship/limitations of MINDy parameter estimates from fMRI and the underlying synaptic connectivity. In the sixth section (Sec. 3.8), we summarize comparisons with Dynamic Causal Modeling which receive fuller treatment in the SI (Sec. 5.8). The final results section directly assesses data-requirements of MINDy and provides a minimum data quantity (>15 minutes) to prevent over-fitting.

### 3.2. MINDy Retrieves Individual Differences

#### 3.2.1. MINDy Retrieves Individualized Connectivity

A key goal of our investigation was to determine whether MINDy was sufficiently sensitive to reveal individual differences in connectivity weights that have become the focus of recent efforts within the rsFC literature ([53], [54]). We tested the model by reconstructing individual differences in connectivity weights of simulated subjects and comparing them against both classical rsFC and the partial correlation matrix. Simulated subjects were generated by permuting MINDy parameter sets across individuals (see methods). We then simulated the resultant model with process noise and hemodynamics to generate realistic BOLD fMRI time series (see methods; Fig. 1 C; SI Fig. 9 B). This provided a ground-truth set of simulated fMRI data, from which we could compute the rsFC/partial correlation matrices for each “subject”, and also determine the fidelity of recovered parameters (i.e., compared against true parameters used to generate the simulated data). To assess the performance of the model estimation procedure, we considered two metrics: the validity of estimated connectivity weight differences between subjects (Fig. 2 B) and the sensitivity of each procedure to different model components (SI Fig. 17 A). These sensitivity analyses reveal whether each approach (rsFC matrix, partial correlation matrix, or model estimation) misclassifies variation in some other model component (e.g. decay rates) as being due to a change in weights. To better assess sensitivity, we generated data after varying only one model component at a time across the simulated subjects: the weight matrix (*W*), transfer functions (*α*) or decay rates (*D*).

**Fig 2.**
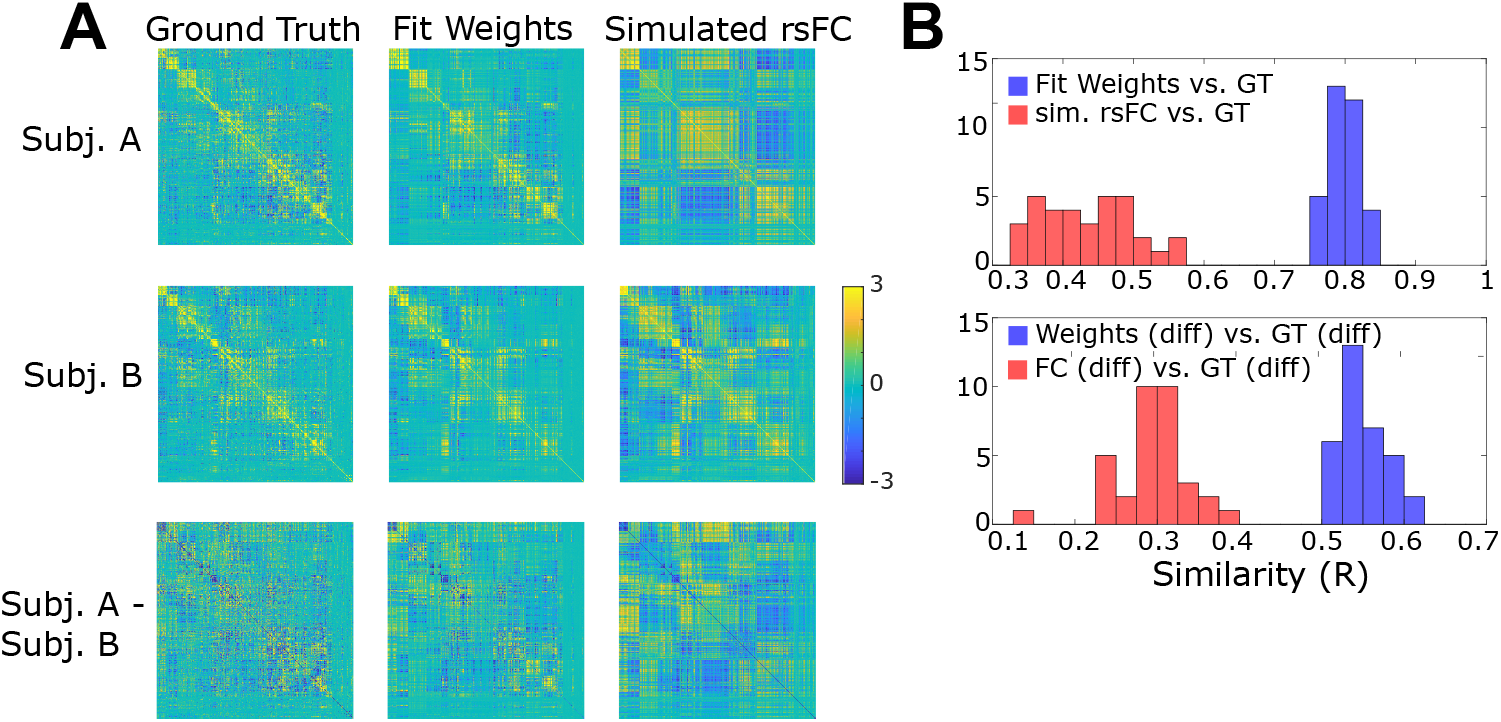
Ground-truth validation of MINDy and rsFC at the level of single-subject and inter-subject variation. A) First column: Example ground-truth weight matrices for two simulated subjects (top two rows) and the difference between ground-truth weights (bottom-row). Second column: Recovered weight matrices using MINDy for both subjects and their difference. Third column: same as second but using the rsFC. Fit weight matrices and simulated FC matrices are shown in standard-deviation (SD) units with SD computed across the offdiagonal elements of each individual matrix. The ground-truth matrices are displayed in units 2/3×SD to aid visual comparison. B) Top row: histogram of performance at the simulated single-subject level (correlation with ground-truth [GT]) for MINDy (blue) and rsFC (red). Bottom row: same as top but for for predicting the differences in matched-pairs of simulation subjects who differed only in ground-truth connectivity. Simulation subjects were generated by sampling from the distribution of empirical (HCP) MINDy parameters (see Sec. 2.6.1).

Results indicated that MINDy was able to accurately recover the ground-truth weight matrix for each individual (Fig. 2 A,B). Thus, the simulated weight changes that differentiated one individual from another were recovered well by the MINDy parameter estimation approach. Moreover, MINDy weight estimates were found to significantly outperform rsFC and partial correlation measures (computed on the simulated timeseries data) in their ability to accurately recover both the ground-truth connectivity matrix of simulated individual subjects, as well as the differences between individuals (Fig. 2 B; SI Table 5). This finding suggests that the modest relation between rsFC and ground-truth connectivity weights is primarily driven by the group-average connectivity as opposed to individual differences. However, rsFC may be disadvantaged in this comparison as it does not typically permit sparseness commensurate with empirical MINDy weights (Fig. 5 A,B). Therefore, we used partial correlations as an additional benchmark. While partial correlations quantitatively improved upon rsFC estimates (single-subject: *R* = .537 ± .032, inter-subject: *R* = .392 ± .027), performance remained significantly lower than MINDy (single-subject: *paired* − *t*(33) = 40.51, *p* ≈ 0, inter-subject: *paired* − *t*(33) = 23.62, *p* ≈ 0).

The above analyses were designed to illustrate the additional utility of MINDy in empiricial contexts over the most common current approaches (rsFC and partial correlation). For this reason, we generated ground-truths from the empirical distributions to ensure maximal realism. In later analyses (Sec. 3.8), we compare MINDy to a much closer modeling approach (Spectral DCM; [7]). We reserve these comparisons for later as they employ a very different approach to generating ground-truth models: seeking to minimize bias and sample over a wide range of potential ground-truth scenarios. The anatomically-detailed models used in the current section are also too large for Spectral DCM to estimate using available computational resources (Sec. 3.8).

#### 3.2.2. MINDy Disentangles Sources of Individual Differences

After we established that MINDy outperforms rsFC and partial correlations in retrieving true individual differences in weights, we benchmarked the sensitivity of each approach to other sources of individual variation. Rather than measuring how well each procedure correctly retrieves connectivity, these tests quantify how well each approach selectively measures connectivity as opposed to other sources of variation (see methods). We quantified sensitivity in terms of how often MINDy and rsFC reported that a connection changed in strength between simulated models, when in reality only the curvature or decay terms were altered (SI Fig. 17 A). Results indicate that MINDy correctly detects the sources of individual variation when due to local changes such as decay rate and transfer function shape, as these have no appreciable impact on MINDy’s connectivity estimates (the false positive rate is near that expected by chance). By contrast, rsFC measurements were highly sensitive to the decay rate (27.5 ± 12% of connections changed vs. 7.6 ± .6% for MINDy, with 5% expected by chance), indicating that some individual differences in FC may be reflective of purely local brain differences as opposed to connectivity between brain regions (SI Fig. 17 A; SI Table 6). These results indicate that MINDy promises to improve both the mechanistic sensitivity and the anatomical accuracy of inferences based upon individual differences in resting-state fMRI. However, it is still the case that resting-state fMRI exhibits *generalized* sensitivity to individual differences in neurobiology, which may suffice for some applications, such as biomarker discovery (see Sec. 3.4).

### 3.3. MINDy Parameters are Reliable

In addition to determining the validity of MINDy parameters, it is also critical to establish their reliability. We examined this question by analyzing measures of test-retest reliability of the parameter estimates obtained for human subjects contributing resting-state scans on two separate days (30 minutes each). Results indicated that MINDy had high test-retest reliability for all parameter estimates (> .75; Fig. 3 A). The reliability of weight estimates was significantly higher than rsFC reliability, although the mean difference was modest (Δ*R* .045, SI Table 7, SI Table 8). By contrast, the variability in reliability was noticeably smaller for MINDy, meaning that while the mean advantage of MINDy in terms of reliability was modest, its performance was much more consistent across subjects (less variable reliability; SI Table 8).

**Fig 3.**
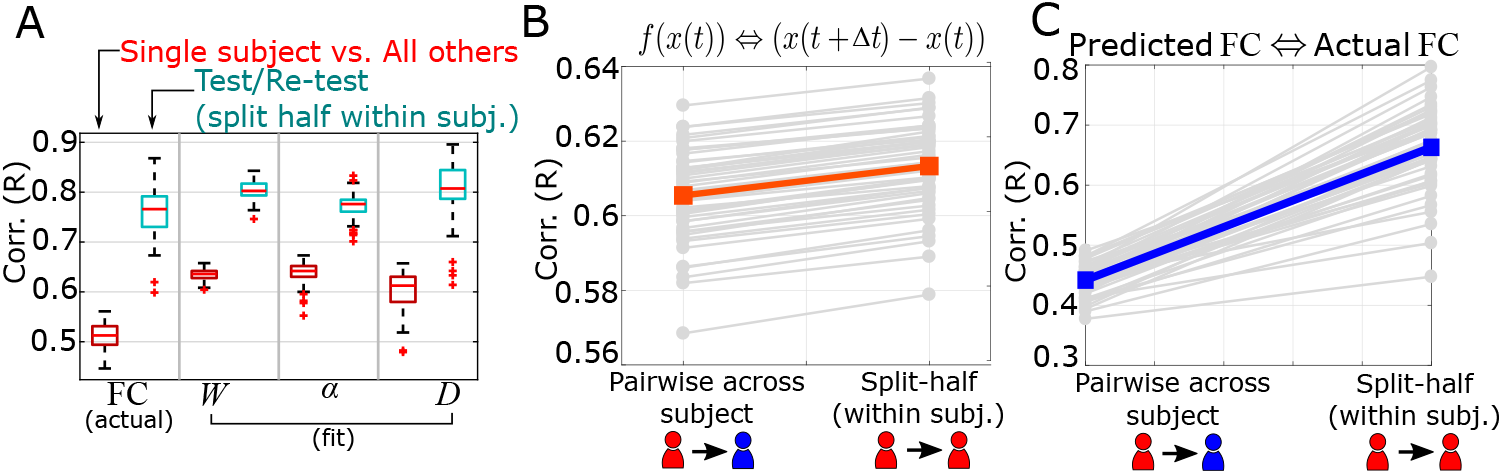
MINDy parameters and predictions are personalized and reliable. A) Comparison of the test-retest similarity between subjects (red) and the test-retest reliability (blue) for rsFC and the MINDy parameters. B) Goodness-of-fit for a single time-step prediction is uniformly (but minutely) greater for comparing test-retest predictions within a subject vs. between subjects. Performance is in terms of predicting the difference time series. Red line indicates group-mean C) This relationship magnifies across time steps as evidenced by far greater similarity in test-retest predicted FC from model simulations of the same subject vs. different subject. Performance is in terms of predicting the empirical rsFC on a different scanning session. For similarity to the same or both scanning sessions see SI Fig. 16. Blue line indicates mean.

### 3.4. MINDy Parameters are Personalized

For sake of comparison with FC we have thus far emphasized the ability of MINDy to extract brain connectivity. However, MINDy fits brain models, with the connectivity weights (Fig. 5A,B) comprising just one component. For the approach to faithfully reflect the stable differences among individual brains, it is important that it not just accurately estimates the neural parameters that describe human brains, but that these parameters accurately capture individual differences and predict brain activity. Using the “connectome fingerprinting” approach ([55]), we compared whether MINDy parameter estimates and the combined model uniquely identify individuals within a sample. This analysis was conducted in two ways. First, we computed separate parameter estimates for each individual in each testing day session. Then we examined whether the parameters estimated from one day showed the highest similarity to the same individual on the other day (relative to all other individuals in the dataset; Fig. 3 A). Secondly, we used the estimated model from one day to test whether the estimated parameters provided the best fit to the fMRI data timeseries recorded on the second day, again relative to the estimated parameters from other subjects. Specifically, this second analysis provides a strong form of cross-validation testing and we performed it for both predictions of the empirical timeseries (Fig. 3 B) and for predictions of each subject’s empirical rsFC, both cross-validated across sessions (Fig. 3 C). In all analyses, we found that the best predicting model for every subject was almost always their previously fit model (Table 7). In particular, we achieved 100% accuracy when conducting connectome fingerprinting based on MINDy weight parameters (SI Fig. 17 B), and when computing cross-validated goodness of fit/cross-validated predicted rsFC (Fig. 3 B,C). For pairwise analyses of subjects, see SI Fig. (17 F).

Similar patterns emerged but also some important differences, when conducting parallel analyses using rsFC. Replicating prior findings ([55]), 100% accuracy was also achieved in connectome fingerprinting (SI Fig. 17E). However, between-subject similarity was significantly lower in the rsFC analysis. Conversely, in rsFC the distinction between across-sessions within-individual similarity scores (i.e. test-retest similarity) and the average similarity obtained between subjects was greater than that observed in the MINDy model weights (SI Table 7). These results suggest that rsFC may actually generate an exaggerated picture of the idiosyncratic nature of connectivity, since MINDy individual differences are partitioned not only into weights, but also into other mechanistic parameters that are attributed locally, to the node/parcel (i.e., the decay [*D*] and curvature [*α*] parameters). In other words, MINDy may provide a richer and more variegated perspective on the nature of individuality, than what can be obtained with rsFC which lumps together what may be multiple dimensions of individual difference, into a simple, undifferentiated measure. For applications such as biomarker discovery, these properties may not be relevant in that the apparent magnification of individual differences in rsFC over MINDy weights could prove beneficial despite the mechanistic ambiguity of rsFC. However, we also note that MINDy provides additional parameters (curvature and decay) which may also prove useful for biomarker discovery. Lastly, the relevant dimensions for biomarker discovery are in terms of separating phenotypes, rather than separating all individuals. Since MINDy can robustly separate individuals, it has the potential to influence biomarker discovery, but whether it possesses quantitative advantages over rsFC will need to be investigated in the context of explicit biomarker questions (and may be phenotype-specific).

### 3.5. Novel MINDy Parameters show reliable individual and anatomic variation

Interestingly, we observed important additional functional utility from examining the novel MINDy parameters that are unavailable in standard rsFC. With regard to individual variation and fingerprinting analyses, we found that even ignoring the weights completely, the transfer function curvature parameter (*α*) associated with each node showed high consistency across sessions within an individual, and also unique patterns across individuals, such that 100% accuracy could also be achieved in fingerprinting analyses (Fig. 3). A slightly lower accuracy (94.3%) was observed when using the MINDy decay (*D*) parameters, though even here performance was still significantly above chance (1.89%) in identifying individuals (Fig. 3 A; Table 7). Pair-wise, between-subject, comparisons of similarity in these parameters are reported in SI Fig. (17 B-E).

We followed-up on the identification of reliable individual differences through MINDy, by conducting exploratory analyses to examine which brain regions/connections exhibited the greatest inter-individual variability (SI Sec. 5.9). We found that the curvature parameter had greatest relative variability in prefrontal cortex, particularly inferior frontal gyrus (SI Fig. 13A), while the decay parameter had high variability in visual regions, the “hand” portion of post-central gyrus, and medial prefrontal cortex (SI Fig. 13B). Connections within the visual networks had the lowest individual variability while connections to/from the Temporal-Parietal network had the greatest (SI Fig. 13C, D). Although these initial findings are intriguing, due to sample size/bias considerations and the exploratory nature of these analyses, we view them as a launching pad for future insights rather than basic neuroscientific results per se (see SI Sec. 5.9).

Although the above analyses focused on individual differences in the unique MINDy parameters, these parameters also exhibited common patterns across individuals (SI Fig. 11 B,F) that revealed interesting anatomical structure and gradients (Fig. 4 A-C; SI Fig. 11 B,F). These may reflect regional variation in intrinsic dynamics (*D*) and efferent signaling (*α*) that vary across brain networks (SI Fig. 11 A,E), but also exhibit consistency even at the finer within-network scale (Fig. 4 A,B; SI Fig. 11 B,F). For example, most nodes within the Temporal-Parietal network showed high curvature, but also low decay parameters; in contrast, in nodes of the Control (A) network, the curvature parameter tended to be low, whereas the decay parameter was high. Group-mean values show the same anatomical gradient across the gwMRF ([21]; SI Fig. 11 D,H) and MMP ([20]; SI Fig. 11 C,G) atlases. It is important to note that the decay parameter only describes temporal integration at time scales commensurate with fMRI sampling. Thus, the decay parameter should not be conflated with the time-constant of traditional neural mass models just as the latter is distinct from the membrane time constant of neuronal models. Interestingly, the decay parameter in MINDy appears to reflect components of both temporally-extended signal integration and the time-constant of local sub-second integration. Whereas the mean value of the decay parameter correlates with absolute global brain connectivity (i.e. the sum of absolute values along a row of the rsFC matrix; *r*(377) = .911, *p* ≈ 0) the principal dimensions of individual variation (Fig. 4 C, SI Fig. 11 I,J) recreate the hierarchical organization of primate cortex as derived from the T1/T2 ratio map (*r*(358) = .583, *p* ≈ 0; using the MMP Hierarchy map by Demirtas and colleagues [12]). As a caveat, it is worth noting that these statistics do not take into account spatial autocorrelation (which is challenging to model, given the large and irregular shape of parcels), which could have contributed in part to the anatomical gradients we observed. This hierarchy has been the subject of recent studies into its relationship with local excitation/inhibition ([13], [12]) which is one physiological mechanism we suspect underlies the decay construct (see SI 5.1). This hierarchy also predicts the time-scales of local microcircuits, patterns of gene-expression, mylein density, and function (sensory-processing hierarchy; see [13], [12]).

**Fig 4.**
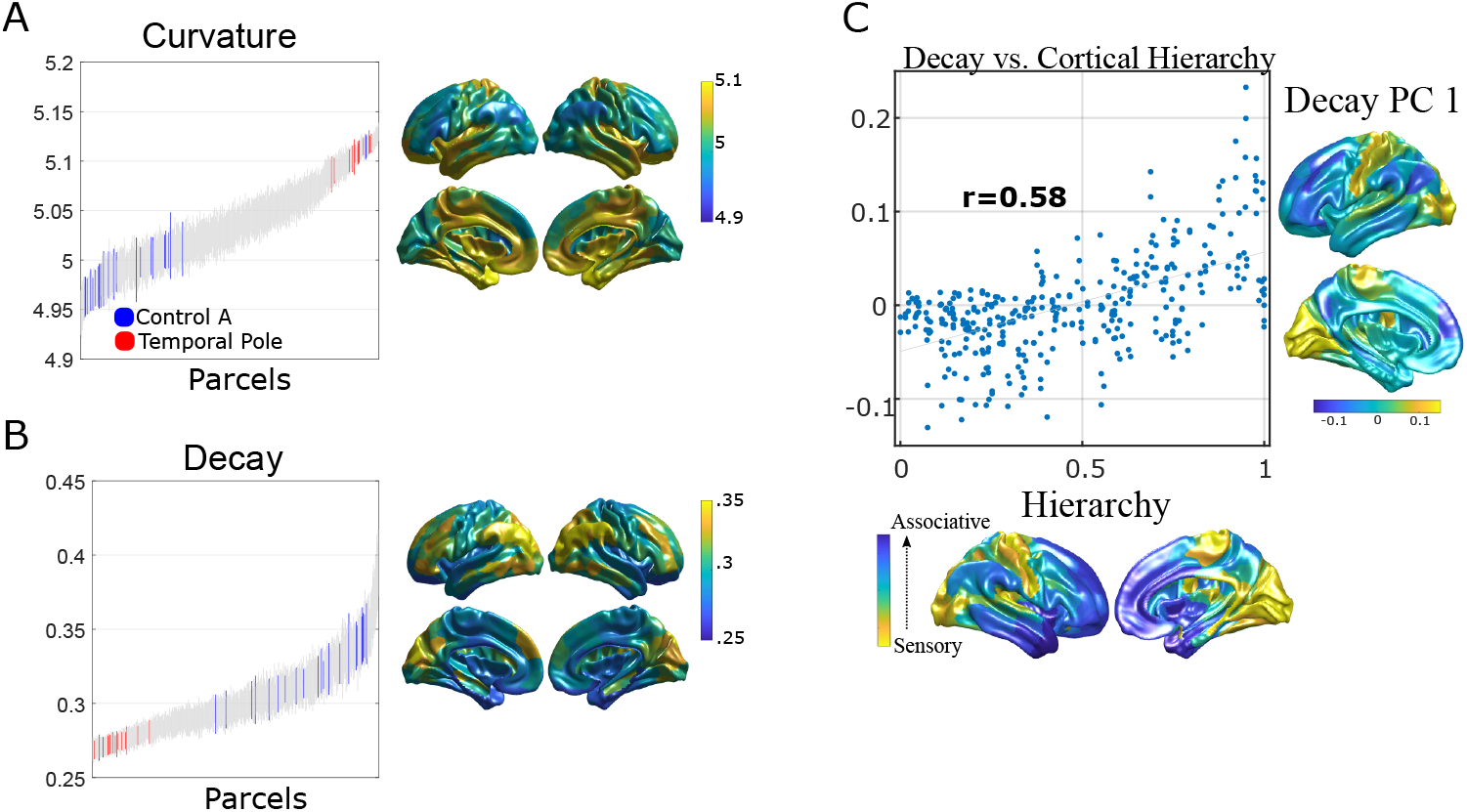
Local MINDy parameters display consistent anatomical distributions. A) The curvature-parameter displays network structure and is consistent across subjects at the finer parcel level. Parcels are ordered from least to greatest value for the curvature parameter (*α*) averaged across subjects and scanning sessions. Surface plots are for mean value. Two representative brain networks are highlighted (Control-A in red and Limbic-(Temporal Pole) in blue) to illustrate anatomical gradients in this parameter. B) Same as A but for the decay parameter (*D*). C) Correlation between the first principal component of MINDy decay and “hierarchical heterogeneity” provided by Demirtas and colleagues ([12]) based upon erf transform of the T1/T2 ratio (MMP parcelation). This measure has been theorized to reflect a hierarchy of cognitive abstraction from sensory to associative cortices.

In addition to the curvature and decay parameters, MINDy also differentiates from rsFC in the structure of the weight matrix (*W*) / connectivity matrix, both in terms of asymmetry (Fig. 5A,C) and sparseness (Fig. 5A,B). The former is a direct consequence of the dynamical systems model that underlies MINDy, which provides an estimate of effective connectivity. Although regularization generally favors sparse solutions, we found that, even without any regularization, the group average Weight matrix was much sparser than rsFC (SI Sec. 5.7). We provide a simple proof-of-concept to illustrate the potential insights that can be gained from investigating such asymmetries. Specifically, MINDy identified a region of left Inferior Frontal Gyrus (IFG) as the parcel with the greatest asymmetry in positive connections. Specifically, this region showed a positive outward-bias in connectivity with the bias primarily exhibited in its feed-forward positive connections to ipsilateral medial temporal lobe, inferior parietal lobule (IPL), and dorsal/ventrolateral PFC (Fig. 5 C). Excitatory connections of the left IFG with temporal cortex are essential features of the “language network” (e.g. [56]). Additional results revealing other brain regions showing directionality biases in connectivity are reported in SI (Sec. 5.10). In a later section, we explicitly test the robustness of asymmetry estimates and how they are affected by assumptions regarding hemodynamics and model mismatch.

**Fig 5.**
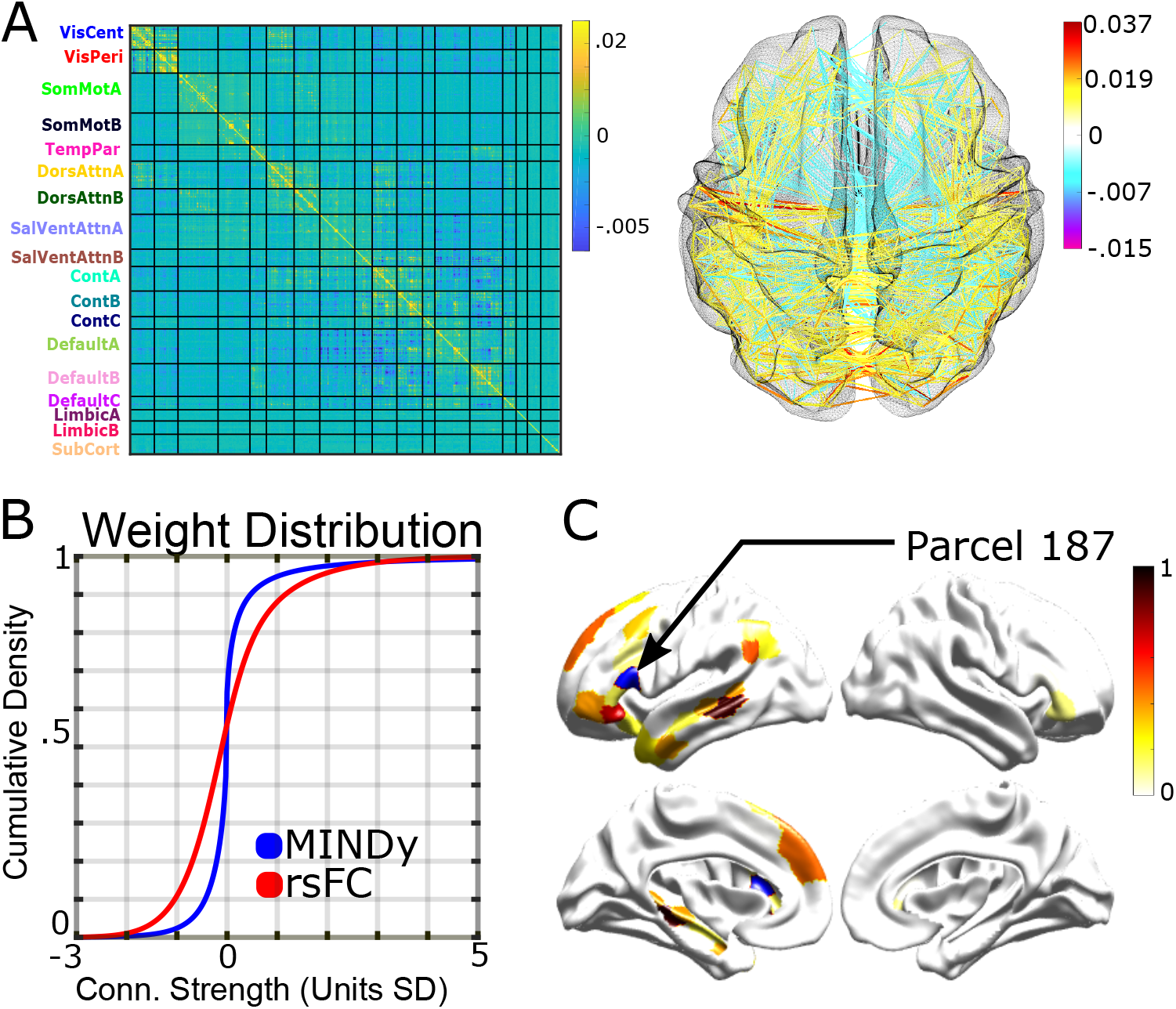
MINDy weights are structured, sparse, and directed. A) Left-side: Mean connection matrix *W* averaged across subjects and scanning session. Parcels are grouped according to the Schaeffer ([21]) 17-network parcellation (hemispheres combined) plus the free-surfer subcorticals. Right-side: thresholded anatomical projection (positive connections≥ 20% max non-recurrent magnitude and negative connections with magnitude≥ 8%). B) The MINDy weight distribution demonstrates sparser connectivity than rsFC. C) Parcel 187 ([21] 17-network), near Inferior Frontal Gyrus, had the strongest source-bias for positive connections (more positive out than in). Plotted surface shows the relative magnitude of this bias (only connections with outward-bias) which largely follows left-lateralized regions implicated in language (e.g. [56]) see SI Sec. 5.10 for additional, preliminary directed-connectivity results. Blue highlights chosen source-parcel.

### 3.6. MINDy Predicts Individual Brain Dynamics

#### 3.6.1. MINDy Predicts Individual Differences in Dynamic Functional Connectivity

We next focused our analyses on the dynamic patterns observed in brain activity, since this has been an area of rapidly expanding research interest within the rsFC literature, termed dynamic functional connectivity or dFC ([57],[58],[59], [60]). Critically, the question of whether MINDy models can predict more slowly fluctuating temporal patterns in the recorded brain data for individual subjects is qualitatively distinct from the ability to predict activity over very short timescales (i.e., 1-step). This is because small biases in fitting individual time points can lead to very different long-term dynamics (e.g. compare panels B and C in Fig. 3, which reflect short and long-term predictions, respectively). To test model accuracy in capturing longer-term dynamic patterns, we used fitted model parameters for each subject to then generate simulated fMRI timeseries, injecting noise at each timestep to create greater variability (see Methods). We then used this simulated timeseries to identify the temporal evolution of short-term correlations between brain regions and compared results with those obtained from the recorded data. Correlation timeseries were estimated using Dynamic Conditional Correlation (DCC; [50]), a method which has been recently shown to improve reliability in the HCP data-set as compared to sliding-window estimates ([61]). We then attempted to recreate DFC measures of individual subjects which have shown the greatest reliability in the actual data. Recent reliability analyses have indicated that simple statistics of temporal variation in individual correlation pairs such as standard deviation of the conditional correlation time-series ([61]) and excursion ([49]) are more reliable than state-based descriptions for the HCP resting-state data ([61]). Therefore, we used these measures (see Sec. 2.8 for equations) to validate dynamics within the model. To avoid confusion, we use the term *σ*-DFC to refer to the temporal standard-deviation of time-varying correlations, which is used as a measure of DFC. Alternatively, the *σ*-DFC may be conceptualized as the signal power of the moving-correlation time series and has proven to be one of the most reliable measures of DFC ([61]). MINDy performed slightly better on recreating another reliable DFC statistic, group-average excursion, so we chose to be conservative and display the results from *σ*-DFC for main-text figures rather than using excursion DFC. Results using excursion DFC and the corresponding figures are provided in SI (Fig. 18).

Results indicate that individual differences in the simulated dynamics of models fit to separate test-retest sessions are at least as reliable as summary dFC measures of individual differences in the original data (SI Fig. 18 A,B). The image intraclass correlation (I2C2, [52]) for the model was .555 for *σ*-DFC and .481 for excursion. In the original experimental data, I2C2 reliabilities were .527 for *σ*-DFC and .380 for excursion. Moreover, individual differences in the DFC of simulated models were highly correlated with those of the original data for most region-pairs (Fig. 6 A). Lastly, we analyzed whether the simulated data recreates the central tendency of observed data. In general, the group-mean *σ*-DFC (SI Fig. 18D) and excursion (SI Fig. 18 E) estimates were highly similar between the simulated and observed data for both the *σ*-DFC (Fig. 6 B; *r*(4948) = .761) and excursion metrics (SI Fig. 18 C; *r*(4948) = .836). Thus, MINDy models recreate measures of DFC at the level of both individual differences and the group-level. Moreover, in some cases (e.g. the excursion metric), MINDy models generate more reliable estimates than those of the original data (SI Fig. 18B). A main advantage of the model in this regard is likely due to the ability to simulate an arbitrarily large amount of data with the model that is also free from nuisance signals/motion.

**Fig 6.**
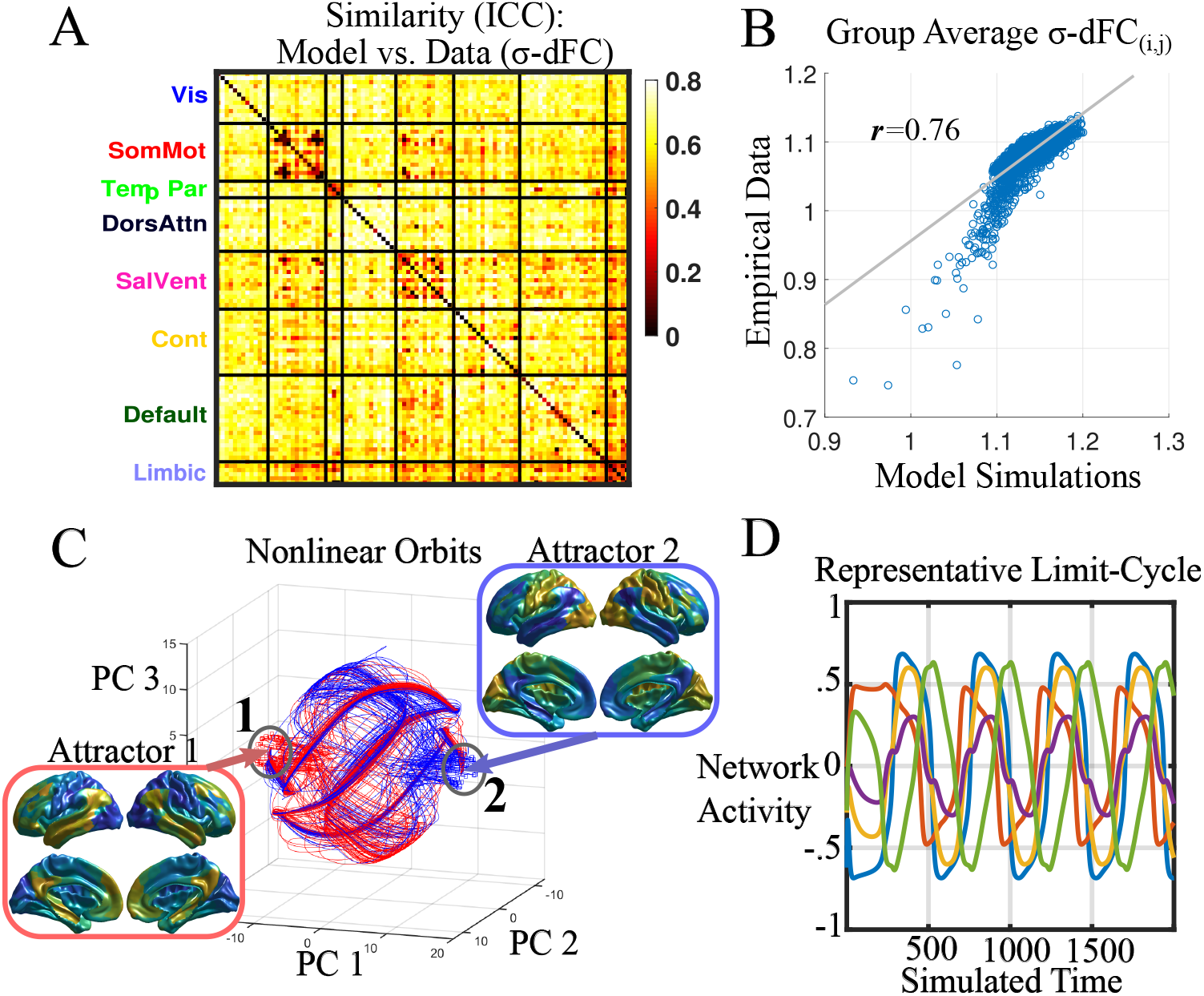
MINDy models predict individual variation and central tendency of pairwise dynamic functional connectivity (DFC) measures. A) Similarity between model and data for predicting each subject’s *σ*-DFC for each pair of brain regions (using the 100-parcel atlas from [21] and collapsing the 17-network grouping down to 8). B) Scatterplot and Pearson correlation of group-average *σ*-DFC for data vs. model. C) Evidence of non-trivial dynamics in MINDy models. Example phase portrait of one subject projected onto the first 3 principal components. Complex orbits link neighborhoods of attractors 1 and 2 (orbits starting from each neighborhood are colored red, blue respectively). Inset figures show these attractors projected onto the brain. D) Example deterministic time-series for a limit cycle in a different representative subject averaged across five networks (Visual [blue], SomMot. [red], Dorsal Attn. [yellow], Salience [purple], and Control [green]). These deterministic dynamics demonstrate significant nonlinearity but are qualitatively different from the simulated model dynamics (e.g. for computing DFC) which include process noise.

#### 3.6.2. MINDy Models Generate Non-Trivial Dynamics

In the previous section we demonstrated that MINDy predicts individual differences in nonstationary dynamics. This finding suggests that the nonlinearities in MINDy are able to account for some features of the data (nonstationarity) that are mathematically absent from linear models. From a dynamics perspective, non-pathological (Schur-stable) linear models predict that spontaneous brain activity consists of noise-driven fluctuations about a single equilibrium. The model parameters for a linear system (e.g. “effective connectivity” in DCM) shape the spatiotemporal statistics of these fluctuations and in the case of white-noise excitation result in a unimodal distribution about the equilibrium in question. Although many nonlinear systems exhibit exotic behavior (e.g. chaos), some systems are dominated by a single equilibrium and may thus possess dynamics that are similar to a linear system. Therefore, we tested whether empirical MINDy models exhibit nontrivial dynamics in the absence of noise (see SI Sec. 5.11). We found that all subjects’ models were dominated by nontrivial dynamics (multistability, homo/heteroclinic cycles, limit cycles, etc.). Example nonlinear dynamics for two representative subjects are provided (Fig. 6C,D), although a thorough characterization of each model’s full phase space is beyond our current scope (see SI Sec. 5.11). Nonetheless, we were able to formally demonstrate that no subject exhibits trivial dynamics (SI Fig. 15A,B; Proposition 2). We conclude that the nonlinearity of MINDy models is not superficial, but rather generates topologically significant dynamics which shape model behavior.

### 3.7. MINDy is Robust

#### 3.7.1. MINDy is Robust to Measurement Noise

We addressed the degree to which MINDy fitted parameters are influenced by potential sources of contamination or artifact in the observed fMRI data. Resting-state fMRI data is thought to be vulnerable to three main contaminants: noise in the BOLD signal, biases induced from post-processing pipelines that attempt to remove this noise, and idiosyncratic variation in the hemodynamic response function that relates the BOLD signal to underlying brain activity. For the first case, we considered two sources of noise in the BOLD signal: additive measurement error and motion artifact. The former case can result from random fluctuations in magnetic susceptibility, blood flow, and responsiveness of radiofrequency coils among other factors. We examined this issue using the ground-truth simulations described above, but systematically varying the amount of measurement noise added at each time-step. This approach allowed us determine how strongly these various sources of noise impacted the ability of MINDy to recover the ground-truth parameters. Results indicated that although the performance of MINDy decreased with the amount of noise added (Fig. 7 A), similarity to the ground-truth values generally remained high. Additional levels of noise are plotted in SI Fig. 20. At the highest level of noise considered, Weight and Decay parameters correlated *R* ≈ 0.7 with ground-truth, while the curvature parameter correlated *R* ≈ 0.6. We note that empirical data exhibiting such a high level of noise would (hopefully) fail quality control.

**Fig 7.**
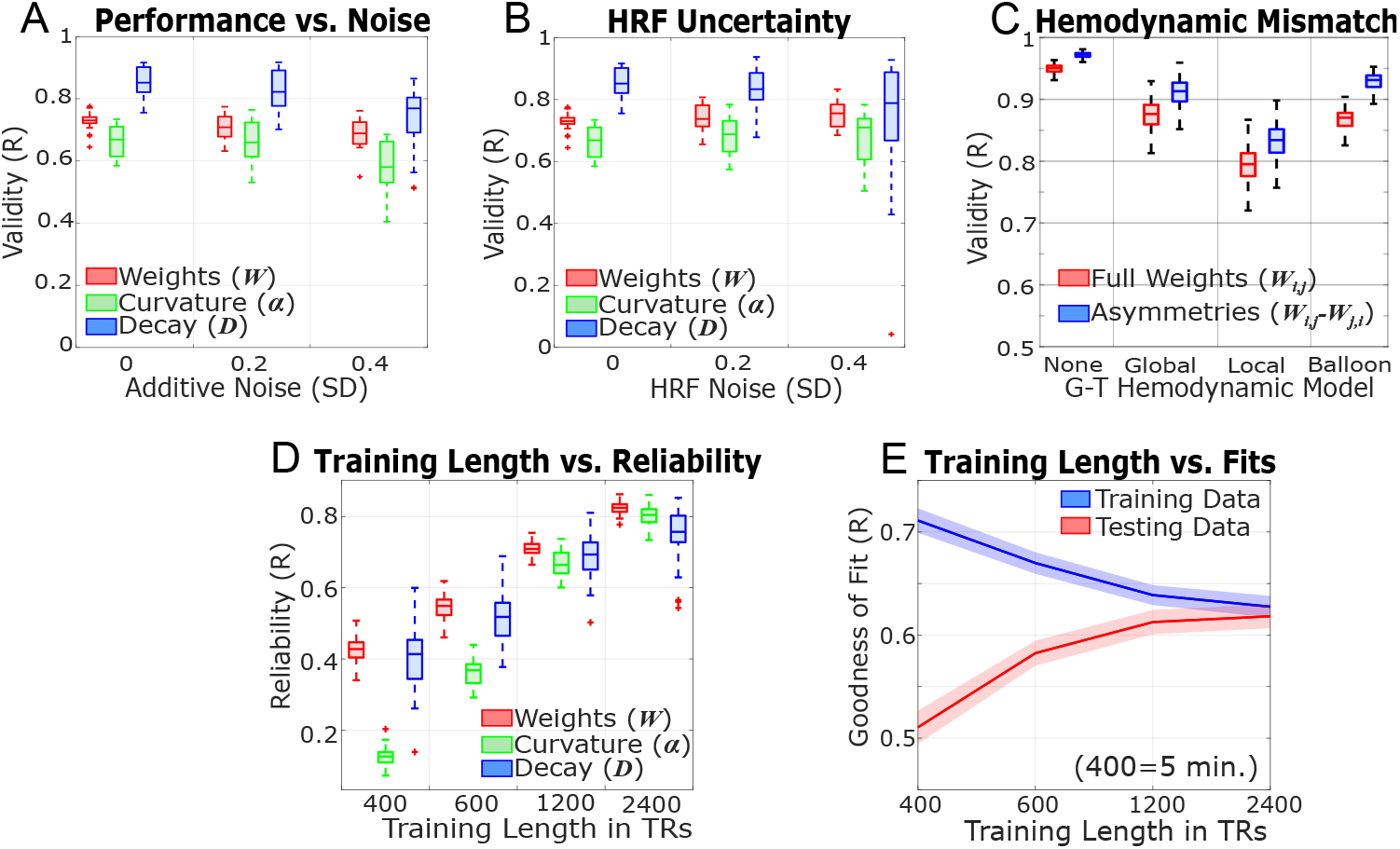
A) Increasing the amount of additive measurement noise slightly decreases MINDy performance in recovering ground-truth parameters. B) Mean performance is unaffected by the uncertainty in the ground-truth HRF, although performance does become more variable (see SI Fig. 20H). C) MINDy performance in retrieving the ground-truth weight matrix (original matrix: red; asymmetric part (*W*_*i,j*_ − *W*_*j,i*_): blue) under mismatch between the ground-truth hemodynamics and the canonical HRF assumed by MINDy (left to right: No hemodynamic modeling, random spatially homogeneous HRF, spatially heterogeneous HRF, nonlinear Balloon-Windkessel model). D) Test-retest reliability of MINDy parameters as the amount of (contiguous) training data is varied. E) MINDy goodness-of-fit for 1-step prediction in training data (blue) and cross-validated with another scanning session (red). The difference between these lines indicates the degree of overfitting. Shading indicates standard deviation.

#### 3.7.2. MINDy is Robust to Individual Differences in Motion

We next examined the impact of motion on MINDy estimates. In this case, we used three standard measures of motion that were derived from the observed fMRI timeseries data: 1) the number of total frames censored due to crossing critical values of frame-wise displacement or DVARS (see Methods), 2) the median absolute framewise-displacement of the subjects head across scanning sessions, and 3) the spatial standard deviation of temporal difference images (DVARS) ([41]). We then examined whether variability in these parameters across individuals contributed to the quality of MINDy parameter estimation and individuation, using test-retest reliability (of estimated parameters from each session) as the index of quality. If MINDy estimated parameters reflected vulnerability to the degree of motion present in an individual, then we would expect higher test-retest reliability in the parameters for the individuals with the lowest estimated motion (e.g., highest data quality). Instead, we found that test-retest reliability was relatively un-impacted by any measure of motion (SI Fig. 19 B-D, I). A parallel analysis used cross-validated fit, in which MINDy parameters were estimated from one session, and then used to predict data in the held-out session, computing goodness-of-fit of the model to the observed data in this session (in terms of variance explained). In this case, we examined a subset of participants that had relatively low motion in one session, but relatively higher motion in the other compared against a second subset that had similar levels of motion in both sessions. If the increased motion in this latter session was problematic, it should reduce the goodness-of-fit (either when used for parameter estimation or when used for cross-validation in the held-out session). In fact, the cross-validated fits were relatively similar in each group (SI Fig. 19 E,J). Together, these results suggest that participant motion (within a reasonable range) may not be strong factor in determining how well MINDy model parameters can be estimated from observed fMRI data timeseries.

#### 3.7.3. MINDy is Robust to Pre-Processing Pipelines

We next examined whether secondary data pre-processing pipelines, which are typically applied to rsFC data prior to analysis, produce biases on MINDy parameter estimates, again examining this issue in terms of test-retest reliability. We considered three variants of a standard published preprocessing pipeline ([28]), one with motion-correction only, one which adds to this CompCor (a standard method that removes noise components associated with white matter and CSF; [42]), and a final, full variant that additionally includes global signal regression (GSR; [62]). We compared test-retest reliability for data-processed with each pipeline (SI Fig. 21 A) and the similarity of parameter estimates obtained when the same data were processed using different pipelines (SI Fig. 21 B). Results indicated that MINDy parameters had high test-retest reliability regardless of preprocessing choices (all *R* > .7, SI Fig. 21 A) and that similar parameter estimates are obtained regardless of preprocessing choices (all *r* > .85, SI Fig. 21 B). By comparison, when a parallel analysis was conducted on rsFC values, the rsFC parameters showed lower test-retest reliability, particularly when more pre-processing was performed on the data, and showed a larger impact of a change in pre-processing on test-test reliability. A direct comparison of the test-retest of MINDy weight parameters relative to rsFC revealed that these were significantly higher (all *p’s* < .05), were more consistent (lower variance of reliability) across pipelines (all *p’s* < .001; Table 8; SI Fig. 21 A), and were less impacted by changing preprocessing pipelines (all *p’s* < .001; SI Fig. 21 B). Together, this set of analyses indicate that the choice of preprocessing pipeline will not have a large effect on estimated MINDy parameters.

#### 3.7.4. MINDy is Robust to Global Hemodynamics

Lastly, we considered the effect of poor estimation of the hemodynamic response function (HRF). Currently, for simplification, the MINDy estimation procedure assumes a canonical HRF model that is constant across individuals and parcels ((although we have recently begun to explore the effect of relaxing this assumption, and estimating a different HRF for each parcel and individual; [63]). Other fMRI models also assume a canonical HRF (e.g. regression-DCM; [19]). However, existing literature suggests that this assumption is likely to be incorrect ([43], [64]). To examine the impact of mis-fitting the HRF, we modeled a variety of ground-truth scenarios. The first set of ground-truth simulations were randomly parameterized according to the empirical MINDy distribution and activity timeseries were convolved with spatially homogeneous, but randomly parameterized HRFs with incrementally greater variability (SI Fig. 20 D). We then attempted to recover MINDy parameters while again assuming the fixed, canonical HRF model ([30]). Results of this analysis suggest that, on average, the MINDy parameters recovered from this analysis remain consistently similar to the ground truth parameters (mean similarity of all parameters, R-value≈0.75, Fig. 7 B). However, the variability of the fits increased across simulations, as the HRF became more variable across regions and individuals (SI Fig. 20 H).

#### 3.7.5. MINDy Parameters are Robust to Model Mismatch

We also considered the effect of violations of the MINDy model in terms of the underlying neural models (MINDy vs. Hopfield, neural mass) and neurovasculature (spatially heterogeneous HRFs and nonlinear hemodynamics). These effects are expected to be most pronounced in estimating asymmetric connections as unaccounted lags can potentially reverse the direction of inferred causality in many other techniques, such as Granger Causality. For the next set of simlations, we generated complex networks from a non-empirical hyperdistribution whose characteristic parameters were randomly sampled at each run. This approach allowed us to sample over a wide range of qualitatively different network structures (Sec. 2.6.2) and these simulations did not depend upon previous empirically-fit MINDy models. We tested the ability of MINDy to recover the weight parameter (Fig. 7 C) from a simple rate model (tanh transfer function) with four levels of hemodynamic variability: 1) no hemodynamics, 2) random, spatially-uniform HRF, 3) random, spatially-heterogeneous HRF, and 4) nonlinear hemodynamics simulated by the Balloon-Windkessel model ([47], [12], Sec. 2.6.4). In the last case, the nonlinear hemodynamic transformation varies implicitly and systematically in space due to spatial variation in the firingrate distribution. Results indicate that MINDy can recover asymmetric connections of ground-truth networks (*W*_*i,j*_ − *W*_*j,i*_) for all cases considered, but performance depends upon the degree of HRF complexity (Fig. 7 C; SI Table 9). When no hemodynamics were included in the model (MINDy received the downsampled neural time-series) performance was near-perfect (*r* = .949 ± .009 overall; *r* = .971 ± .007 for asymmetries, *n* = 1700). Performance also was high for random, spatially homogeneous HRF’s both overall (*r* = .874 ± .024) and at estimating asymmetries (*r* = .910 ± .023, *n* = 1600). Spatial heterogeneity of the HRF decreased MINDy performance in recovering overall ground-truth connectivity (*r* = .793 ± .029; *t*(3071.8) = −86.72, *p* ≈ 0; unequal-variance), but did not differentially impair the estimation of asymmetries (*r* = .832 ± .028; *t*(3057.7) = 11.74, *p* ≈ 1, 1-tailed, unequal variance).

We also found that MINDy still performed well in recovering asymmetric connectivity when a nonlinear (Balloon-Windkessel) ground-truth hemodynamic model (*r* = .865 ± .022 overall; *r* = .927 ± .019 for asymmetries, *n* = 2020) was used to generate simulated fMRI time-series data as compared to when a spatially homogeneous, linear HRF model was used (*t*(3073.3) = 23.03, *p* ≈ 0; unequal variance). Thus, violations of spatial homogeneity in the hemodynamic response appear more relevant to MINDy than violations of hemodynamic linearity. However, performance was still strong in all cases considered (median *r* ≥ .80). We also conducted preliminary tests of MINDy’s ability to recover synaptic conductances (weights) from the simulated BOLD signal (Balloon-Windkessel) of a biophysically parameterized neural mass model ([48]) which evolves at a much faster timescale than the fMRI TR. MINDy was generally able to recover connection weights (synaptic conductance in the neural-mass model) for this case as well (*r* = .684 ± .039 overall). However, unlike in the other simulations, performance in recovering asymmetries (*r* = .624 ± .052) was lower than that of the overall weight matrix (*paired t*(1399) = −109.172, *p* ≈ 0). This result indicates that the difference in time-scales between neuronal and BOLD activity is a more relevant constraint on directional inferences than hemodynamic variability. Although these simulations represent but a small subset of possible ground-truth models, they indicate that the directionality of MINDy connectivity estimates remains largely accurate under violations of the assumed spatially homogeneous hemodynamic response.

### 3.8. Comparing MINDy with Dynamic Causal Modeling

The earlier analyses, in which we compared MINDy and rsFC (Sec. 3.2.1), serve to demonstrate the potential linkages between methods and how MINDy can resolve some ambiguities inherent in rsFC (e.g., directionality). However, these analyses should not be interpreted as stating that MINDy is unambiguously “better” than rsFC as the two approaches represent fundamentally different constructs. The correlation matrix (rsFC) is a statistical quantification whereas MINDy is an approach for estimating a dynamical-systems model and they may have complementary roles for exploring individual differences/biomarker discovery. In order to benchmark MINDy as a model-fitting technique we compared performance with spectral DCM ([7]) in recovering connectivity weights for a variety of simulated ground-truth scenarios. Spectral DCM (spDCM) is a recently developed Dynamic Causal Modeling (DCM) approach for simultaneously estimating linear dynamical systems and (region-specific) hemodynamic kernels from resting-state fMRI ([7]). To be clear, we view the primary contributions of MINDy relative to modeling approaches such as spDCM in terms of its scalability, biological interpretability, and the ability to predict nonstationary resting-state dynamics. However, the question remains whether these advantages come at the expense of accuracy—i.e. whether MINDy is inferior to DCM within the latter’s scope.

We compared performance of MINDy and spDCM across a variety of ground-truth scenarios (see SI Sec. 5.8) to test whether MINDy performs at least as well as spDCM in the lower-dimensional scenarios in which the latter is applicable (i.e., estimating parameters for a small number of nodes or neural masses). These simulations were specifically designed to reduce bias based upon either model’s assumptions (see SI Tab. 1) and considered ground-truths based upon mesoscale Hopfield-style models (SI Fig. 12A) and biophysical neural mass models (SI Fig. 12B). In the former case, we manipulated the degree of spatial variability in the hemodynamic response (SI Fig. 12C). When arbitrary choices were necessary, we chose the option than empirically favored spDCM. Results support that MINDy’s advantages do not come at a cost to accuracy. In all settings considered, MINDy was at least as accurate, on average, as spDCM and significantly (orders of magnitude) faster. We observed that spDCM was more robust than MINDy to spatial variability in the ground-truth HRF (although see extensions in [63]), but even under the most extreme cases considered, MINDy was at least as accurate as spDCM (SI Fig. 12). The empirical examination of run-time overwhelmingly favored MINDy (SI Fig. 12D-F). For example, the largest network we tested contained 16 neural masses (SI Fig. 12D) for which MINDy estimation took 3.5s on average vs. 2.7 hours for spDCM. We estimate that fitting spDCM models using our chosen parcellation, involving 419 brain regions/nodes (400 of which are cortical [21]) would take a minimum of 44 years each (and likely much longer; see SI Sec. 5.8). We conclude that MINDy’s advantages (scalability, dynamics etc.) do not come at the expense of accuracy relative contemporary approaches.

### 3.9. MINDy Requires 15-20 minutes of Data

In most fMRI experiments scanner-time is a precious resource and particularly so with sensitive populations. While the Human Connectome Project affords a full 60 minutes of resting-state scan time, this quantity of data may not be a reasonable expectation for other datasets, so we varied the training data length to determine how much data was necessary for MINDy to reliably estimate models. We first evaluated reliability in terms of test-retest on MINDy parameters estimated from separate scanning days using only a subset of the total data for model fitting. As expected, when the length of data used to estimate parameters increased, the test-retest reliability of the estimated parameters also increased, up to the maximum interval considered (30 minutes). Nevertheless, acceptable levels of reliability (*R* ≈ .7) were obtained with 15 minutes of data (Fig. 7 D). We next examined bias or overfitting of MINDy parameters by comparing the fit to the trained data, relative to a cross-validation approach, examining the fit to held-out (testing) data. As would be expected with over-fitting bias, as the length of the training data increased, the fit to the trained data decreased. In contrast, the fit to the held-out (test) data increased, and the two values converged at around 15 minutes of data in training and test sets (Fig. 7 E). Thus, we do believe the current method is too prone to over-fitting biases and unacceptably low reliability with fewer than 15 minutes of total scan time using the HCP scanning parameters. However, since the data does not need to be acquired in a single continuous run, we believe that this requirement is reasonable and in concert with current recommendations for rsFC analyses ([53],[54]). Future study with other fMRI acquisition techniques may illuminate how data-requirements change with sampling rate (e.g. shorter TRs may potentially compensate for less scan time).

## 4. Discussion

### 4.1. Relationship with Functional Connectivity

There are three primary advantages to using MINDy over rsFC. First, rsFC is limited as a statistical descriptive model. This means that even though rsFC may be found to be reliable and powerful as a biomarker that can characterize individuals and effects of experimental variables, it is unable to predict how the nature of an experimental manipulation relates to the observed changes in brain activity. By contrast, MINDy is a true mechanistic causal neural model, which attempts to capture the physical processes underlying resting-state brain activity in terms of neurobiologically realistic interactions and nonlinear dynamics ([8]). This feature is powerful, as it enables investigators to perform exploratory analysis in how altering a physical component of the brain (e.g. the connection between two brain regions) will affect brain activity ([11]).

Second, MINDy provides a richer description of brain mechanisms than rsFC. While rsFC and MINDy both attempt to parameterize the connection strength between brain regions, MINDy also describes the local mechanisms that govern how each brain region behaves. Neural processes are thought to involve the combination of anatomically local computations and spatially-extended signal propagation, so it is important that descriptions of brain activity be able to define the degree to which this activity is generated by local vs. distributed mechanisms. Although the elements of the rsFC matrix are often interpreted as reflecting interregional components of neural processing, we have demonstrated that the rsFC is also sensitive to purely local characteristics of brain regions, such as their decay rate (SI Fig. 17 A). Conversely, we have demonstrated that both the transfer function and decay rate of brain regions can also serve reliable markers of individual differences and anatomical structure. By using MINDy, researchers can identify which neural mechanisms (i.e. which of MINDy’s parameters) give rise to individual differences of interest.

Lastly, MINDy greatly improves upon rsFC’s characterizations of effective connectivity between brain regions. Unlike the elements of a correlation matrix, MINDy’s weight parameters can describe asymmetric connectivity strengths and thus make inferences regarding the directional flow of activity between brain regions. We provide tantalizing illustration of the potential utility of these types of findings (e.g., Fig. 5 C, SI Fig. 14). Further, we demonstrated that MINDy may prove a more valid measure of brain connectivity and individual differences in connectivity than rsFC (Fig. 2 E).

### 4.2. Relationship with other Models

There are currently two classes of generative models used to study fMRI: proxy-parameterized neural-mass models (e.g. models using diffusion-imaging data as a proxy for synaptic efficacy [9], [11]) and directly-parameterized linear models (e.g. Dynamic Causal Modeling [14], [7]). Although nonlinear variants of DCM have been proposed for task-fMRI ([65]), the techniques used for resting-state fMRI (e.g. [7]) are fundamentally linked with the statistics of linear systems. These two approaches (proxy-parameterized neural-masses and linear models) represent opposite ends of a trade-off between realism and tractability for mesoscale human brain modeling. The first case (proxy-parameterized neural-mass models) excels in terms of interpretability of the model framework, since the state-variables can always be traced back to population firing rates. These models operate at relatively fast time-scales and can produce predictions with the spatial resolution of MRI and the temporal resolution of EEG, which make them a parsimonious and general-purpose investigative tool that can be utilized across temporal scales. Current approaches in this respect are limited, however, in the manner by which these models are parameterized. Even in state-of-the-art techniques (e.g. [12],[13]) most parameters are fixed a priori (local neural-mass parameters), or determined from diffusion imaging data, with only a limited subset taken from fMRI functional connectivity estimates, typically at the group-average level. Thus, the vast majority of parameters are not sufficiently constrained by the relevant individual-level data, and instead are adapted from measurement of proxy variables, which is likely to limit the accuracy of model predictions. Diffusion imaging data, for instance is inherently unsigned and undirected, so the resultant models are unable to consider hierarchical connection schemes or long-distance connections that depress activity in the post-synaptic targets. Moreover, it remains unknown how to convert units from white matter volume to synaptic conductance even when these assumptions are met. In practice the conversion is performed by choosing a single scaling coefficient, which assumes that this relationship is linear with a universal slope. Due to these sources of uncertainty, proxy-parameterized models do not necessarily fit/predict raw functional time series. To be fair, however, this limitation may not be relevant for all scientific questions (e.g. studying long-term phenomena such as FC, [22], [12], [13]).

By contrast, the ability of a model to fit the observed time-series implies that its predictions are valid within the vicinity of observed data. This property holds even when the underlying model is likely to be inaccurate in its long-term predictions. Evidence of this can be seen in the success of dynamic-causal modeling approaches, which can recreate task-driven activity ([66]) despite using a simplified linear model. However, the downside of using a linear modelling framework is that the long-term predictions of these models are most likely inaccurate, given that brain activity in a linear model will always converge to a noise-driven stationary distribution. Thus, even though these models may be more accurate than forward-parameterized neural-mass models in their short-term predictions (by virtue of fitting parameters), the linear form guarantees that they will be unable to capture the extended pattern of brain spatio-temporal dynamics. Analytically, it is known that linear dynamical systems cannot exhibit non-trivial deterministic dynamics and are characterized by a steady-state covariance when driven by noise (which can be calculated by solving a Lyapunov equation). For this reason, they are often employed as surrogate models for testing whether proposed measures of DFC can distinguish between noise-driven trivial (linear) dynamics and those observed in the data ([67], [68]). Thus, DFC measures which have been shown to be non-spurious through surrogate methods cannot, by definition, be reproduced by a linear dynamical system with or without noise . Likewise, these models will also be limited in their ability to identify the neural mechanisms underlying predictions. Since the model takes a reduced (linear) form, it remains unknown whether the coefficients fit to the linear models are the same as would be retrieved by fitting a more realistic model (e.g. neural mass model). That is not to say that the coefficients are uninterpretable; indeed meaningful predictions have been made by inferring effective connectivity from the model coefficients (e.g. [7]). On the other hand, the models’ simplicity may lead to nonlinear components of brain activity being mixed into the linear model estimates, just as we have shown that intrinsic dynamics influence FC estimates (SI Fig. 17 A).

MINDy attempts to leverage the advantages from both approaches. Like current neural-mass models, MINDy employs a nonlinear dynamical systems model which is capable of generating long-term patterns of brain dynamics. However, MINDy is also a data-driven approach, in that models are fit from the ground-up using functional time-series rather than using surrogate measures such as diffusion imaging (although in principle, such information could be used to initialize or constrain MINDy parameter estimates). From the perspective of biologically-plausible models, MINDy extends parameter fitting from the relatively small number of local parameters that constitute the current state-of-the-art ([12], [13]) to fitting every parameter in biologically-plausible individualized whole-brain mesoscale models (i.e., increasing the number of fitted parameters by orders of magnitudes). Likewise, MINDy extends data-driven modeling of resting-state data from linear models containing tens of nodes ([7],[19]) to nonlinear models containing hundreds. It is also worth noting that the computational innovations made in the optimization process make MINDy parameterization many orders of magnitude faster than comparable techniques ([13],[7]; see SI Sec. 5.8) despite fitting many more parameters (e.g.,over 176,000 free parameters can be estimated in a minute vs. several hours to fit hundreds of parameters). This efficiency has enabled us to interrogate the method’s validity and sensitivity in ways that would probably not be computationally feasible for less efficient methods (e.g., building error distributions for sensitivity analyses in Sec. 3.2.1).

### 4.3. Comparison with Diffusion Imaging Seeded Neural Mass Models

Although we emphasize the ability to generate individualized brain models, previous studies using neural-mass models with weights seeded by diffusion imaging have emphasized predicting group-level data ([22],[12],[13]). Two recent studies fit free parameters with the explicit optimization objective of predicting the group-average rsFC matrix ([12],[13]). By contrast, MINDy seeks to predict the short-term evolution of the neural activity time series for single subjects, which often results in the simulated individual rsFC correlating highly with the empirical rsFC (Fig. 3 C). Averaging across simulated rsFC’s produces a group-level simulated rsFC that correlates extremely well with the empirical group-average rsFC (*r*(87, 398) = .94; see SI Fig. 16). As such, the group-average MINDy fit compares very favorably with the analogous measures for diffusion-parameterized models which typically don’t surpass *r* = .6 ([69],[12],[13]).

### 4.4. Limitations

There are two primary limitations of MINDy. The first relates to the properties of fMRI data: MINDy is limited by the spatial and temporal resolution at which data is gathered. This means that MINDy is more sensitive to slow interactions that occur over larger spatial scales and is limited to predicting infraslow dynamics (as opposed to higher-frequency bands). Interactions that are more heterogeneous in time or space may also be missed by MINDy as the model assumes that the transfer function is monotone. While the strength of signaling between regions is allowed to vary according to the transfer function, the sign of signaling (inhibitory vs. excitatory) is not. Thus, MINDy cannot describe relationships which, depending upon local activity, change sign from net excitatory to net inhibitory. This feature is inherent in region-based modeling and so this limitation is not unique to MINDy.

A second limitation relates to the model used to specify MINDy. Unlike the conventional neural mass models ([8]), MINDy employs a single population rather than two or more local subpopulations of excitatory and inhibitory neurons. The model does contain local competitive nonlinearities in that the decay term (*D*) competes with the recurrent connectivity of *W* but the precise mechanisms underlying these dynamics are not explicated. By comparison, multipopulation neural mass models contain separate terms for the interactions between subpopulations and the time-constants of firing rate within each subpopulation, both of which likely influence the local parameters of MINDy. Similarly, while MINDy can specify that the directed interaction between a pair of regions is positive, it cannot distinguish between excitatory projections onto an excitatory subpopulation and inhibitory projections onto an inhibitory subpopulation (both of which could be net excitatory; see SI sec 5.1).

### 4.5. Future Applications and Directions

We view MINDy models as providing a rich and fertile platform that can be used both for computationally-focused explorations, and as a tool to aid interrogation and analyses of experimental data. The most immediate potential of MINDy is in providing new parameter estimates for studies of individual differences or biomarkers. There is also immediate potential for MINDy in model-driven discovery of resting-state dynamics (e.g.[48]), in which case MINDy simply replaces diffusion imaging as a method to parameterize neural mass models. The potential benefit of using MINDy over diffusion imaging is that MINDy identifies signed, directed connections in a data-driven manner which may improve realism. Going forward, other applications of MINDy may be in bridging the gap between resting-state characterizations of brain networks and evoked-response models of brain activity during task contexts. We envision two lines of future work in this domain: one in improving estimates of task-evoked effects, and the other concerning the effect of task contexts or cognitive states on brain activity dynamics.

#### 4.5.1. Isolating Task-Evoked Signals

One future use of MINDy may be in improving estimates of task-related brain activity. Current methods of extracting task-related brain signals are based upon comparing BOLD time courses during windows of interest using generalized linear models. However, the effects of task conditions are related to both ongoing brain activity ([70]) and intrinsic network structure ([71]). Viewing the brain as a dynamical system, any input to the brain will have downstream consequences, so brain activity observed during task contexts likely contains some mixture of task-evoked activity and its interaction with spontaneous activity. Using MINDy, it may be possible to isolate task-evoked responses by subtracting out what would have been predicted to occur via the resting-state MINDy model. The resultant estimate for task-related activity would be the time-series of MINDy prediction errors (i.e. residuals), ideally adjusted for the model’s error at rest. In forthcoming work, we have been using MINDy to estimate task-related activity in this manner, and the initial results strongly indicate that this approach improves the statistical power and temporal specificity of estimated neural events ([72]). Thus, MINDy has the potential to improve estimates of task-evoked activity from fMRI data, although future validation is needed.

#### 4.5.2. Illuminating Dynamics

Present results indicate that MINDy is able to replicate some features of infraslow brain-dynamics observed in the data (see Sec. 3.6.1). Although these slower frequency bands have been less studied in task-contexts, growing evidence implicates them in slowly evolving cognitive states such as states of consciousness ([73],[74]) and daydreaming ([75]). MINDy may benefit future studies in these domains by providing a formal model by which to identify the mechanisms underlying dynamical regimes. Moreover, MINDy may illuminate the behavioral significance of infraslow dynamics. Previous studies have found that timing of pre-cue brain activity and infraslow dynamics interact to predict behavioral performance ([76],[77],[78]), so future characterizations of task-activation may benefit from considering how exogeneous stimuli interact with endogenous neural processes. Generative models of resting-state brain activity may prove critical in these efforts by predicting how endogenous brain states modulate the effects of exogeneous perturbations.

#### 4.5.3. Conclusion

We have developed a novel and powerful method for constructing whole-brain mesoscale models of individualized brain activity dynamics, termed MINDy. We demonstrate that MINDy models are valid, reliable, and robust, and thus represent an important advance towards the goal of personalized neuroscience. We provide initial illustrations of the potential power and promise of using MINDy models for experimental analysis and computational exploration. It is our hope that other investigators will make use of MINDy models to further test and explore the utility and validity of this approach. Towards that end, we have made MATLAB code and documentation for developing and testing MINDy models available via the primary author’s GitHub: https://github.com/singhmf/MINDy-Beta.

## 5. Supplemental Information

### 5.1. Interpreting Model Parameters

As discussed in the main text, the neural modeling framework of MINDy is inherently phenomenological in that it is not directly derived from biophysical first-principals. The weight parameter (*W*), for instance, serves to measure effective connectivity and should not be confused with synaptic efficacy (or any other directly measurable anatomical metric, such as white matter integrity). The phenomenological nature of these equations gives tractability to the fitting problem. However, this fact does not preclude the model’s interpretable and predictive nature. The parametric form that we have chosen leads itself to interpretability by separating the dynamics into three distinct components, interregional-signaling, local decay, and a nonlinear mapping between local excitation and output, which parallel the components used in conventional neural mass models. In the following subsections, we present fuller descriptions of the potential relations between model parameters and underlying biological processes.

#### 5.1.1. Interpreting Model Weights

In our model, the connectivity matrix defines the causal ability of mean regional activity in the sender region to monotonically change mean regional activity levels in the receiving region within a specific time window. This causal influence has standardly been termed effective connectivity within the fMRI (and EEG) literature. More precisely, however, in the model the effects must begin within the duration of one TR (720ms in our case), and last long enough to invoke a metabolic response. As such, our definition is slightly more specific than the notion of effective connectivity, as we specify that these relations must be weakly monotone: all else being equal, increasing (decreasing) the activity of region A will never decrease (increase) the activity of region B (Fig. 8 A, B). We use the term weakly monotone as regions may exhibit saturated activity within our model and thus have little room to increase/decrease. In contrast with our definition, effective connectivity does not specify the nature of the relationship between regions. In this case, model-free methods such as transfer entropy ([79]) can be employed to study non-monotone relationships within a very small number of dimensions. We also define our temporal range of interactions to be between 500ms and 2s. We do not use a tighter temporal range such as 500ms-1s as temporal variations inherent in BOLD imaging, such as physiological changes in the hemodynamic response (e.g. under anesthesia; [80],[81]) lead to some uncertainty in timing. In addition, there are methodological limitations inherent in rapid acquisition methods such as multiband imaging ([82]), which have led some investigators to prefer TR’s closer to 2s. In either case, our definition limits the duration of interest to the order of a typical fMRI trial.

**Fig 8.**
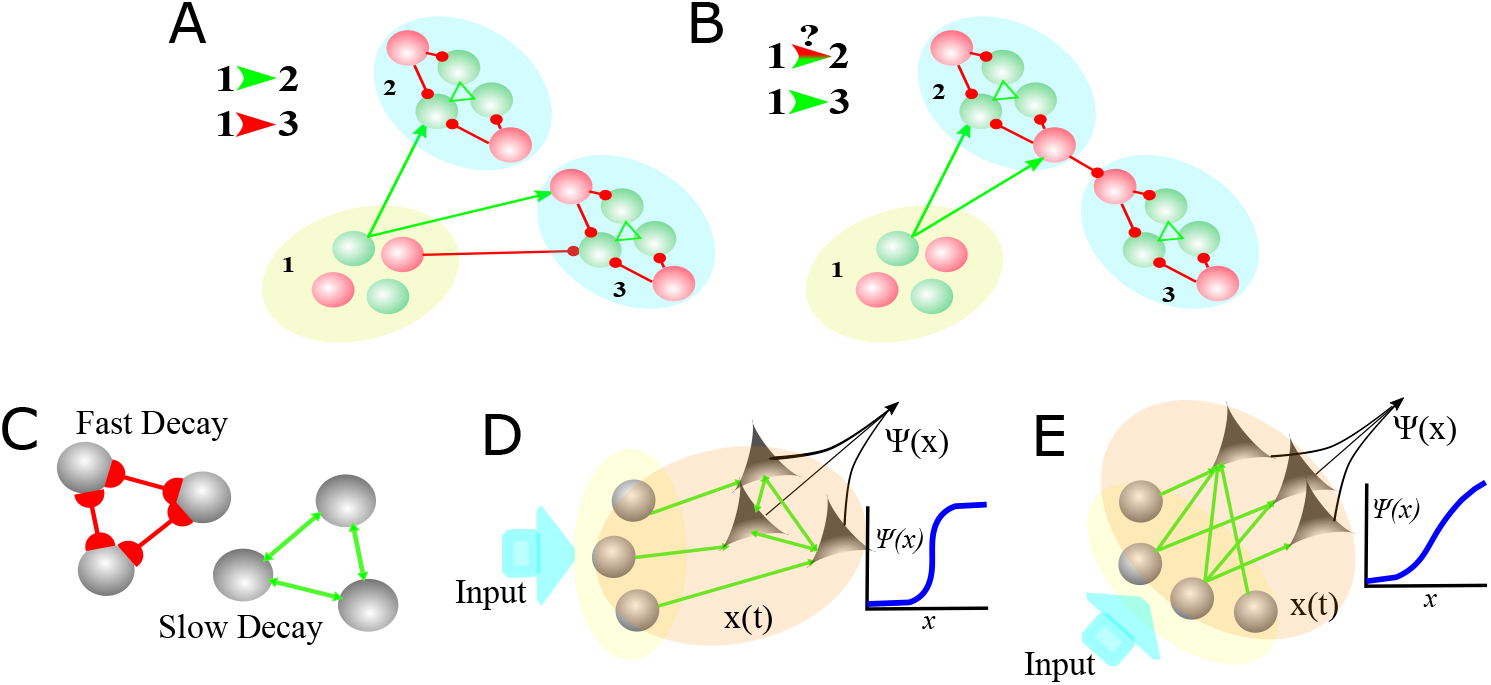
Interpreting Phenomenological Model Parameters. A) The weight matrix largely captures monotone causal relationships. However, the sign of the causal relationship depends upon the sign of the actual inter-regional connection and which neurons are involved. Excitatory connections/cells are depicted with green arrows and inhibitory in red dots. B) When the sign of connections between regions is mixed, it is possible for indirect relationships to appear stronger than direct connections. Local network structure could influence transfer function and decay parameters. C) Networks with greater reciprocal inhibition (red lines) have a faster time-scale, hence greater decay than those with reciprocal excitation (green). D) A toy example of a network with near binary output due to reciprocal excitation in the output cells (triangles). E) A toy example with a more graded output rule due to inhomogeneities in the excitation of output cells.

The monotone and temporal constraints can also differentiate our W matrix from structural connectivity, the latter of which does not necessarily reflect how regions interact. If two regions communicate in a very heterogeneous manner and/or these interactions only result in very transient changes, these regions would not be connected in our W matrix, even if a direct white matter tract linked them (Fig. 8B). Of course, this scenario also suggests that those portions of brain would also not meet the definition of a cortical parcel due to their heterogeneity. Finer parcellation schemes lead to correspondingly more homogeneous “regions”, so, with a sufficiently high resolution parcellation, we expect that most forms of structural connectivity would meet the monotone requirement, with a single cell as the theoretical limiting case. In summary, our form of connectivity in the W matrix describes not just the ability of regions to causally influence each other but to do so with easily predictable (monotone) consequences in a specific time scale. For ease of presentation, however, we use the term “effective connectivity” to refer to this matrix and also make connection with existing terminology.

#### 5.1.2. Interpreting Model Curvature

In the original theory of neural mass models ([31], [83]), the decay-term and transfer function were meant to capture phenomenological components of the individual population without corresponding to a singular biological feature. For instance, the transfer function of neural mass models is usually derived from the probability of neuronal spiking as a function of excitation. If cells within each population are assumed homogeneous, the population level activity is proportional to the individual spiking probability when refractory periods are negligible. Under this homogeneity assumption, inter-parcel variation in the transfer function slope would directly reflect variation in the cellular spiking probability between parcels. For cortical neurons with low-firing rates at rest, the spiking threshold is essentially constant (unlike bursting cells for instance), so a high slope might be interpreted as low noise. Since the ground-truth relation with excitation is binary for each cell at a given time (“all or none” spiking), all deviations from that relation must be due to variations in how much of the population-level excitation each neuron receives.

However, if we instead allow parcels to be internally heterogeneous, the transfer function slope parameter may indicate heterogeneities in either the spiking threshold or how excitation is distributed within the parcel (Fig. 8 D, E). For a simple leaky integrate-and-fire model of neurons the individual transfer functions are binary (infinite slope). However, as the variation in firing thresholds between cells increases, the cumulative probability of population spike count becomes more graded corresponding to the sum of binary functions with different thresholds. Other sources of variation such as noise or inhomogeneities in projections to cells within the population would have a similar effect (Fig. 8 E). Thus, although the exact source of variation (i.e., between regions or individuals) in the transfer function slope is unknown, a likely contributor is the degree of within-parcel variation, which may be due to inhomogeneities in internal/external inputs or neuronal dynamics.

However, there are at least three other potential physiological influences in transfer function slope. The first is the relationship between neuronal activity and the BOLD response. The neural components of the BOLD signal are more closely related to synaptic activity than neuronal spiking, so the likelihood of synaptic activity achieving a spike may also be a factor. For instance, for a given number of excitatory synaptic events, the likelihood of the post-synaptic cell firing generally increases with the synchrony of these events. Thus, the degree of synchronization could be another factor in the transfer function slope with parcels having greater synchronization of excitatory inputs having a higher slope. Alternatively, variation in neurovascular coupling between regions may affect the relationship. Regions with less predictable or less uniform hemodynamics would likely receive a lower transfer function slope similar to the case of neuronal variation. In this case, however, the lower slope results from uncertainty in observations rather than variation (“noise”) within the generative system.

A final factor may be the intrinsic dynamics of each population. As the BOLD-based observations are temporally coarse (i.e., low resolution), the activity level of each population is more reflective of the average level of synaptic activity over hundreds of milliseconds. Thus, the transfer function seeks to relate the sum of parcel output over hundreds of milliseconds to the sum of parcel input (internal and external) over hundreds of milliseconds. Populations with more temporal integration (better “memory”) are less sensitive to variation in input timing so transfer function slope might also increase with parcel memory. However, results actually indicated the opposite: parcels with greater slopes consistently had parameters reflecting less temporal integration (larger decay; see Results). Temporal integration within our model is reflected by the decay parameter, with high decay indicating less temporal integration.

**Fig 9.**
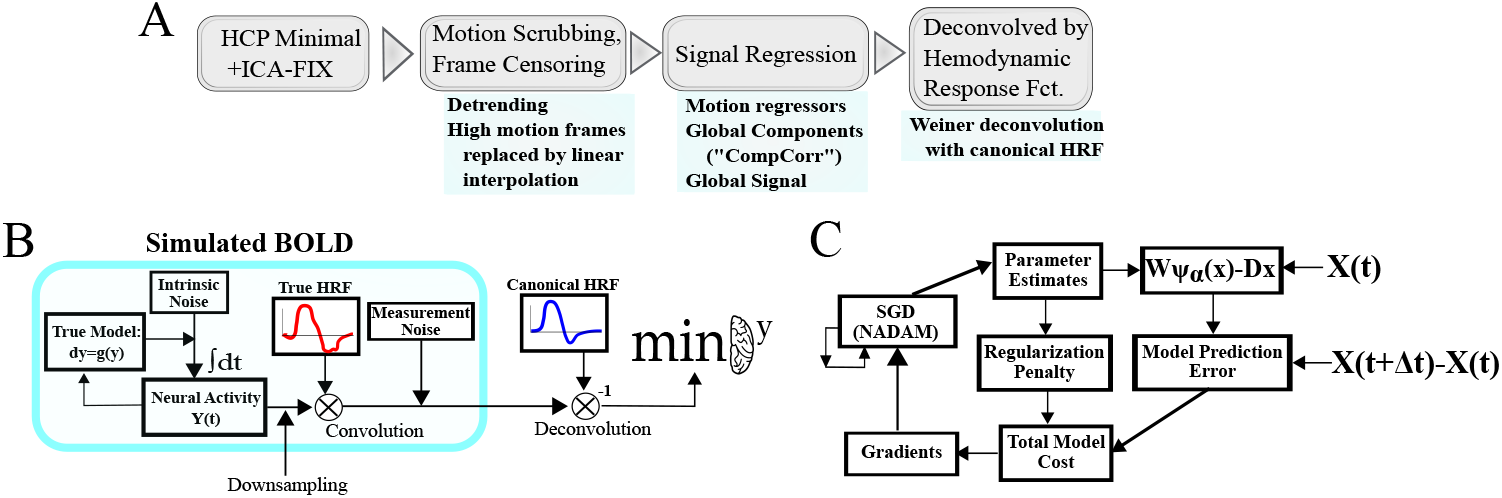
Schematics of data-processing and data-generation pipelines. A) Secondary preprocessing consisted of the pipeline proposed by Siegel and colleagues ([28]) followed by Weiner deconvolution ([29]) with a canonical HRF function. B) Simulated BOLD signals were generated by integrating MINDy models as stochastic differential equations, downsampling results to the scanner TR, convolving with a (potentially random) HRF and adding measurement noise. The resultant signal was then deconvolved with the canonical HRF. C) MINDy parameter estimation consists of iteratively updating estimates using current and past error gradients according to NADAM ([34]).

#### 5.1.3. Interpreting Model Decay

For neural mass models, the decay term describes how quickly a homogeneous population returns to its baseline level of activity. It is assumed that, in the absence of external inputs, the time course will be exponential, leading to the linear term −*Dx*. Many cellular models also contain a linear decay term corresponding to the leak current, with *D* equal to the membrane time constant. At the population level, however, the decay term cannot be easily related to any biophysically comparable parameter, e.g., leak potassium conductance. Instead, the decay parameter should be considered as a phenomenological fit to the general pattern of homogeneous populations returning to some baseline rest level. In the current model, however, we relax the assumption of linear decay by also allowing “self-connections” in the connectivity matrix. That is not to say that the individual population members (neurons) contain autoconnections, but that by allowing both a nonlinear term and a linear term we allow a greater range of possible intrinsic dynamics including self-excitation at the population level (Fig. 8 C). When the model is fit to the HCP data, all individuals were found to display nonnegative values for the nonlinear self-interaction. The resultant intrinsic dynamics for each isolated parcel consist of a nonlinear self-excitation and a linear self-inhibition which can lead to either a single stable equilibrium (near the mean BOLD signal) or bistability wherein initial conditions sufficiently above the mean will all converge to one equilibria and those sufficiently below the mean converge to another. The bistable case generally results when the maximal slope of the self-excitatory component is larger than the decay term (see Supp. for precise conditions). In general, we expect that the decay parameter is related to the relative proportions of local excitation/inhibition within each parcel (Fig. 8 C). The anatomical distribution of decay terms across parcels was largely consistent across subjects (Fig. 4B).

### 5.2. Derivation of the MINDy Transfer Function

The use of a new transfer function is motivated by the desire to unify the three main classes of transfer function employed in both artificial neural networks and biological neural models: the rectified linear unit (ReLu), softplus, and the logistic sigmoidal function. These functions differ in their curvature (ReLu is piecewise linear, while the others are smooth) and their boundedness (ReLu and softplus are unbounded, linear in the positive limit). Rather than specifying one of these functions explicitly, as is usually done, we chose to create a more general functional form and let the data select the function’s shape on a person x region basis. This form consists of a generalized class of sigmoidal functions which can be varied from smooth to piecewise-linear. We’ll later show that this property enables approximation of the other two classes (ReLu/softplus) over bounded domains. Our function is generated by integrating the difference of two shifted sigmoidal functions (denoted *σ*(*y*)).

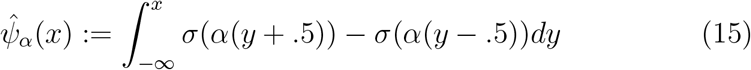

By Proposition 1 (below), the shift by ±.5 guarantees that 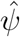 will have the same limits as the original function *σ*(*x*) and also retain any of the original function’s reflection symmetries about *x* = 0. Moreover, 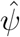 reduces to the definite integral (Proposition 1):

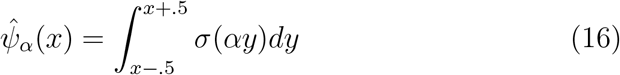

When *α* is small, this formulation generates smooth sigmoidal functions with curvature dependent upon the choice of *σ*. However, in the limiting case of large *α*, the function approaches a shifted ReLu function for sigmoids (*σ*) that have a lower-limit of zero and appropriate rescaling (*b*):

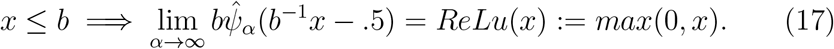

For finite values of *α, b* the function is smoothed and can be rescaled/shifted to behave like a soft-plus function over desired intervals. An analogous means of generating functions was previously used for modeling dendritic saturation ([84]) starting from the logistic sigmoid function:

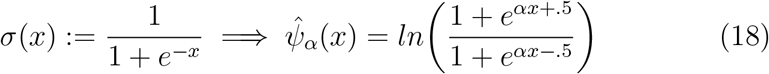

We chose to use the sigmoidal function:

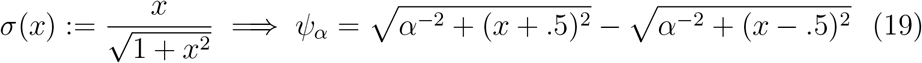

which takes values on (−1, 1) similar to the hyperbolic-tangent (tanh). We favored the chosen sigmoidal basis over tanh/logstic (as was used in [84]), because the resultant transfer function *ψ* involved slightly faster and more stable operations (i.e. avoided using log). Additional terms to further customize the slope/intercept of the transfer function were initially considered, e.g.:

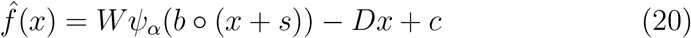

However, we observed in early tests that the scaling term could be reduced to a scalar constant (*b* = 20/3) and fitted values for *s* were effectively zero (for z-scored data). We also observed that when transfer functions were bounded over [−1, 1] the *c* term became effectively zero which was not the case when we tested transfer functions bounded over [0, 1]. Thus, we chose to use functions bounded over [−1, 1] so that the *s* and *c* terms could be removed. Of course, MINDy models can always be rewritten in an equivalent form featuring a non-negative transfer function and constant drive

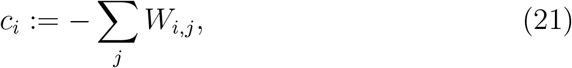

since *ψ*(*x*) + 1 is non-negative and

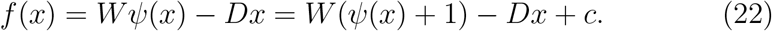

#### Proposition 1.

*Define the operator* Φ : *C*^0^ → *C*^1^:

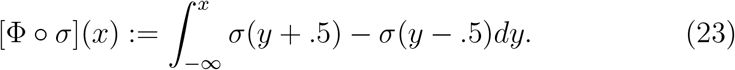

*Suppose that σ is non-decreasing and bounded. Then* lim inf[*σ*] = lim inf[Φ ○ *σ*] *and* lim sup[*σ*] = lim sup[Φ ○ *σ*]. *Moreover*,

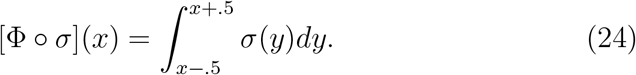

*Proof.* By Lebesgue’s Theorem for Monotone Functions, *σ* is differentiable almost everywhere and we can write a function *Dσ* equal to the derivative of *σ* at differentiable points and zero otherwise which satisfies:

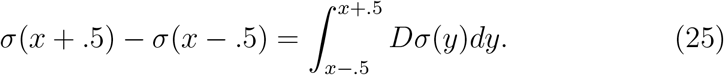

By rearranging limits of integration we produce:

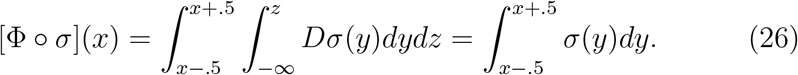

By monotonicity we have:

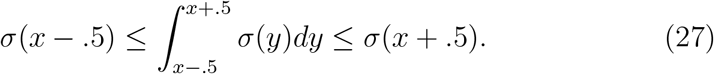

By the monotone convergence theorem, *σ* converges to its infimum/supremum. Taking the negative limits for *x*:

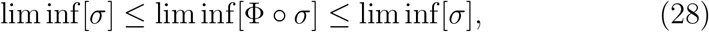

and similarly for the positive limit (lim sup). Applying the squeeze theorem completes the proof.

### 5.3. Accelerated Stochastic Gradients through NADAM

To fit the models, we use a variant of the stochastic-gradient descent (SGD) method: NADAM (Nesterov-accelerated adaptive moment estimation [34]) which builds upon the earlier ADAM algorithm ([85]). Gradient descent methods are algorithms that attempt to minimize a cost function, by updating parameters based upon the cost function’s current slope (gradient). For an error function *E* and a parameter *θ*, the original gradient descent algorithm updates the estimate of the parameter (denoted *θ*_*k*_) at each iteration (*k*) of the algorithm according to:

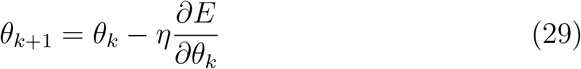

In which *η* is the user-chosen learning rate parameter. Although highly efficient, gradient descent algorithms are not guaranteed to reach a global minimum for non-convex problems; further, the original gradient-descent method is prone to getting “trapped” in local minima. Additionally, global-minima of highly non-convex problems may not be desirable as they sometimes poorly generalize (10). Since the development of first-generation gradient-descent algorithms, substantial progress has been made in generalizing the method to handle non-convex surfaces, often by adding a “momentum” term. Momentum in SGD makes the system’s evolution a function of not only the current gradient, but also past gradients. Like physical momentum, this memory allows the algorithm to “roll past” small dips in the error surface. The NADAM algorithm is one of the most recent advances in momentum-based SGD ([34]). Rather than just updating the parameter estimate (*θ*_*k*_) at each time step, NADAM also updates a moving average of the gradient (*m*_*k*_) and the squared gradient (*n*_*k*_). The moving average of the gradient adds momentum, while the moving average of the squared gradient is used to adaptively scale updates according to the mean square error. The memory of the moving average gradients and squared gradients are controlled by the hyperparameters *μ* and *ν*, respectively. A “regularization” hyperparameter (*∊*) stabilizes the learning rate and prevents division by zero. The NADAM algorithm thus updates as follows:

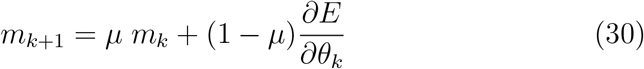

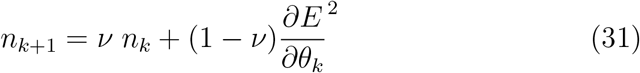

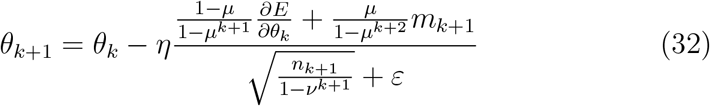

**Fig 10.**
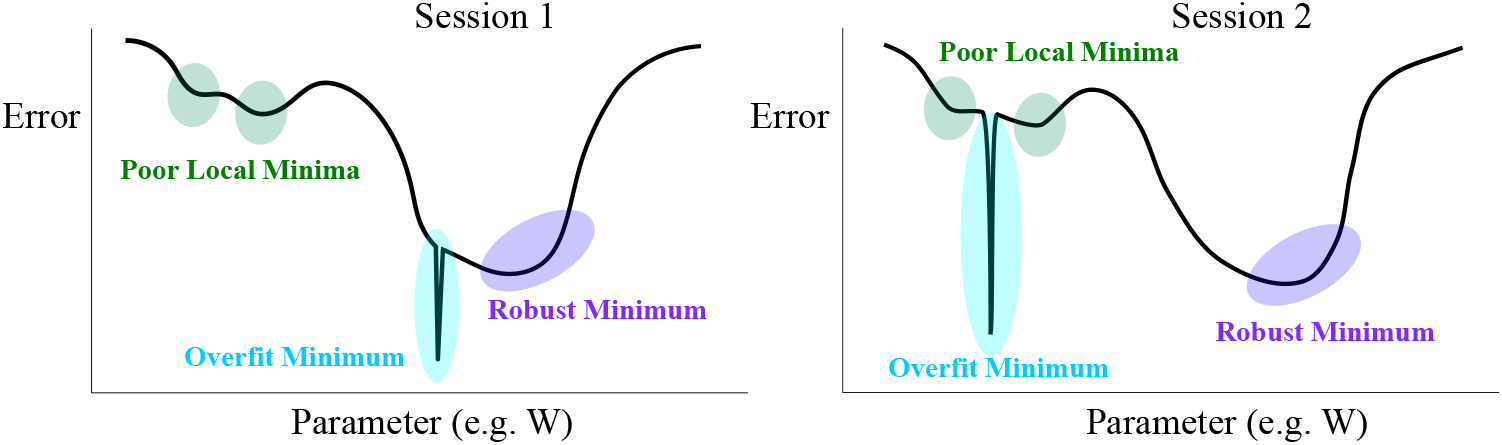
Schematic of NADAM benefits with illustrative error surfaces (y-axis) for fitting a parameter (x-axis values) on the first scanning session (left) and the second (right). The NADAM algorithm uses momentum to avoid shallow local minima (green). This feature also prevents convergence to overly sharp minima (even if they are global) because such error surfaces can often correspond to overfitting (blue) and hence do not generalize across sessions. Rather, NADAM emphasizes solutions to deep basins (purple) which may prove the most robust.

Like its predecessor, the ADAM algorithm ([85]), NADAM makes use of momentum to avoid converging to shallow minima and also incorporates estimates of the error surface curvature ([34]). However, like all SGD methods, the NADAM algorithm is still only guaranteed to converge to a local minimum. The advantage, however, is that the NADAM algorithm improves the depth and breadth of that local minimum (10). Due to the limited amount of data per subject we prioritize robustness over goodness-of-fit so the global-minimum is not necessarily desirable and might actually correspond to over-fitting. There are thus two main advantages to using modified SGD over a global-optimizer: 1) computational efficiency, which enables us to fit very large networks, and 2) emphasis on robust solutions, which improves cross-validation and prevents over-fitting.

### 5.4. Hyperparameters in Model Fitting

In deconstructing the connectivity matrix, we produce three terms: one *n × n* sparse component (*W*_*S*_) and one *n × m* rectangular matrix for each of the two diffuse components 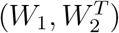 in which *n* denotes the number of parcels and *m < n* denotes the chosen dimensionality of the diffuse matrix. Hence, 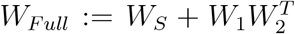. The sparsity of *W*_*S*_ is achieved with *L*_1_ regularization with penalty *λ*_1_ and both of the diffuse components are also *L*_1_ penalized with the same coefficient (*λ*_2_) for both halves. The full diffuse matrix 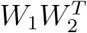 also receives *L*_2_ regularization.

The full integrated cost function which includes the regularization penalty is thus:

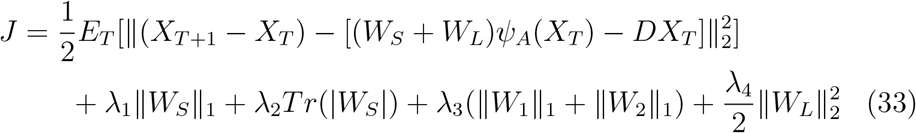

with the notation *E*_*T*_ denoting the expected value over all time points within the minibatch. The NADAM algorithm itself involves four parameters: an update rate parameter, two decay parameters for computing moving averages, and one “regularization” parameter ([34]). Unlike the regularization parameters for the weight matrices, which factor into the error and steer the model towards sparse solutions, the NADAM regularization parameter simply serves to stabilize the speed of updates and prevent division by zero. We chose parameters for each variable: *W*_*S*_, *W*_*L*_, *α, D*. As with the regularization terms, we used the same parameters for the two halves of the diffuse component: *W*_1_ and *W*_2_. We found that the least impactful hyperparameters are the NADAM decay rate hyperparameters, which only need to be slightly less than one. The most impactful hyperparameters are the *L*_1_ regularization penalties for the weight matrices which control the balance between over-fitting and under-fitting.

### 5.5. Interpretations of the Weight-Decomposition and Well-Posedness of the Problem

From a Bayesian perspective, this penalty function is equivalent to maximum a posteriori (MAP) estimation with fixed Laplace distribution (symmetric exponential) priors for each of the individual weight matrices and a normal prior on the combined low-rank component. The Laplace (distribution) prior is unrelated to Laplace approximation as used in Bayesian estimation. Since we assume that process noise is iid. between parcels, its influence (scaling the prediction error term) gets absorbed in the regularization coefficients (by multiplying all terms of the log-likelihood by the noise variance). From a linear-algebra perspective, the regularization prioritizes matrices which can be minimally perturbed to produce a skewed eigenvalue spectrum with sparse eigenvectors. It is important to note that the sum of sparse and low-rank matrices (e.g. *W*) need not be sparse nor low-rank so this decomposition is quite flexible. The values of each *λ*_*i*_ and their determination is detailed in SI Table 13 and SI Sec. 5.4.

The primary function of this decomposition is to prevent over-fitting. In all contexts, the potential for over-fitting is related to the difference of model and data degrees of freedom (parameters vs. measurements). While the scale of the current MINDy model might induce initial skepticism, the current problem is similarly well-posed as several recent attempts at fitting a smaller number of parameters. For instance, a recent approach by Wang and colleagues ([13]) to fit just local parameters (2 per node) in modeling the functional connectivity matrix results in 138 parameters estimated from 2,278 data points collapsed across the whole brain (approximately 16.5 measurements per parameter). Although the current approach estimates far more parameters (n+2 per node) it also utilizes many more data points: the whole multivariate time series is used for estimation rather than just the functional connectivity matrix. This results in a ratio of roughly 12 measurements per parameter using the full HCP resting-state data for subjects with MMP cortical and Freesurfer subcortical parcellation (11 for the gwMRF-400). The HCP dataset also contains double this quantity for a subset of subjects who also participated in a later retest session. Of course, these “back-of-the-envelope” calculations assume the worst-case scenario of no parameter covariation. In reality, we expect the set of underlying effective connectivity matrices to be much more constrained—a fact that we exploit via our weight decomposition. Moreover, the sparse regularization priors result in many weights becoming negligible (i.e. very near zero), so even fewer non-trivial estimates are made (Fig. 5A,B). The fitting process is also tractable due to the use of the NADAM algorithm ([34]) which is optimized for simultaneously fitting very large numbers of parameters in a highly efficient manner (approximately one minute per model on a latop).

### 5.6. Reliability and Individual Differences in the Weight Decomposition

In the main text (Sec. 2.2), we introduced a linear decomposition of the weight matrix into sparse and low rank components:

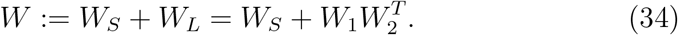

This decomposition was motivated by the dual influences of sparse, long-distance connections between “hub” regions (which motivates *W*_*S*_) and the propagation of these signals along subnetworks (which motivates *W*_*L*_). This formulation was developed as a fitting heuristic by which we could approach the high-dimensional model-estimation inherent in MINDy (Eq. 4). All previous analyses have focused upon the final weight matrix *W*, since the dynamical systems models in MINDy do not require explicit consideration of the components (*W*_*S*_ and *W*_*L*_). In this section, we present preliminary analyses which suggests reliable individual differences in this decomposition. We do not separately analyze the two rectangular matrices *W*_1_ and *W*_2_ which define *W*_*L*_ since they are not unique (e.g. their column indices are arbitrary). As with the full weight-matrix, we measured the degree to which estimates were similar within-subject (different sessions) vs. between-subject. We only consider non-diagonal elements of the matrices since the recurrent elements (diagonals) are distinguished by a separate regularization term in the cost function (Eq. 4). The results for recurrent connections in isolation were: within-subject: *r* = .56±.06, between-subject: *r* = .42 ± .07. We found that both components had greater similarity within-subject (*W*_*S*_ : *r* = .53 ± .06 and *W*_*L*_ : *r* = .67 ± .04) than between-subjects (*W*_*S*_ : *r* = .35 ± .03 and *W*_*L*_ : *r* = .39 ± .04). Thus, the component matrices had significant reliability and exhibited individual differences although the reliabilities were lower than that of the full weight matrix (see Sec. 3.3).

We also considered the degree to which sparsity vs. low-dimensionality of a subject’s weight matrix was a reliable trait. We quantified this value by comparing the (log) relative magnitude of the sparse and low-rank matrices (*U*):

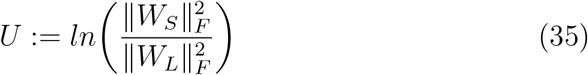

With 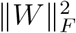 denoting the squared Frobenius-norm (sum of squared matrix elements). Each model produces a single (scalar) value for *U* . We tested whether individual differences in this quantity were reliable either with or without including recurrent connections. We found that results were reliable in either case (with: *ICC* = .763, without: *ICC* = .710) and that individual subject’s values were highly correlated for the two cases (*r*(51) = .919 for the mean across sessions). When recurrent connections are included, the sparse component is larger (i.e. *U* > 0) on average: *U* = .265 ± .261 while the low-rank component otherwise dominates: *U* = − .528 ± .215. Thus, results indicate that the relative magnitude of sparse vs. low-rank components is a reliable marker of individual differences in MINDy models. However, the degree to which the overall weight matrix is dominated by either component will depend upon whether the recurrent (sparse) connections are considered. Moreover, we expect that the average value of this ratio will depend upon the particular choice of regularization parameters which will favor either component. Therefore, while results are promising in terms of individual differences, we do not recommend using the weight decomposition to quantify the general sparseness/dimensionality of brain networks without considering the influence of regularization hyperparameters.

**Fig 11.**
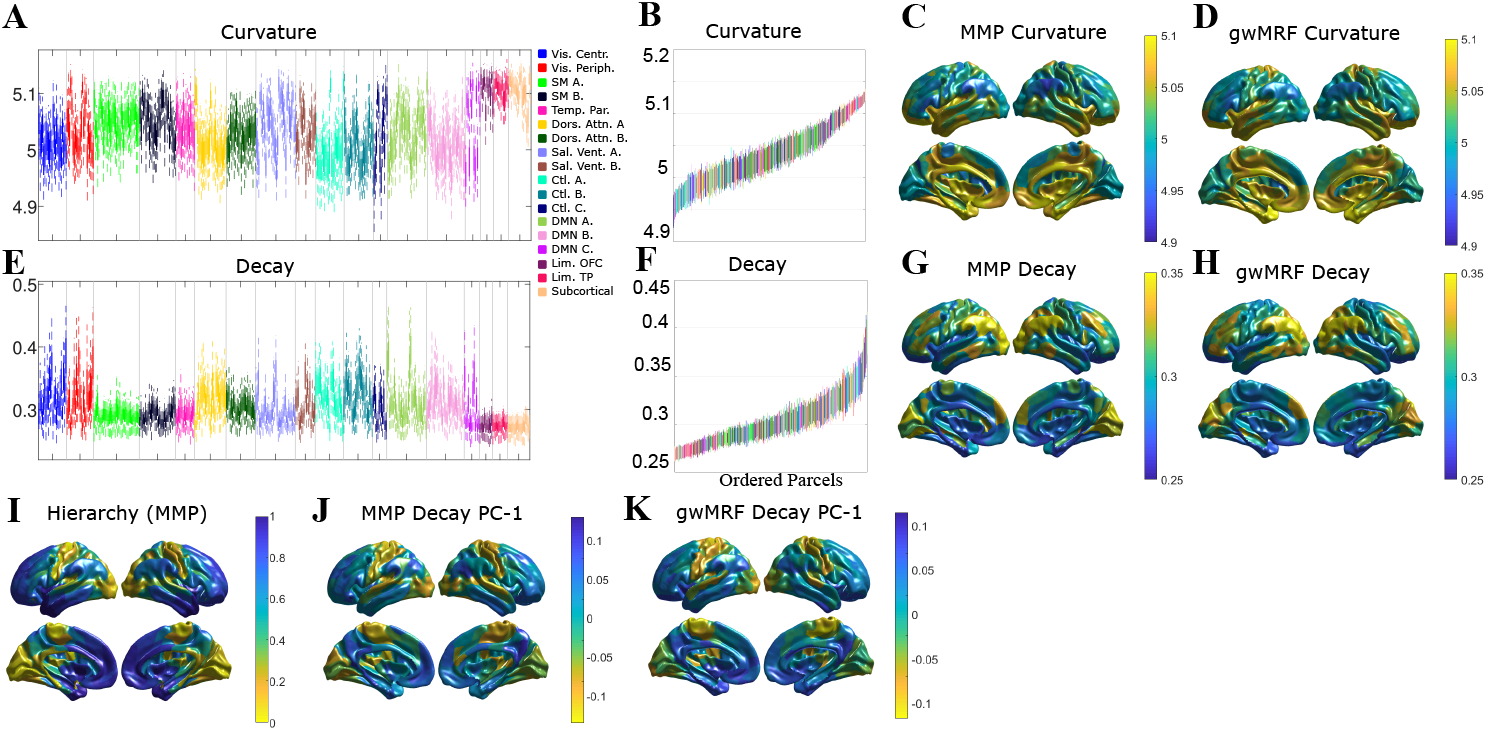
Local MINDy parameters (curvature and decay) exhibit consist anatomical structure within and between networks. A) Distribution of curvature parameter values for each brain parcel grouped according to network (17-network [21]). B) Curvature parameters reordered according to mean demonstrate that within-network variability is also consistent. C) Anatomical profile of group-mean curvature for the MMP atlas ([20]). D) Profile for the gwMRF ([21]) parcellation. E-H) same as A-D but for the decay parameter. I) Hierarchical Heterogeneity map by Demirtas and colleagues ([12]) using group T1/T2 ratio. J,K) same as G,H but for the first principal component of decay across subjects.

### 5.7. The Influence of Hyperparameter Choices on Sparsity

The relative value of the hyperparameters *λ*_1_ vs. *λ*_2_ can influence the sparsity of the MINDy connection matrix. Overly small values of *λ*_1_ will not generate sparsity in the “sparse” matrix (*W*_*S*_) while overly large values of *λ*_1_ relative *λ*_2_ will also decrease sparsity by biasing solution coefficients toward the low-rank component (*W*_*L*_). However, in practice, even the low-rank matrix is substantially sparser than the rsFC which generates overdispersion (“heavy-tailedness”). We quantify this property via kurtosis:

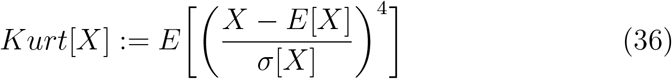

which, for a normal distribution, is 3. The kurtosis for each subject’s low-rank component (35.6 ± 8.1) substantially exceeds that of the rsFC matrix (7.2 ± 0.9) although both are dwarfed by the sparse component (188.7 ± 27.8). Thus, both components of the weight matrix are more sparse than *rsFC* so hyperparameter choices which bias towards either term will still result in sparser solutions than rsFC. Lastly we considered the case in which all regularization terms are equal zero which forms a lower-bound case for sparsity (i.e. all other regularization values should produce more sparse estimates). Individual model estimates without regularization are extremely noisy (hence the need for regularization) and less sparse (*Kurt* = 5.9 ± 0.7) than rsFC. However, the group-mean of these noisy estimates (*Kurt* = 24.4) is also more sparse than the group-mean for rsFC (*Kurt* = 9.0). Moreover, the group-mean without regularization was highly correlated (*r*(175559) = .917) with the group-mean for the full MINDy model. We conclude that individual weight estimates are sparse for a range of hyperparameter values and the group mean of estimated weights for our dataset is more sparse than rsFC, irrespective of hyperparameter choices.

### 5.8. Comparing MINDy and Spectral Dynamic Causal Modeling

We view the primary contributions of MINDy in its scalability, biological interpretability, and the ability to predict nonstationary resting-state dynamics. However, one non-unique benefit of MINDy is the data-driven characterization of effective connectivity via the weight parameter. Other methods, such as stochastic DCM and spectral DCM have also used (linear) dynamical systems models to estimate effective connectivity. By converting problems into the frequency domain, spectral DCM (spDCM; [7]) has been applied to brain models consisting of 36 regions, but still has significantly higher computational cost than MINDy. One question, therefore, is whether the scalability of MINDy comes at the cost of accuracy. We tested this question in a series of ground-truth simulations. To be clear, we are not seeking to demonstrate that MINDy is necessarily a better estimator of low-dimensional effective connectivity, but rather that the scalability of MINDy does not significantly impair accuracy (i.e. MINDy is at least as good as DCM). One inherent advantage of spectral DCM is the ability to estimate region-specific hemodynamic kernels which is not part of the currently proposed MINDy model (although extensions for HRF estimation are being developed [63]). Thus, we consider two features when comparing MINDy and spectral DCM (spDCM): scalability and robustness to spatial variation in the HRF.

#### 5.8.1. Benchmarking with Unbiased Ground-truths

The main difficulty in comparing MINDy and DCM is the different underlying assumptions—spDCM, for instance, has only been validated using a linear ground-truth ([7]). We took a number of steps to prevent bias based upon differing assumptions (SI Tab.1) and when bias was inevitable, we made choices that favored spDCM. First, we used two ground-truths which were not based upon either technique: a neural mass + balloon-Windkessel ground-truth and a continuous asymmetric Hopfield-model ground truth with either a random global HRF or spatially variable hemodynamics (the same models as for model-mismatch analyses in Sec. 3.7.5). Random HRFs were generated by sampling 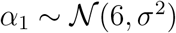 and 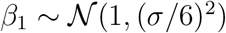 in Eq. 5. For the global HRF simulations, *σ* = .5 and the same values of *α*_1_ and *β*_1_ were used for each node (for a given simulation). In the spatially-variable HRF simulations, values were independently drawn for each node. We implemented spectral-DCM using the MATLAB code provided with SPM-12 (function name: “spm_dcm_fmri_csd”) to compute the expected value for each connection weight based upon cross-spectral density (spectral DCM). Simulations and model-fitting were performed single-core on Intel Xeon E5-2630v3 CPUs. Since linear models like DCM do not separate recurrent connections and decay, we only compared accuracy for non-recurrent (off-diagonal) elements of the connectivity matrix for each technique. Both ground-truth simulations were integrated at time-scales faster than the sampling rate (dt=.025 vs. 725ms TR for the neural mass and dt=100ms vs. 700ms TR for Hopfield to mirror HCP acquisitions). This feature ensures that results are not biased against spDCM due to simulation time-scale, since MINDy discretizes the model in terms of TR, while spDCM maintains a continuous-time estimation framework.

**Fig 12.**
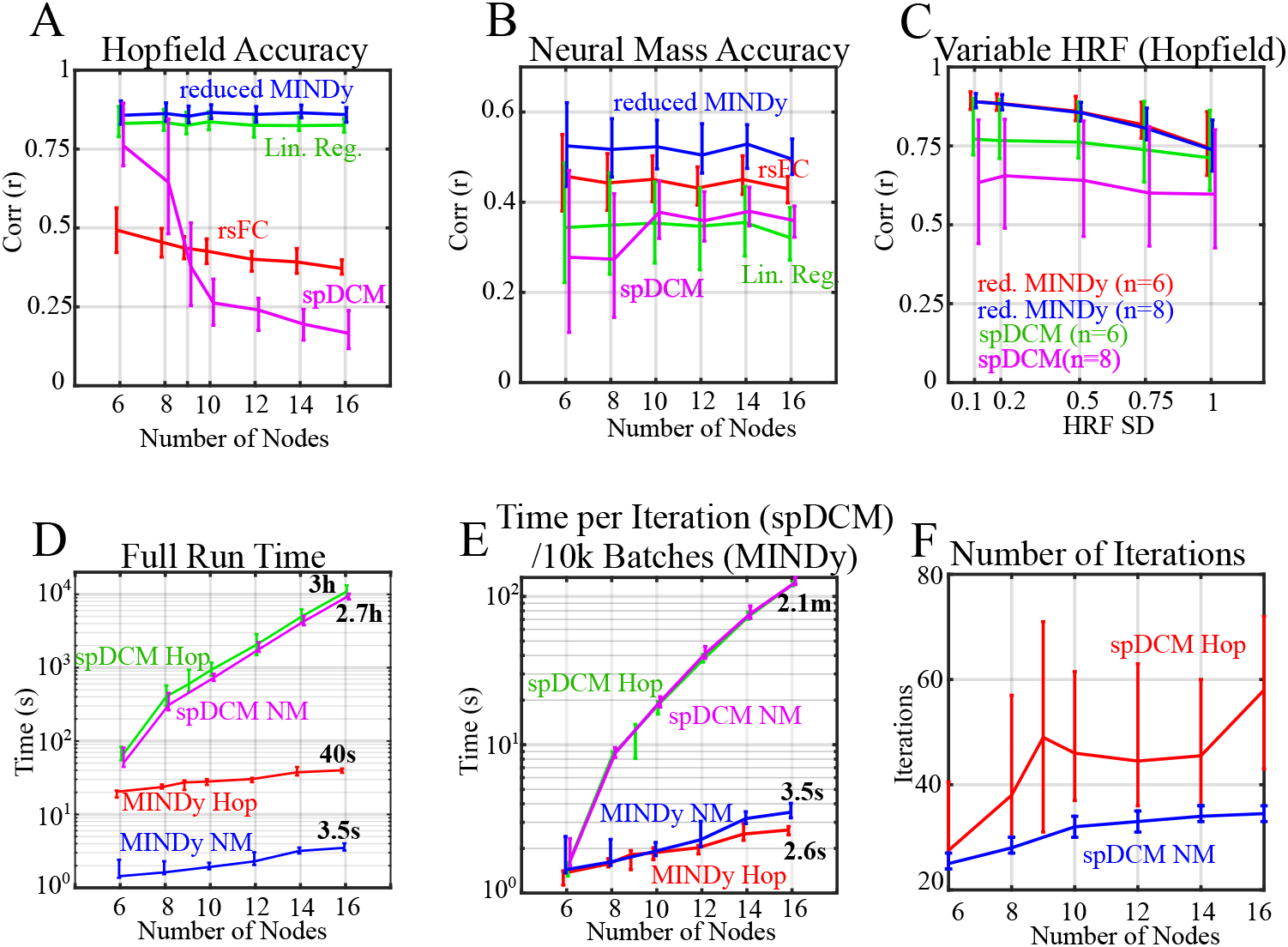
Comparison of accuracy and run-time for MINDy and spDCM. A) Accuracy in estimating ground-truth connectivity from Hopfield-network simulations by network size. “Reduced MINDy” indicates that all regularization terms were removed from MINDy to avoid bias (analogous results for the full MINDy model are in Sec. 3.7.5). Lines indicate mean and bars indicate first/third. quartiles. B) Accuracy in estimating ground-truth connectivity from Neural Mass simulations by network size. Results for the full MINDy model are in Sec. 3.7.5. C) Model performance as a function of HRF spatial variability for two network sizes: 6 and 8 nodes. Note that MINDy performance decreases with HRF spatial variability, whereas the effect for spDCM is minor. D) Full run times for MINDy and spDCM for each simulation as a function of simulation type/size. E) Run time per EM iteration (spDCM) and for 10,000 mini-batches in MINDy. We chose to compare with 10,000 mini-batches so that the run-times would be comparable for the smallest network size (*n*_*Pop*_=6). F) Model performance as a function of HRF spatial variability for two network sizes: 6 and 8 nodes. Note that MINDy performance decreases with HRF spatial variability, whereas the effect for spDCM is minor.

We used the same hyper-distributions to parameterize neural-mass simulations and Hopfield networks as in Section 3.7.5, but for smaller network sizes. We made three further adjustments to reduce bias: first, we set all regularization terms from MINDy equal to zero, so there would be no inherent advantage to MINDy based upon the hyper-distributions of network structure. The regularization-based calculations were still performed (preserved run time), but they had no effect on the solution (were always zero). We also increased the simulation length of the neural-mass simulation from 3000 to 5000 TRs and removed the nearest-neighbor smoothing from the Hopfield Network simulations which had been included to mirror the empirical processing-pipeline. These adjustments did not affect MINDy but were found to increase the performance of spDCM.

In addition to MINDy and spDCM, we tested the accuracy of functional connectivity and one-step prediction by multiple-regression (solving Δ*x* = *Mx*_*t*_). The latter case provides an additional control by providing an alternative method to parameterize linear dynamical systems. This control is important because it can indicate that cases in which spDCM underperforms are due to the estimation technique rather than linearity per se. (i.e. cases in which multiple-regression is accurate but spDCM is not). When all regularization terms are set equal to zero and the transfer function is linear, the MINDy and multiple-regression models are equivalent. The regression approach differs from spDCM, however, in the strength of the linearity assumption. Whereas spDCM seeks a linear model that best explains statistics drawn from the full time-course (i.e. assumes global linear dynamics), the regression approach (like MINDy) considers the changes at each TR (i.e. the collection of local dynamics).

#### 5.8.2. MINDy Performs Competitively with DCM

Results indicated that MINDy scaled-well in terms of performance and run-time. Moreover, this scalability did not generally come at a cost to performance relative contemporary higher-complexity techniques (i.e. spectral DCM). In all situations tested (model x size x HRF), the reduced MINDy model (no regularization) performed at least as well as all competitors in retrieving ground-truth connection weights (SI Fig. 12 A-C). All methods performed poorer for the neural-mass ground-truth (SI Fig. 12B; SI Tab. 2) than for the Hopfield network (SI Fig. 12A; SI Tab. 3; SI Tab. 4) which was expected due to the greater difficulty of the problem (much faster time-scales and more complicated models). We note that the full MINDy model performs substantially better in neural mass simulations than the reduced version (see Sec. 3.7.5). However, even when regularization was removed to prevent bias due to assumptions on network structure (“reduced” MINDy), performance remained competitive with spDCM. We also observed that spDCM performance decreased with network size for the Hopfield simulation (SI Fig. 12B; SI Tab. 3), but the regression-based model did not. Thus, this feature cannot be explained solely in terms of functional form (linearity) although the reduced MINDy did consistently outperform regression.

Moreover, the computational complexity of MINDy is substantially lower than that of spDCM. The theoretical limiting computational complexity of MINDy is a second-degree polynomial in the number of nodes since the highest-complexity operation in terms of *n*_*Pop*_ is analytically calculating the error-gradient with respect to the weight-matrix 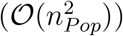 although the empirical complexity was substantially lower for these simulations (the quadratic term only dominates for much larger *n*_*Pop*_; SI Fig. 12D). By contrast, the spectral-DCM code packaged with SPM has at least fourth-order complexity in terms of the population size for a fully-connected model (SI Fig. 12D). Each Expectation-Maximization (EM) iteration is dominated by 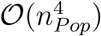 (SI Fig. 12E), but the total complexity can be even greater if the number of iterations until convergence also increases with *n*_*Pop*_ (SI Fig. 12F). For instance, the median total runtime for the neural mass and Hopfield network simulations scaled with 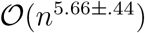 and 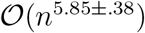, respectively (95% confidence estimated using the “fit” function in MATLAB 2020a for linear power functions: “power1”). The empirical complexity of each EM iteration was roughly the theoretical limit of 4: 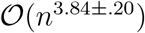 and 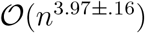 for the neural mass and Hopfield simulations, respectively.

However, even the minimal case of fourth-order complexity can severely limit scalability. For context, a recent “large-scale” spectral DCM paper ([7]) employed 36 brain ROIs with a run-time of between 1,280 and 2,560 minutes per model (between 64 and 128 iterations at 20 minutes each). Increasing resolution from 36 ROIs to the 419-node parcellation we employed (19 subcortical + 400 cortical [21]) would increase CPU time by a factor of over 18,000 (roughly 44 to 89 years per model for the same data and hardware as [7]). By contrast, the current MINDy models for HCP data were locally fit on a laptop in less than one minute each (Intel i7-8750H CPU, 2.2GHz, 6 cores).

**Table 1.**
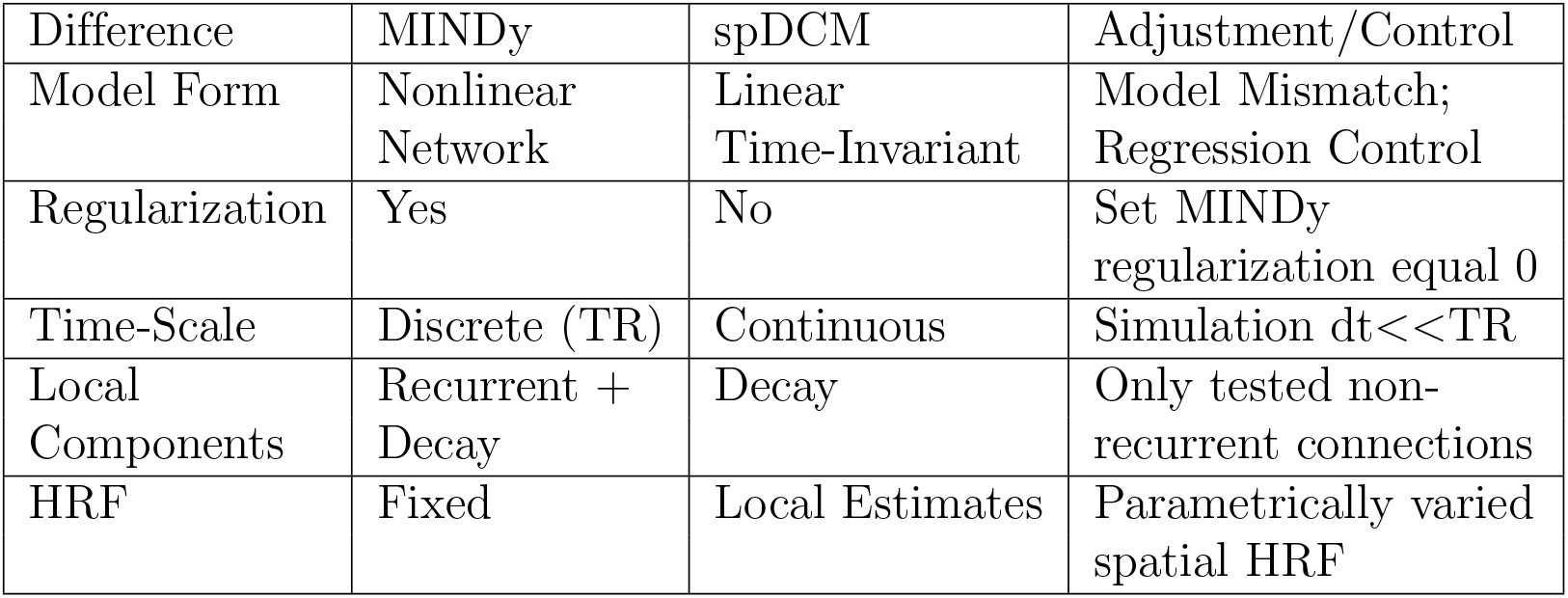
Types of assumptions (“Difference”) made by MINDy vs. spDCM and our controls to mitigate these differences in simulated comparisons (SI Sec.5.8).

One area in which spectral DCM proved advantageous, however, was in robustness to spatial variability in the hemodynamic response (SI Fig. 12.C; SI Tab. 4). As with the analogous simulations in Section 3.7.5, MINDy performance decreased with the underlying HRF’s spatial variability (see also [63]). This pattern is expected since the currently proposed MINDy assumes a fixed HRF which these simulations violate. The performance benefit of MINDy over spDCM, likewise decreased with HRF spatial variability. For the smallest network (6 nodes) and highest level of HRF variability considered, the difference between models’ accuracy was negligible (although statistically significant; SI Tab. 4). Thus, there may be cases of extreme HRF spatial variability in which spectral DCM outperforms MINDy for sufficiently small networks. Although MINDy scales significantly better than competing approaches like spectral DCM, the current version is less robust to spatial variability in the HRF (although see [63] for upcoming extensions). Nonetheless, for all simulations considered, MINDy performed competitively with spDCM within the latter’s scope. We conclude that MINDy performs at least as well as spDCM, while scaling far better.

### 5.9. Anatomical Distribution of Individual Differences

In concert with the previous analyses of individual differences (Sec. 3.5) across parameter-types (weights, curvature, decay) we also investigated the anatomical distribution of individual differences within each parameter-type. These analyses are post-hoc (exploratory) so we report results as a potential launching pad for future investigations and as a means to understand how MINDy models encode individual differences. We do not perform hypothesis-testing and we caution against interpreting these analyses as stand-alone findings due to their exploratory nature and relatively low sample-size. We also note that these analyses are performed upon a biased sample of subjects—those that had no high-motion scans (>1/3 frames censored) so these results may also fail to describe variability in the full HCP subject pool (see [28] for cognitive covariates of motion) and its target population (American young adults). These caveats aside, we considered the degree of individual variation for each parcel/connection. We used the quartile coefficient of dispersion (QCD) to quantify the degree of variability within parcels/connections. Conceptually, the QCD is a robust analogue of the more commonly used Coefficient of Variation and is defined in terms of the first (*Q*_1_) and third (*Q*_3_) data quartiles as:

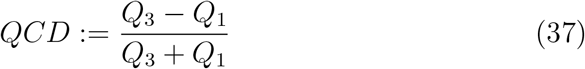

**Table 2.** Accuracy (*r*) for each method (Regr.=regression) for the neural-mass simulation (SI Sec. 5.8) with variable population sizes (left side). Test-statistics (paired t-test; 2-tailed) for MINDy-spDCM are provided on the right side. The number of nodes per simulation is listed under “nodes” whereas the number of simulation instances is listed under “N”. The paired difference in accuracy between MINDy and spDCM is denoted Δ*r*. We denote ∗ = *p* < .01 and ∗∗ = *p* < .001.

| Nodes | rsFC | Regr. | MINDy | spDCM | N | $\Delta r$ | t |
| --- | --- | --- | --- | --- | --- | --- | --- |
| 6 | .46(.14) | .34(.20) | .52(.15) | .28(.24) | 240 | .25(.26) | 14.8** |
| 8 | .44(.10) | .35(.16) | .52(.11) | .27(.20) | 320 | .24(.21) | 20.4** |
| 10 | .45(.08) | .35(.12) | .52(.09) | .38(.10) | 130 | .15(.10) | 17.3** |
| 12 | .43(.07) | .35(.11) | .50(.09) | .36(.09) | 80 | .15(.08) | 16.6** |
| 14 | .45(.06) | .35(.09) | .53(.07) | .38(.07) | 40 | .15(.06) | 14.5** |
| 16 | .43(.05) | .32(.09) | .50(.08) | .36(.06) | 20 | .14(.03) | 22.8** |

**Table 3.**
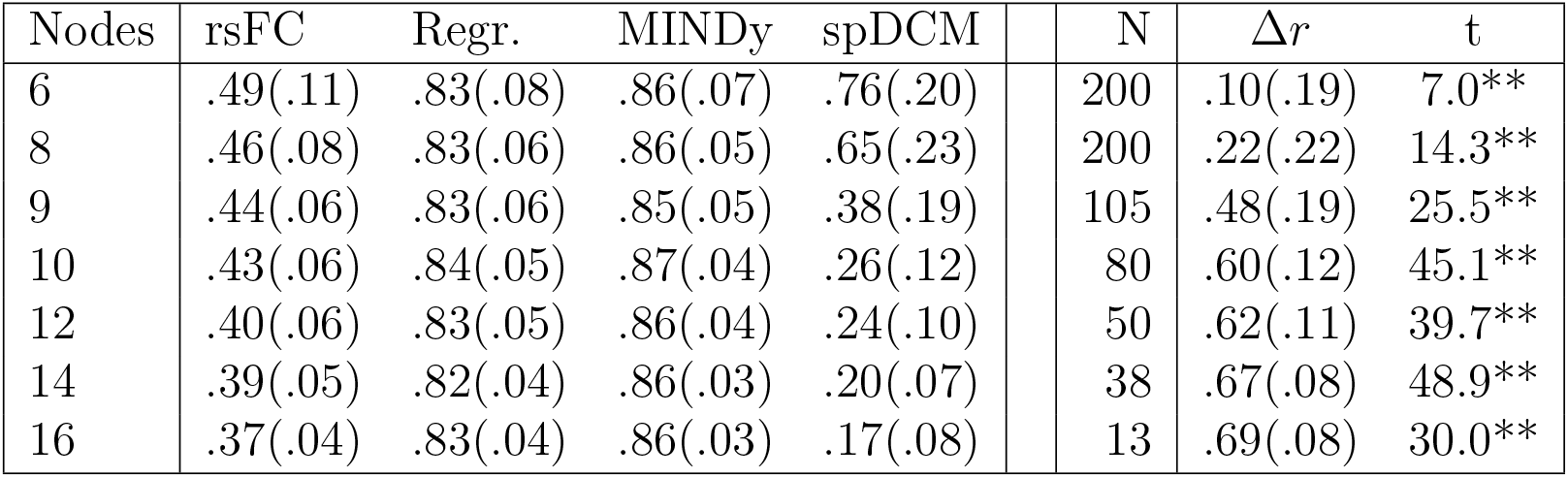
Accuracy (*r*) for each method (Regr.=regression) for the Hopfield-network simulation (SI Sec. 5.8) with variable population sizes (left side) and a random global HRF (*σ* = .5). For this simulation we added a new population size (9 Nodes) post-hoc to see whether the unexpected decrease in spDCM accuracy between 8 and 10 nodes was continuous (it was). Since this special case contained an odd number of nodes the hyperdistribution parameter *q* in Sec. 2.6.2 was always equal to one (instead of one and two with equal probability). Test-statistics (paired t-test; 2-tailed) for MINDy-spDCM are provided on the right side. The number of nodes per simulation is listed under “nodes” whereas the number of simulation instances is listed under “N”. The paired difference in accuracy between MINDy and spDCM is denoted Δ*r*. We denote ∗ = *p* < .01 and ∗∗ = *p* < .001.

| Nodes | rsFC | Regr. | MINDy | spDCM | N | $\Delta r$ | t |
| --- | --- | --- | --- | --- | --- | --- | --- |
| 6 | .49(.11) | .83(.08) | .86(.07) | .76(.20) | 200 | .10(.19) | 7.0** |
| 8 | .46(.08) | .83(.06) | .86(.05) | .65(.23) | 200 | .22(.22) | 14.3** |
| 9 | .44(.06) | .83(.06) | .85(.05) | .38(.19) | 105 | .48(.19) | 25.5** |
| 10 | .43(.06) | .84(.05) | .87(.04) | .26(.12) | 80 | .60(.12) | 45.1** |
| 12 | .40(.06) | .83(.05) | .86(.04) | .24(.10) | 50 | .62(.11) | 39.7** |
| 14 | .39(.05) | .82(.04) | .86(.03) | .20(.07) | 38 | .67(.08) | 48.9** |
| 16 | .37(.04) | .83(.04) | .86(.03) | .17(.08) | 13 | .69(.08) | 30.0** |

**Table 4.** Accuracy (*r*) for each method (Regr.=regression) for the Hopfield-network simulation (SI Sec. 5.8) with spatially variable HRF and 6 nodes (left side). Test-statistics (paired t-test; 2-tailed) for MINDy-spDCM are provided on the right side. The standard-deviation of the HRF parameters is listed under “*σ*” whereas the number of simulation instances is listed under “N”. The paired difference in accuracy between MINDy and spDCM is denoted Δ*r*. We denote ∗ = *p* < .01 and ∗∗ = *p* < .001.

| σ-HRF | rsFC | Regr. | MINDy | spDCM | N | $\Delta r$ | t |
| --- | --- | --- | --- | --- | --- | --- | --- |
| .1 | .48(.10) | .87(.06) | .89(.04) | .77(.18) | 280 | .12(.17) | 11.8** |
| .2 | .48(.11) | .86(.06) | .89(.05) | .77(.19) | 280 | .12(.18) | 11.1** |
| .5 | .49(.12) | .83(.08) | .86(.07) | .76(.18) | 280 | .10(.18) | 9.3** |
| .75 | .47(.12) | .80(.10) | .82(.10) | .74(.20) | 280 | .08(.21) | 6.5** |
| 1.0 | .47(.11) | .72(.16) | .75(.15) | .71(.21) | 280 | .03(.21) | 2.7* |

**Fig 13.**
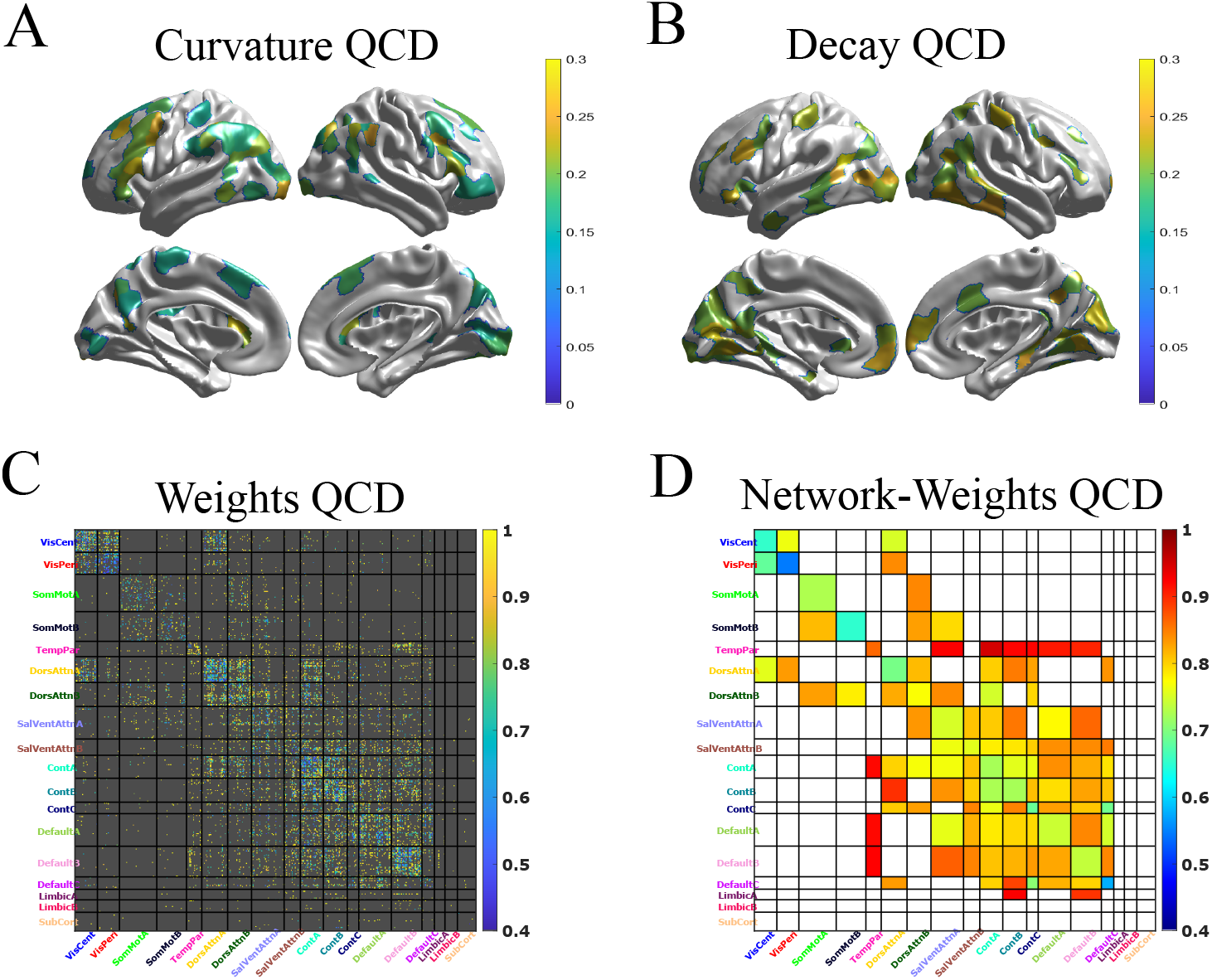
Anatomical distribution of inter-individual variation in MINDy. A) QCD of the curvature parameter with top-20% threshold. B) Same as A) but for the decay parameter. C) QCD for MINDy connection weights. Weights in which the sign was inconsistent across subjects (<75% agreement) or low reliability (Fisher ICC<.5) were censored (grey). D) Mean weight QCD within each network combinations. If over 95% of parcel-wise connections were censored, the network-level connection was also censored (white).

The reason that we apply QCD instead of Coefficient of Variation is that, while both assume the true population follows a ratio-scale (e.g. are one-sided) the QCD is robust to extreme values which, due to measurement error violate the ratio-scale assumption. We censored connections which were unreliable (Fisher’s ICC≤.5) or incompatible with QCD (*Q*_1_ and *Q*_3_ differed in sign). We then transformed the other variables onto admissible distributions (non-negative ratio scales) by shifting the curvature and decay parameters so that they had a minimum value of zero. Interestingly, the anatomical distribution of QCD appeared to differ between the curvature and decay parameters. The curvature had the highest QCD in parcels of inferior frontal gyrus, early visual cortex, and a large posterior section of frontal cortex (SI Fig. 13A). For the decay parameter, QCD was highest in visual regions and the bilateral portion of somatosensory cortex traditionally associated with hands (SI Fig. 13B). By contrast, connection weights had the lowest QCD for connections within the visual networks (especially peripheral visual; SI Fig. 13C,D). Some of the highest QCD connections involved the temporal-parietal network (inter and intra-network) and connections to the limbic system. Of course, as previously mentioned, these analyses are purely exploratory and should only be interpreted as an example of how MINDy separates the sources of individual differences (weights, curvature, decay) rather than as a basic neuroscientific result.

### 5.10. Directed Connectivity Identified by MINDy

The simplest way to characterize connection asymmetries is in terms of regions being sinks (input weights greater than output weights in absolute value) vs. sources (output weights greater than input weights in absolute value). For now we focus upon sources and do so separately for positive (SI Fig. 14 A) and negative connections (SI Fig. 14 B). For positive sources MINDy most strongly identifies inferior frontal gyrus (IFG), bilateral parieto-occipetal sulcus, and dorsal prefrontal cortex (SI Fig. 14 A). MINDy identifies the strongest excitatory source as a region of left IFG (see main text; SI Fig. 14 C) and identified bilateral IFG as the strongest negative sources with negative outward-biased connections primarily to components of the Default Mode Network (IPL and medial PFC) with a general contralateral bias (SI Fig. 14 D,E). The role of right IFG in inhibition is well-documented within neuroimaging (e.g. [86]) and lesion studies suggest an inhibitory role for left IFG as well ([87]). These results indicate that the asymmetries within MINDy weights are functionally interpretable. However, these initial findings only scratched the surface of possible analyses.

**Fig 14.**
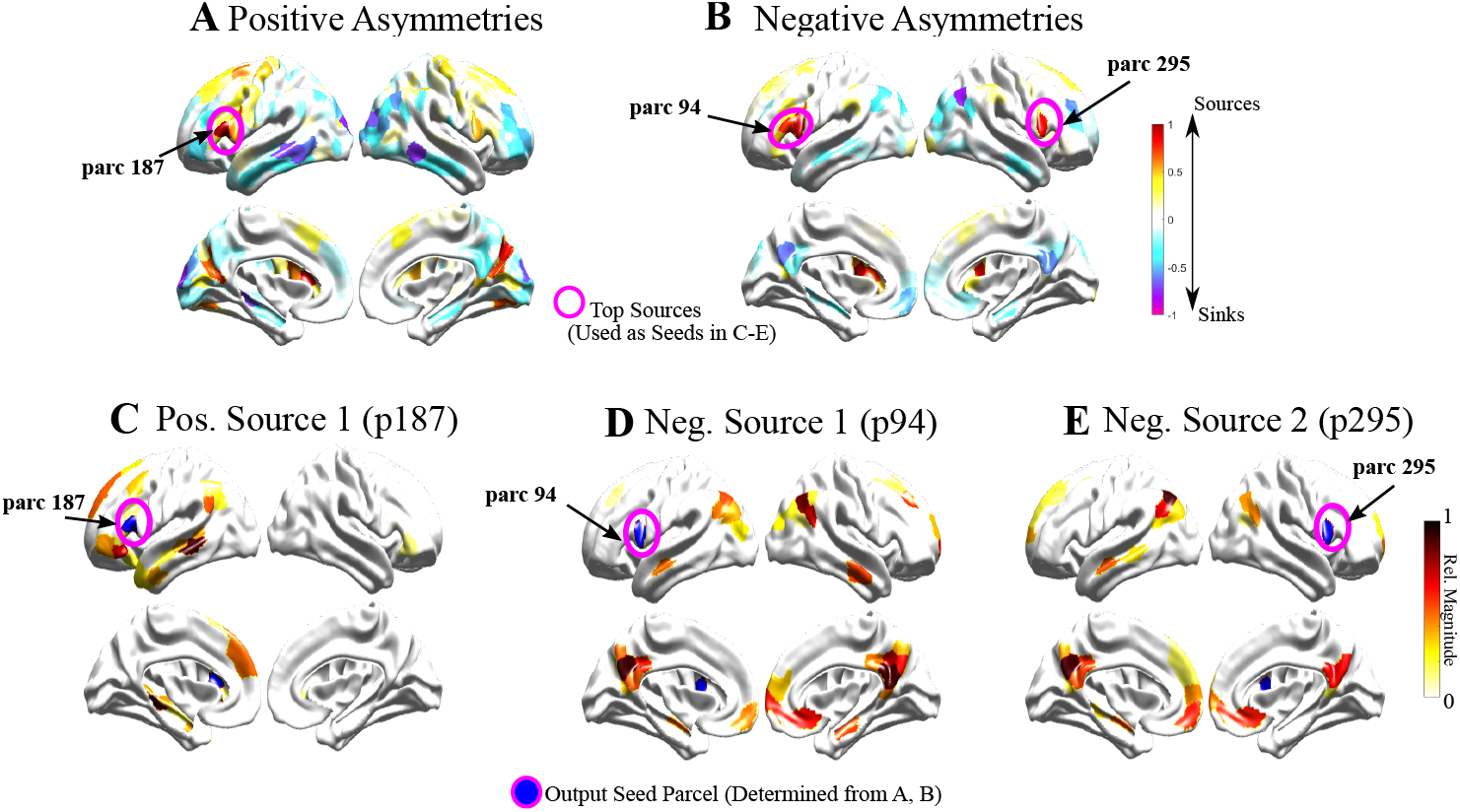
Connection asymmetries identified by MINDy A) Difference of total input weights minus output weights for positive connections only (normalized units). B) Same as (A) but for negative connections only (using difference in magnitude of input/output weights). C) Difference of output and input weights for positive output-biased connections in the parcel with the greatest positive output-bias. D) Same as C), but for the parcel with greatest negative output bias. E) The parcel with the second-greatest negative output bias is the contralateral analogue to the parcel in (D). Parcel numbers are labeled for the 17-network gwMRF parcellation ([21])

### 5.11. Nonlinear Dynamics in MINDy

As discussed in the main-text (Sec. 3.6.2), the fact that MINDy is nonlinear does not inherently imply that the model behaves qualitatively different from models relying upon a linear approximation (e.g. DCM). This distinction is critical to understanding how MINDy models resting-state dynamics: as random fluctuations about a single equilibrium (like DCM) or as topologically significant (nontrivial) dynamics. In general, proving the existence of global behavior in MINDy models constitutes a nontrivial endeavor given their high-dimensionality. However, we can easily rule out trivial dynamics (a Lyapunov-stable global attractor) by examining eigenvalues of the Jacobian at zero. Since all subjects contain at least one positive (real part) eigenvalue (SI Fig. 15), we conclude (by Proposition 2) that no empirically-parameterized MINDy model is globally Lyapunov-stable.

**Table 5.** Ground-truth validation performance of MINDy and rsFC in recovering the weight matrix of a single subject and the arithmetic difference of weight matrices between subjects. We denote significance with ** = *p* < .001, 2-tailed for the contrast Weights minus rsFC.

|  | Weights | FC | paired-t (df=33) |
| --- | --- | --- | --- |
| Single Subject | .800 (.025) | .436 (.065) | 45.629** |
| Indiv. Differences | .544 (.049) | .293 (.046) | 58.618** |

#### Proposition 2.

*Consider a continuous-time dynamical system evolving according to* 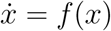, *with* 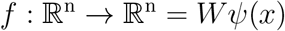 *with* 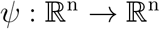 *an odd function (ψ*(−*x*) = −*ψ*(*x*)*) and n × n matrices W, D. Suppose that the Jacobian at the origin (F’*(0)*) has at least one eigenvalue with positive real part and none with zero real part. Then for any fixed point x*_*s*_ : *f*(*x*_*s*_) = 0, *at least one of the following hold*:

1. *There exists a set U of non-zero measure, whose positive limit-set ω*^+^(*U*) *does not contain x*_*s*_.
2. *x*_*s*_ *is not Lyapunov stable—i.e. there exists ∊* > 0 *for which there is no δ* > 0 *satisfying* ‖*x*(0) − *x*_*s*_‖ *< δ* ⇒ ∀*t* > 0, ‖*x*(*t*) − *x*_*s*_‖ < ∊.

*Proof.* We consider two cases depending upon whether *x*_*s*_ = 0. If *x*_*s*_ ≠ 0 is a fixed point then so is its reflection −*x*_*s*_ ≠ *x*_*s*_ since *f*(−*x*) = −*f*(*x*). Suppose that *x*_*s*_ is locally Lyapunov stable (violating implication 2) and thus possesses an attractive basin *V* of non-zero measure. Then −*V* is an attractive basin of −*x*_*s*_ hence *x*_*s*_ ∉ *ω*^+^(−*V*) which confirms the first implication. Thus, consider the alternative case: *x*_*s*_ = 0. By the hypothesis, *x*_*s*_ is a hyperbolic fixed-point so there exists an open neighborhood *N* containing *x*_*s*_ for which the dynamics on *N* are topologically conjugate those of the linearization 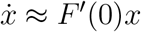 (Hartman-Grobman Theorem). The linearization *F’*(0) is unstable (at least one eigenvalue with positive real part) which implies that any sufficiently small ball about *x*_*s*_ = 0 will also be an unstable set. This contradicts Lyapunov stability (confirming implication 2).

### 5.12. MINDy Optimization: Under the Hood

To stabilize MINDy’s fitting procedure we use two changes of variable during the fitting process. Instead of directly fitting the term *D* we fit *D*_2_ satisfying the relation 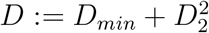. This change keeps the estimated parameters away from the pathological conditions in which *D* is either negative or very small which can cause models to explode in the long term. The term *D*_*min*_ is a constant, positive hyperparameter. This step does not significantly alter computational complexity and we found that it did not alter results for our current initialization setting of *D* as our estimates never approached the pathological regions. However, we included this change of variable in the code as a safeguard should it become relevant for future users.

**Fig 15.**
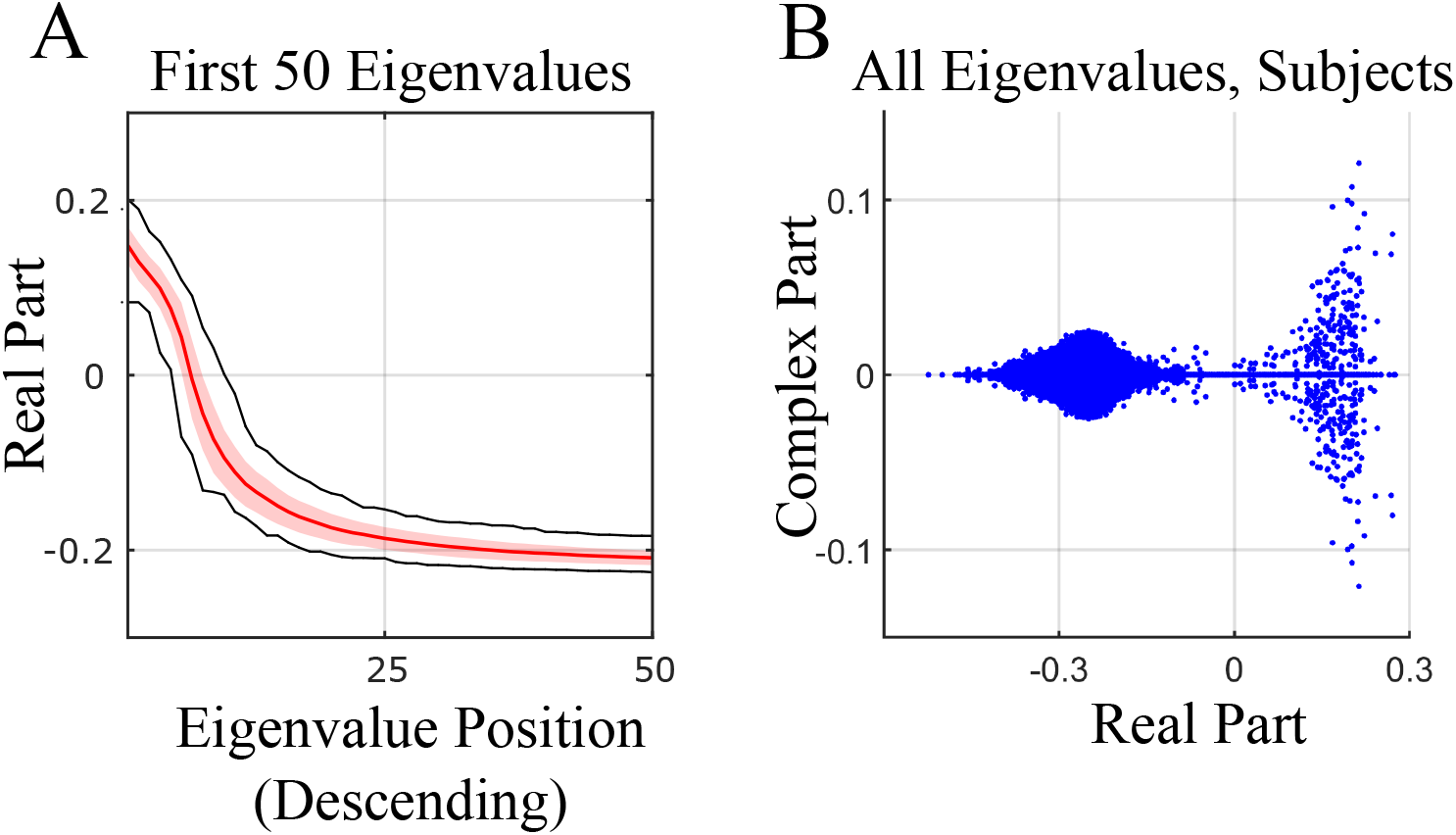
Eigenvalue analyses indicate that empirical MINDy models do not possess a global, Lyapunov stable attractor. A) Distribution of eigenvalues (real-part) for local-linearization about the origin. Shading indicates ±SD and black lines give the data’s maximum/minimum. Note the presence of positive eigenvalues which indicate nontrivial dynamics (Proposition 2). B) Scatterplot of these eigenvalues (all subjects) in the complex plane suggests that the greatest “spin” (complex components) occurs along the unstable subspaces (corresponding to eigenvalues of positive real-part).

**Fig 16.**
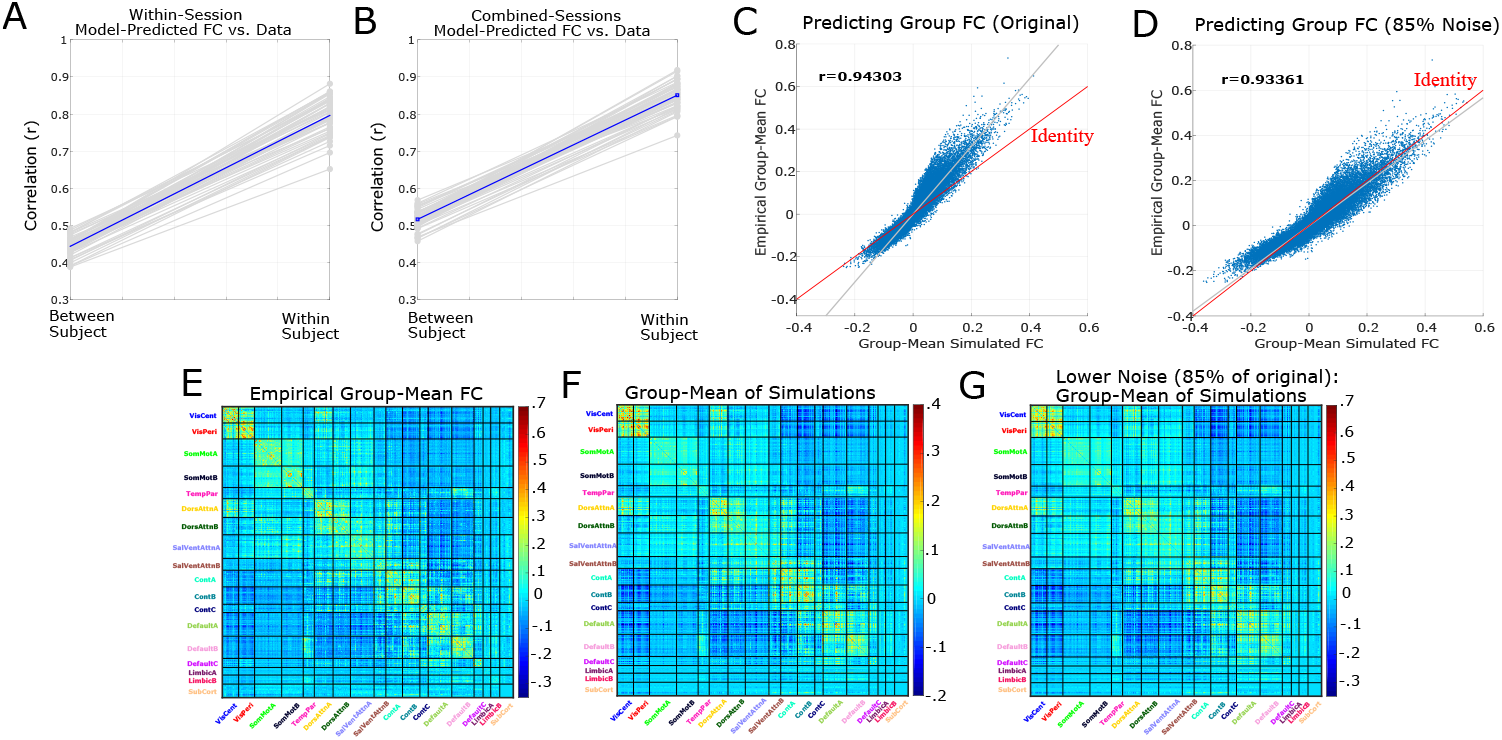
MINDy strongly predicts individual and group FC. A) Correlation between empirical FC and simulated FC from the same scanning day for either the same subject or a different subject (mean across other subjects). Blue line indicates group mean. Correlations are averaged across scanning sessions. B) Same as A) but for with all data combined across sessions (simulations from each session’s model were combined) C) Group-mean of empirical FC vs. group-mean of simulated FC (both combined across sessions). Notice that while the correlation is high, the magnitude of simulated FC is smaller than empirical. D) Slightly decreasing noise produces a predict group-mean FC that is nearly indistinguishable from that observed empirically. E) Group-average empirical FC combined across sessions. F) Group-average FC of simulated data combined across sessions. The identity line (perfect match) is indicated in red G) Same as F) but with a slightly decreased noise term (85% of original).

**Fig 17.**
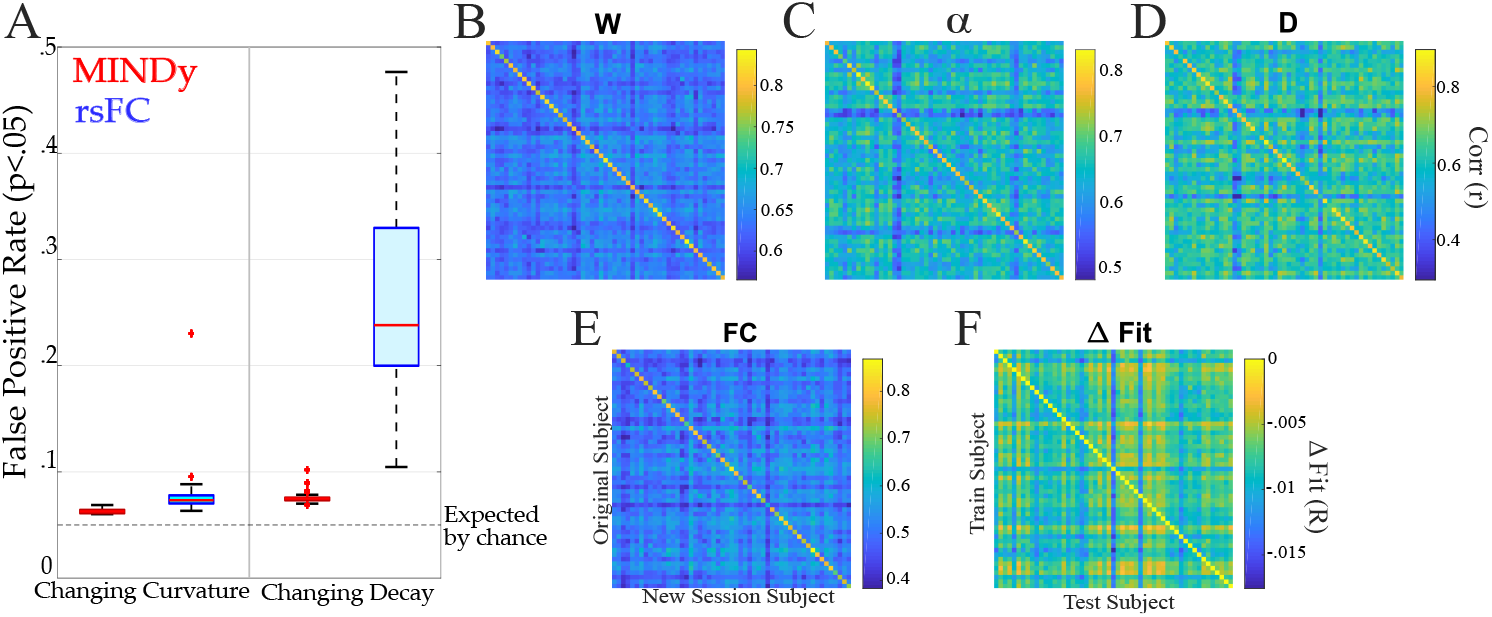
MINDy differentiates model parameters between individuals and identifies the source of individual differences. A) Changing only the curvature (left) or decay (right) parameters of a simulated subject has more impact on the simulated rsFC (blue) then on MINDy weight estimates refit to the new simulated data (red). B) Weight matrices are individualized: weight matrices derived from different scans of the same subject are universally more similar than weights fit to another subject. C,D) same as B) but for the curvature and decay parameters, respectively. E) The rsFC matrix is more similar for different scans of the same subject than between subjects. F) Individualized models better generalize to new data from the same subject than to a new subject.

**Table 6.** Sensitivity of MINDy weights and rsFC to changes in non-connectivity parameters in a ground-truth simulation Performance is measured in terms of false positives=the percentage of connections that change (thresholded by *p* < .05) due to a change in the ground truth model’s curvature/decay. Thus, lower values indicate less false positives (less sensitivity) due to non-connectivity variables. We denote significance with ** = *p* < .001, 2-tailed for the contrast Weights minus rsFC.

|  | Weights | FC | paired-t (df=339) |
| --- | --- | --- | --- |
| Changing Curvature | .063 (.002) | .079 (.027) | -10.639** |
| Changing Decay | .076 (.006) | .2749 (.120) | -31.753** |

**Table 7.** Test-retest correlation across scanning sessions for MINDy parameters when sessions were drawn from the same subject or from different subjects. Group-level statistics are present in mean correlation (SD) form. Group level permutation testing (100,000 each) produced *p*^i^*s* ≈ 0 for all parameters vs. chance. Accuracy is in correct assignment for subjects based upon maximal similarity between sessions (e.g. how often is the subject most similar to themselves?). Statistical tests are for the contrast weight vs. FC. W/in=within subject, Btwn=between subject, Diff=w/in subject minus between, Acc.=accuracy. We denote significance with * = *p* < .05 and ** = *p* < .001, tailed and Bonferroni corrected

| | FC | Weight(W) | t(mean) | Curv($\alpha$) | Decay(D) |
| --- | --- | --- | --- | --- | --- |
| W/in. | .757(.052) | .802(.018) | 7.883** | .772(.029) | .798(.062) |
| Btwn. | .511(.044) | .635(.021) | 38.143** | .637(.041) | .600(.069) |
| Diff. | .246(.038) | .167(.019) | -23.076** | .135(.023) | .198(.052) |
| Acc. | 100 | 100 | 0 | 100 | 94.3 |

**Fig 18.**
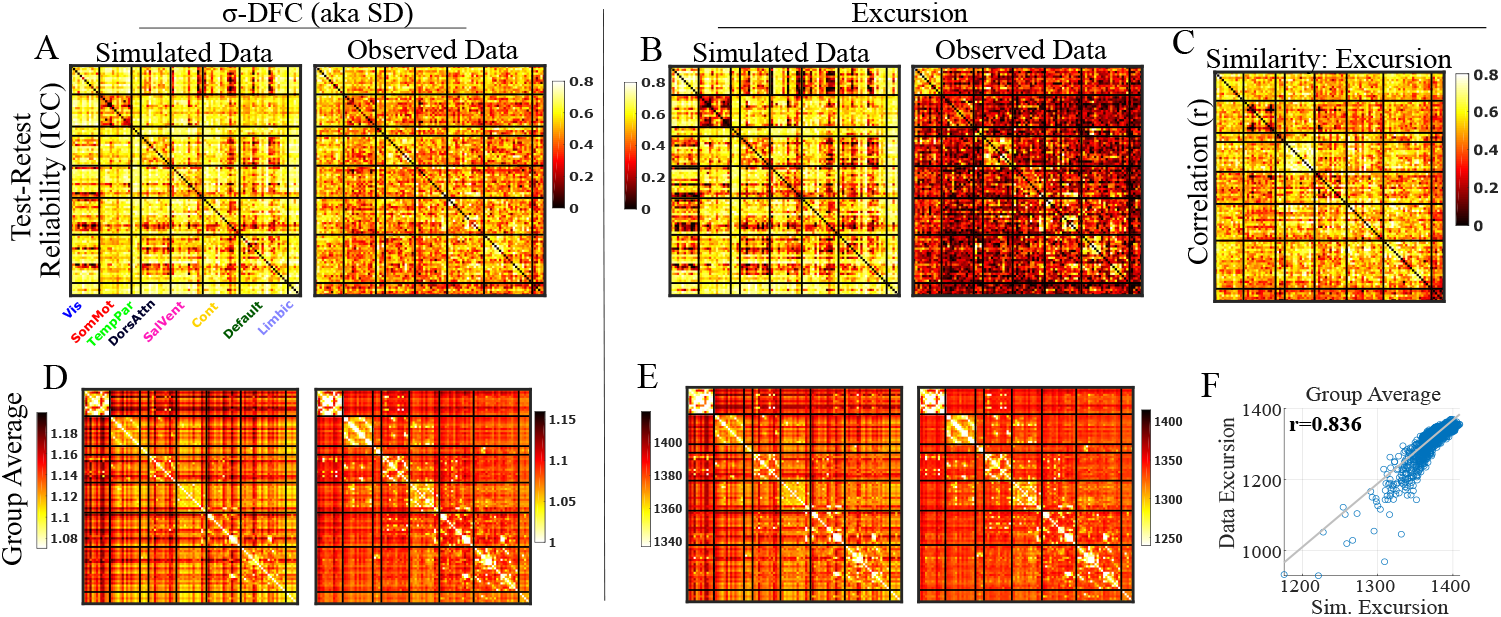
MINDy reproduces reliable, accurate estimates of dynamic functional connectivity (DFC). We use *σ*-DFC to disambiguate the standard-deviation measure for DFC (e.g. [61],[49]) from other uses of standard-deviation. A) Data simulated from test-retest models (models fit to separate sessions) has at least as high reliability on average as the original data for *σ*-DFC. B) Same as A) but for the excursion metric of DFC. C) Correlation between observed and simulated excursion across subjects by region-pair (combining across scanning sessions). D) Predicted group average *σ*-DFC for the model simulations (left) and recorded data (right) combining across scanning sessions. E) Same as D) but for excursion. F) Correlation between observed and predicted group-average excursion across region-pairs. The *σ*-DFC analogues of C and F are reported in the main text (Fig. 6)

**Table 8.** Comparing the test-retest reliability and pre-processing sensitivity of the MINDy connectivity parameter and the resting state functional connectivity. The pipelines correspond to using motion without CompCor or GSR correction, using motion + CompCor or using motion + CompCor + GSR (default). Results are presented in mean(SD) form for the group distribution of individual test-retest correlations or correlations between different levels of preprocessing applied to the same session. Statistical tests consisted of paired t-tests for the mean correlation, and F-tests for testing heterogeneity of variance. Results generally favored the MINDy connectivity matrix over the FC matrix (greater mean reliability and less variation) but the absolute differences, although highly statistically significant, are not profound. We denote significance with * = *p* < .05 and ** = *p* < .001, 2-tailed and Bonferroni corrected.

|  | Weights | FC | t (mean) | F (var) |
| --- | --- | --- | --- | --- |
| Motion | .8096 (.0268) | .7909 (.0619) | 3.1507* | 5.3219** |
| CompCor | .8039 (.0185) | .7481 (.0504) | 10.5096** | 7.378** |
| GSR | .8021 (.0180) | .7571 (.0523) | 7.8834** | 8.4200** |
| M vs. C | .9323 (.0178) | .8512 (.0582) | 13.6332** | 10.7247** |
| M vs. G | .9116 (.0191) | .7910 (.0776) | 14.1496** | 16.4823** |
| C vs. G | .9736 (.0037) | .9669 (.0143) | 4.2162** | 14.6271** |

**Fig 19.**
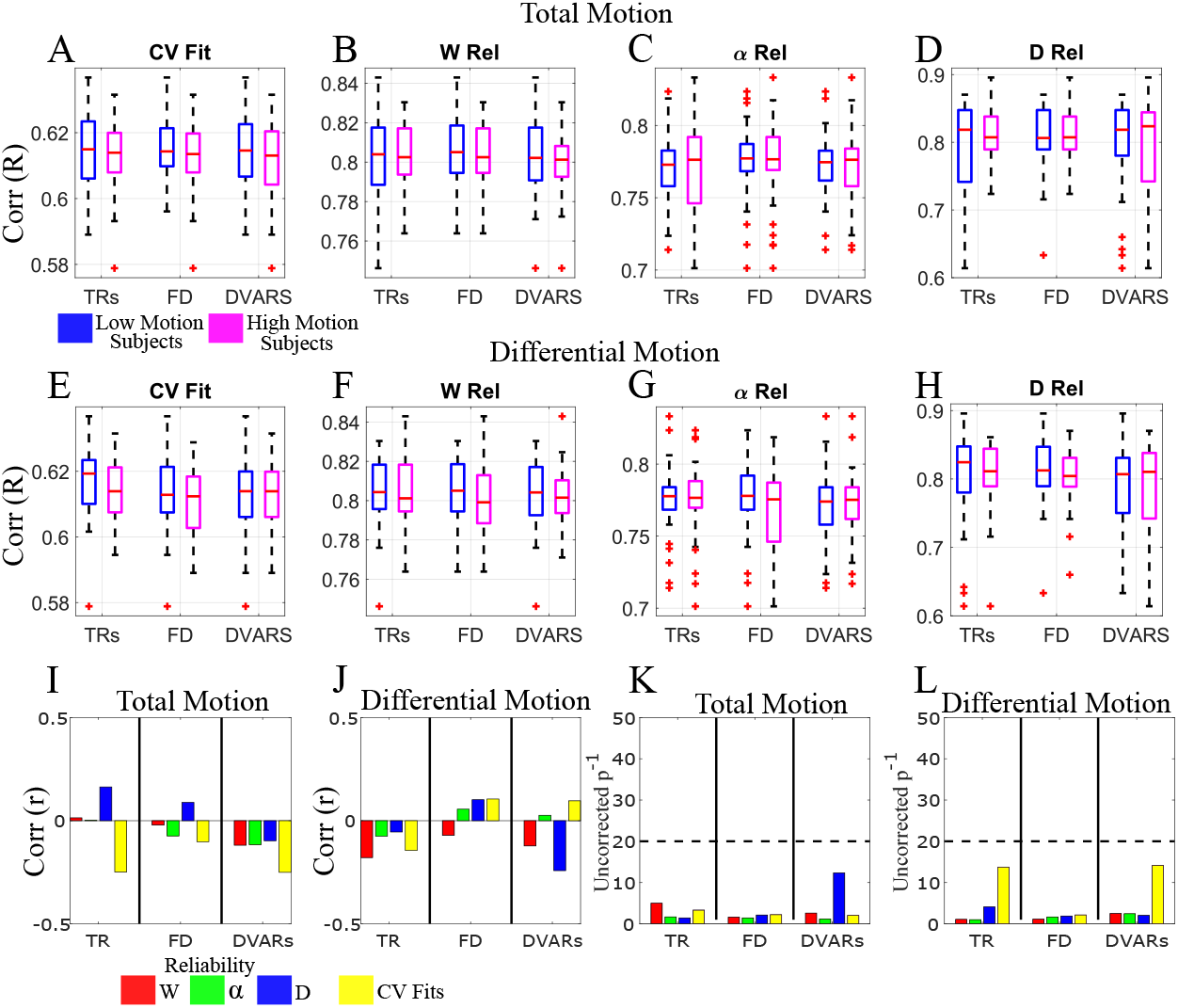
After pre-processing, MINDy fits are robust to motion. Fitting performance was measured by the cross-validated goodness of fit (A,E) and the reliability for each parameter (B-D,F-H). Individual differences in motion were quantified by either membership in median-split high vs. low motion groups (first two rows) or as a continuous variable (bottom row). Groups were assigned for each combination of motion measurement (number of TRs censored, median Framewise Displacement, or Median Absolute Deviation (MAD) of DVARS) and motion type: either the total motion of a subject averaged across scanning sessions (A-D, I,K) or the absolute difference in motion artifact between sessions (E-H,J,L). There was no significant relationship with motion as a discrete characteristic (e.g. high vs. low: A-H) or as a continuous characteristic: group level correlations between motion measures and fitting performance in (I,J) and the associated (uncorrected) inverse *p*-values in (K,L).

**Table 9.** MINDy performance in inverting the weight matrix and its asymmetries in cases of model mismatch. Ground truth models were either a tanh rate-model downsampled to the fMRI TR, a rate model with spatially heterogeneous hemodynamic response functions, or a neural mass model using the nonlinear Balloon-Windkessel model of hemodynamics.

| Model | Full $W$ | $W - W^T$ | N |
| --- | --- | --- | --- |
| Rate-Model | 0.949 (0.009) | 0.971 (0.007) | 1700 |
| Rate+spatial HRF | 0.754 (0.026) | 0.770 (0.027) | 1680 |
| Neural Mass + B-W | 0.642 (0.032) | 0.567 (0.036) | 1480 |

**Fig 20.**
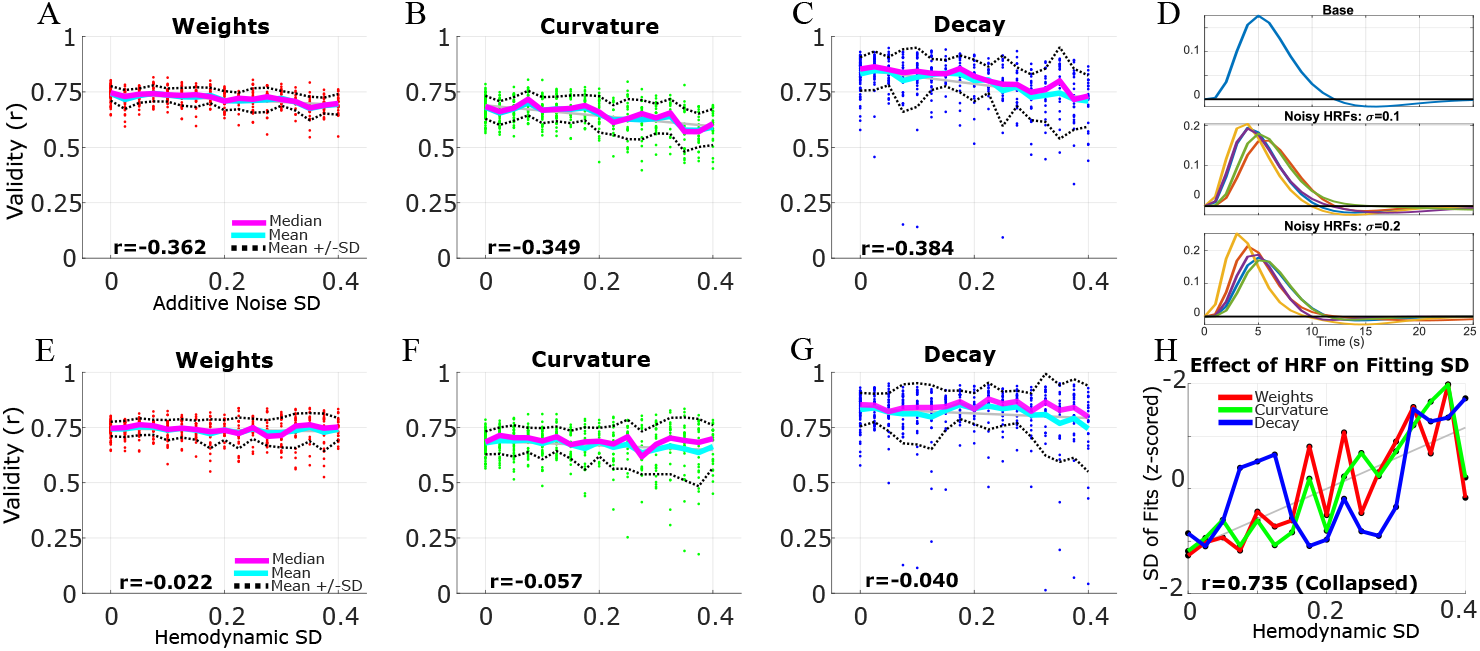
Sensitivity to various forms of noise in data (Same as Fig. 7 A,B, but with 17 levels of noise). A-C) MINDy estimates all parameters of ground-truth models accurately even in the presence of additive measurement noise. D) Examples of how increasing the variability of hemodynamic parameters changes the shape of randomly drawn hemodynamic response functions (HRF). E-G) Hemodynamic variability does not alter the mean performance of MINDy estimates. H) Hemodynamic variability decreases the consistency of MINDy performance (more variable correlations with ground truth).

**Fig 21.**
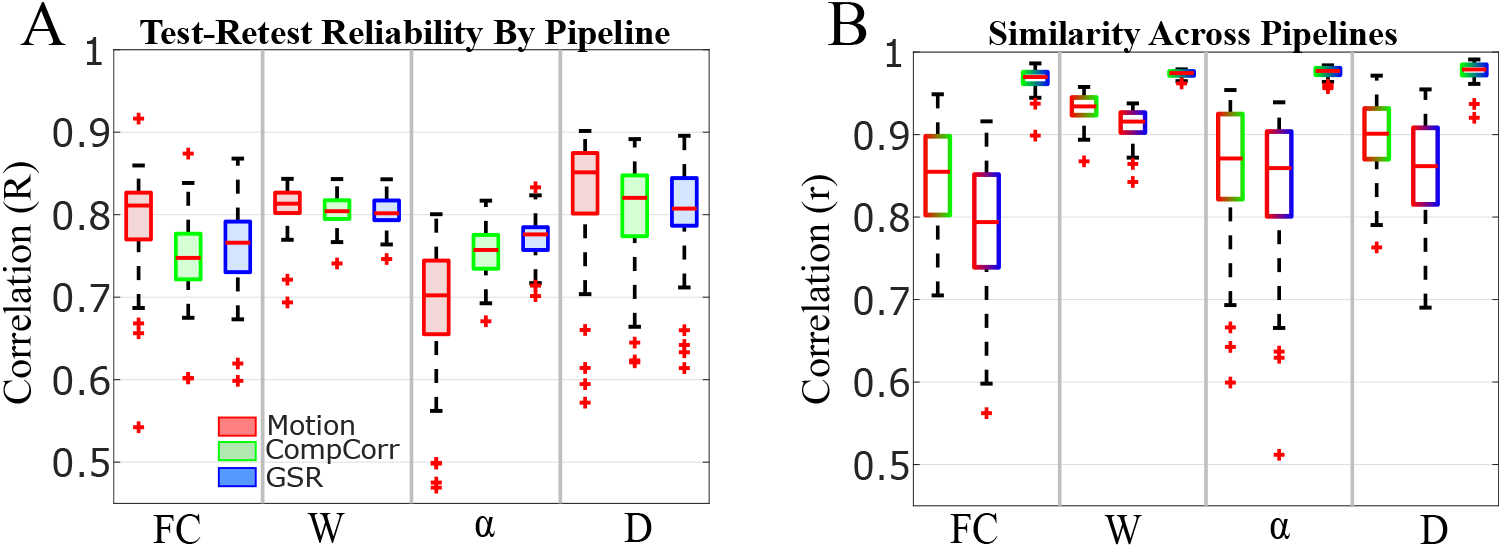
MINDy estimates are robust to secondary preprocessing choices: Motion-only (Red), Motion+CompCor (Green), Motion+CompCor+GSR (Red). A) Test-retest reliability of rsFC and the MINDy weights (W), curvature (*α*) and Decay (D) parameters by secondary preprocessing pipeline. B) Correlation between parameter estimates for different preprocessing pipelines applied to the same scanning data (single session). Color of the bar indicates which pipelines are being compared: R+G indicates similarity between Motion-only and CompCor estimates, R+B indicates Motion-only vs. GSR, G+B indicates CompCor vs. GSR (results are displayed in this order left to right within each column)

**Table 10.** Definition of variables (“Name”) used in MINDy gradient calculations and their interpretation (“Meaning”). The term *n*_*Batch*_ denotes the number of samples (time-points) included in each minibatch (the training data for a given NADAM iteration).

| Name | Equation | Meaning |
| --- | --- | --- |
| $W$ | $W_S + W_L$ | Full weights |
| $P_1(x)$ | $\sqrt{\xi^{-2} + x \circ (x + b^{-1})}$ | Part of $\psi$ |
| $P_2(x)$ | $\sqrt{\xi^{-2} + x \circ (x - b^{-1})}$ | Part of $\psi$ |
| $\psi(x, \varepsilon)$ | $b(P_1 - P_2)$ | Transfer Function ($\psi$) |
| $D$ | $D_2^2 + D_{min}$ | Full Decay |
| $R$ | $dX - W_K \psi(x) + Dx$ | Residual Error |
| $Q$ | $W_K^T R$ | $2 \frac{\partial R}{\partial \psi}$ |
| $Z$ | $R \psi(x)^T / n_{Batch}$ | $E_T[\frac{\partial R}{\partial W}]$ |
| $Y$ | $Z - \lambda_4 W_L$ | $\frac{\partial J}{\partial W_L}$ |

The second change of variable served to linearize the effects of the nonlinear curvature p√arameter *α*. Rather than explicitly fitting *α* we fit the variable 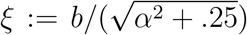 which satisfies 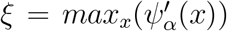. This transformation smooths the relation between the nonlinear parameter (*ξ* instead of *α*) and its effects on the model’s vector field. The new parameter *ξ* is constrained to be smaller than or equal to 2*b* so that *α*^2^ ≥ 0. Efficient gradient calculations were performed by first calculating the variables in Table 10 and then calculating gradients as in Table 11. In all cases MINDy was run for 5000 iterations with batch size 300 (300 time-points used in each iteration).

**Table 11.** Equations used to efficiently calculate MINDy parameter gradients. These equations leverage the additional variables defined in SI Tab. 10. Note that the decay parameter is updated in terms of its square-root (*D*_2_) and the curvature parameter is updated in terms of the linearized form *ξ*.

| Parameter | Negative Error Gradient ($-\frac{\partial J}{\partial \omega}$) |
| --- | --- |
| $W_S$ | $Z - \lambda_1 \text{sgn}(W_S) - \lambda_4 \text{diag}(\text{sgn}(W_S))$ |
| $W_1$ | $YW_2^T - \lambda_3 \text{sgn}(W_1)$ |
| $W_2$ | $W_1Y - \lambda_3 \text{sgn}(W_2)$ |
| $\xi$ | $-\xi^{-3}bE_T[Q \circ (1/P_1 - 1/P_2)]$ |
| $D_2$ | $-2D_2 \circ E_T[R \circ X]$ |

**Table 12.**
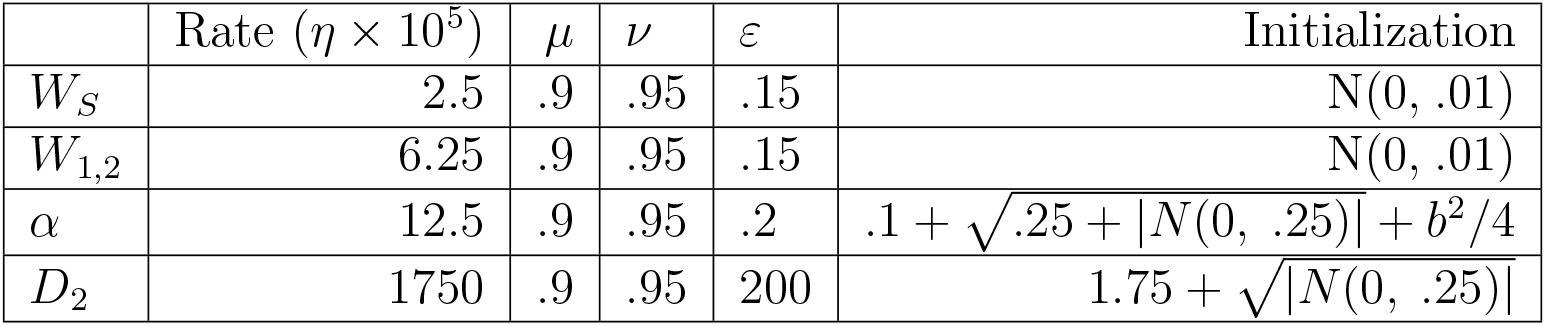
NADAM hyperparametes for each MINDy parameter and the distributions used to initialize each parameter. NADAM hyperparameters consist of the update rate (“Rate”), decay rate of gradients (*μ*), decay rate of squared-gradients (*ν*), and regularization term *∊*.

**Table 13.**
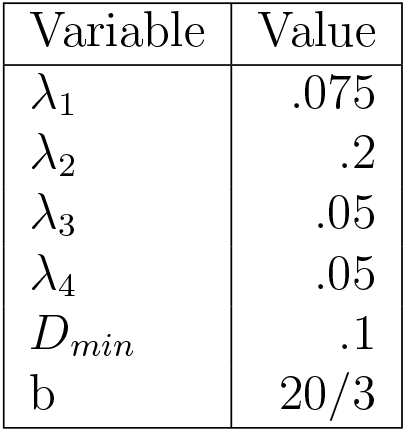
Chosen values for other hyperparameters used in MINDy. These (non-NADAM) hyperparameters consist of the four regularization terms (*λ*_*i*_) in the cost function (Eq. 4), the minimum allowable value for *D* (*D*_*min*_), and the scaling factor of the transfer-function (*b*)

| Variable | Value |
| --- | --- |
| $\lambda_1$ | .075 |
| $\lambda_2$ | .2 |
| $\lambda_3$ | .05 |
| $\lambda_4$ | .05 |
| $D_{min}$ | .1 |
| $b$ | 20/3 |

## Acknowledgments

MS was funded by NSF-DGE-1143954 from the US National Science Foundation. TB acknowledges R37 MH066078 from the US National Institute of Health. SC holds a Career Award at the Scientific Interface from the Burroughs-Wellcome Fund. Portions of this work were supported by AFOSR 15RT0189, NSF ECCS 1509342 and NSF CMMI 1537015, NSF NCS-FO 1835209 and NIMH Administrative Supplement MH066078-15S1 from the US Air Force Office of Scientific Research, US National Science Foundation, and US National Institute of Mental Health, respectively.

## References

[1] H. Markram, The blue brain project, Nature Reviews Neuroscience 7 (2006) 153–160. doi:10.1038/nrn1848.

[2] H. Markram, K. Meier, T. Lippert, S. Grillner, R. Frackowiak, S. Dehaene, A. Knoll, H. Sompolinsky, K. Verstreken, J. DeFelipe, S. Grant, J.-P. Changeux, A. Saria, Introducing the human brain project, Procedia Computer Science 7 (2011) 39 – 42. doi: 10.1016/j.procs.2011.12.015.

[3] H. Okano, E. Sasaki, T. Yamamori, A. Iriki, T. Shimogori, Y. Yamaguchi, K. Kasai, A. Miyawaki, Brain/minds: A japanese national brain project for marmoset neuroscience, Neuron 92 (3) (2016) 582 – 590. doi:10.1016/j.neuron.2016.10.018.

[4] D. C. V. Essen, S. M. Smith, D. M. Barch, T. E. Behrens, E. Yacoub, K. Ugurbil, The wu-minn human connectome project: An overview, NeuroImage 80 (2013) 62 – 79. doi:10.1016/j.neuroimage.2013.05.041.

[5] B. Biswal, F. Z. Yetkin, V. M. Haughton, J. S. Hyde, Functional connectivity in the motor cortex of resting human brain using echo-planar mri, Magnetic Resonance in Medicine 34 (4) (1995) 537–541. doi:10.1002/mrm.1910340409.

[6] R. L. Buckner, F. M. Krienen, B. T. T. Yeo, Opportunities and limitations of intrinsic functional connectivity mri, Nature Neuroscience 16 (2013) 832–837. doi:10.1038/nn.3423.

[7] A. Razi, M. L. Seghier, Y. Zhou, P. McColgan, Large-scale dcms for resting-state fmri, Network Neuroscience 1 (3) (2017). doi:10.1162/NETN\_a\_00015.

[8] M. Breakspear, Dynamic models of large-scale brain activity, Nature Neuroscience 20 (2017) 340–352. doi:10.1038/nn.4497.

[9] P. Sanz Leon, S. Knock, M. Woodman, L. Domide, J. Mersmann, A. McIntosh, V. Jirsa, The virtual brain: a simulator of primate brain network dynamics, Frontiers in Neuroinformatics 7 (2013) 10. doi:10.3389/fninf.2013.00010.

[10] M. Schirner, S. Rothmeier, V. K. Jirsa, A. R. McIntosh, P. Ritter, An automated pipeline for constructing personalized virtual brains from multimodal neuroimaging data, NeuroImage 117 (2015) 343 – 357. doi:10.1016/j.neuroimage.2015.03.055.

[11] V. Jirsa, T. Proix, D. Perdikis, M. Woodman, H. Wang, J. Gonzalez-Martinez, C. Bernard, C. Bènar, M. Guye, P. Chauvel, F. Bartolomei, The virtual epileptic patient: Individualized whole-brain models of epilepsy spread, NeuroImage 145 (2017) 377 – 388. doi:10.1016/j.neuroimage.2016.04.049.

[12] M. Demirtás, J. B. Burt, M. Helmer, J. L. Ji, B. D. Adkinson, M. F. Glasser, D. C. Van Essen, S. N. Sotiropoulos, A. Anticevic, J. D. Murray, Hierarchical heterogeneity across human cortex shapes large-scale neural dynamics, Neuron 101 (6) (2019) 1181–1194. doi:10.1016/j.neuron.2019.01.017.

[13] P. Wang, R. Kong, X. Kong, R. Liègeois, C. Orban, G. Deco, M. P. van den Heuvel, B. Thomas Yeo, Inversion of a large-scale circuit model reveals a cortical hierarchy in the dynamic resting human brain, Science Advances 5 (1) (2019). doi:10.1126/sciadv.aat7854.

[14] K. Friston, L. Harrison, W. Penny, Dynamic causal modelling, NeuroImage 19 (4) (2003) 1273 – 1302. doi:10.1016/S1053-8119(03)00202-7.

[15] C. Tu, R. P. Rocha, M. Corbetta, S. Zampieri, M. Zorzi, S. Suweis, Warnings and caveats in brain controllability, NeuroImage 176 (2018) 83 – 91. doi:10.1016/j.neuroimage.2018.04.010.

[16] S. Ryali, K. Supekar, T. Chen, V. Menon, Multivariate dynamical systems models for estimating causal interactions in fmri, NeuroImage 54 (2) (2011) 807 – 823. doi:10.1016/j.neuroimage.2010.09.052.

[17] A. Roebroeck, E. Formisano, R. Goebel, The identification of interacting networks in the brain using fmri: Model selection, causality and deconvolution, NeuroImage 58 (2) (2011) 296 – 302. doi: 10.1016/j.neuroimage.2009.09.036.

[18] G. Lohmann, K. Erfurth, K. Müller, R. Turner, Critical comments on dynamic causal modelling, NeuroImage 59 (3) (2012) 2322 – 2329. doi:10.1016/j.neuroimage.2011.09.025.

[19] S. Frässle, E. I. Lomakina, A. Razi, K. J. Friston, J. M. Buhmann, K. E. Stephan, Regression dcm for fmri, Neuroimage 155 (2017) 406–421.

[20] M. F. Glasser, T. S. Coalson, E. C. Robinson, C. D. Hacker, E. Harwell, John andYacoub, K. Ugurbil, J. Andersson, C. F. Beckmann, M. Jenkinson, S. M. Smith, D. C. Van Essen, A multi-modal parcellation of human cerebral cortex, Nature 536 (2016) 171–178. doi: 10.1038/nature18933.

[21] A. Schaefer, R. Kong, E. M. Gordon, T. O. Laumann, X.-N. Zuo, A. J. Holmes, S. B. Eickhoff, B. T. Yeo, Local-global parcellation of the human cerebral cortex from intrinsic functional connectivity mri, Cerebral Cortex (2017) 1–20doi:10.1093/cercor/bhx179.

[22] C. J. Honey, O. Sporns, L. Cammoun, X. Gigandet, J. P. Thiran, R. Meuli, P. Hagmann, Predicting human resting-state functional connectivity from structural connectivity, Proceedings of the National Academy of Sciences 106 (6) (2009) 2035–2040. doi:10.1073/pnas.0811168106.

[23] S. Knock, A. McIntosh, O. Sporns, R. K’otter, P. Hagmann, V. Jirsa, The effects of physiologically plausible connectivity structure on local and global dynamics in large scale brain models, Journal of Neuroscience Methods 3 (1) (2009) 86–94.

[24] E. A. Ashley, The Precision Medicine Initiative: A New National Effort, JAMA 313 (21) (2015) 2119–2120. doi:10.1001/jama.2015.3595.

[25] B. M. Psaty, O. M. Dekkers, R. S. Cooper, Comparison of 2 Treatment Models: Precision Medicine and Preventive Medicine, JAMA 320 (8) (2018) 751–752. doi:10.1001/jama.2018.8377.

[26] T. D. Satterthwaite, C. H. Xia, D. S. Bassett, Personalized neuroscience: Common and individual-specific features in functional brain networks, Neuron 98 (2) (2018) 243 – 245. doi:10.1016/j.neuron.2018.04.007.

[27] J. J. Hopfield, Neurons with graded response have collective computational properties like those of two-state neurons, Proceedings of the National Academy of Sciences 81 (10) (1984) 3088–3092. doi:10.1073/pnas.81.10.3088.

[28] J. S. Siegel, A. Mitra, T. O. Laumann, B. A. Seitzman, M. Raichle, M. Corbetta, A. Z. Snyder, Data quality influences observed links between functional connectivity and behavior, Cerebral Cortex 27 (9) (2017) 4492–4502. doi:10.1093/cercor/bhw253.

[29] N. Weiner, Extrapolation, interpolation, and smoothing of stationary time series: with engineering applications, MIT Press, New York City, 1949.

[30] K. Friston, P. Fletcher, O. Josephs, A. Holmes, M. Rugg, R. Turner, Event-related fmri: Characterizing differential responses, NeuroImage 7 (1) (1998) 30 – 40. doi:10.1006/nimg.1997.0306.

[31] H. R. Wilson, J. D. Cowan, Excitatory and inhibitory interactions in localized populations of model neurons, Biophysical Journal 12 (1) (1972) 1–24. doi:10.1016/S0006-3495(72)86068-5.

[32] G. Deco, V. K. Jirsa, A. R. McIntosh, Emerging concepts for the dynamical organization of resting-state activity in the brain, Nature Reviews Neuroscience 12 (2011) 43–56. doi:10.1038/nrn2961.

[33] A. C. Marreiros, J. Daunizeau, S. J. Kiebel, K. J. Friston, Population dynamics: Variance and the sigmoid activation function, NeuroImage 42 (1) (2008) 147 – 157. doi:10.1016/j.neuroimage.2008.04.239.

[34] T. Dozat, Incorporating nesterov momentum into adam, Proceedings of 4th International Conference on Learning Representations, Workshop Track, 2016 (2016).

[35] D. L. Donoho, For most large underdetermined systems of linear equations the minimal l1 norm solution is also the sparsest solution, Communications on Pure and Applied Mathematics 59 (6) (2006) 797–829. doi:10.1002/cpa.20132.

[36] F. Mastrogiuseppe, S. Ostojic, Linking connectivity, dynamics, and computations in low-rank recurrent neural networks, Neuron 99 (2018) 609–623. doi:10.1016/j.neuron.2018.07.003.

[37] M. F. Glasser, S. N. Sotiropoulos, J. A. Wilson, T. S. Coalson, B. Fischl, J. L. Andersson, J. Xu, S. Jbabdi, M. Webster, J. R. Polimeni, D. C. V. Essen, M. Jenkinson, The minimal preprocessing pipelines for the human connectome project, NeuroImage 80 (2013) 105–124. doi: 10.1016/j.neuroimage.2013.04.127.

[38] L. Griffanti, G. Salimi-Khorshidi, C. F. Beckmann, E. J. Auerbach, G. Douaud, C. E. Sexton, E. Zsoldos, K. P. Ebmeier, N. Filippini, C. E. Mackay, S. Moeller, J. Xu, E. Yacoub, G. Baselli, K. Ugurbil, K. L. Miller, S. M. Smith, Ica-based artefact removal and accelerated fmri acquisition for improved resting state network imaging, NeuroImage 95 (2014) 232 – 247. doi:https://doi.org/10.1016/j.neuroimage.2014.03.034.

[39] G. Salimi-Khorshidi, G. Douaud, C. F. Beckmann, M. F. Glasser, L. Griffanti, S. M. Smith, Automatic denoising of functional mri data: Combining independent component analysis and hierarchical fusion of classifiers, NeuroImage 90 (2014) 449 – 468. doi:10.1016/j.neuroimage.2013.11.046.

[40] J. D. Power, B. L. Schlaggar, S. E. Petersen, Recent progress and outstanding issues in motion correction in resting state fmri, NeuroImage 105 (2015) 536 – 551. doi:https://doi.org/10.1016/j.neuroimage.2014.10.044.

[41] J. D. Power, K. A. Barnes, A. Z. Snyder, B. L. Schlaggar, S. E. Petersen, Spurious but systematic correlations in functional connectivity mri networks arise from subject motion, NeuroImage 59 (3) (2012) 2142 – 2154. doi:10.1016/j.neuroimage.2011.10.018.

[42] Y. Behzadi, K. Restom, J. Liau, T. T. Liu, A component based noise correction method (compcor) for bold and perfusion based fmri, NeuroImage 37 (1) (2007) 90 – 101. doi:10.1016/j.neuroimage.2007.04.042.

[43] G. Aguirre, E. Zarahn, M. D’Esposito, The variability of human, bold hemodynamic responses, NeuroImage 8 (4) (1998) 360 – 369. doi:10.1006/nimg.1998.0369.

[44] B. Fischl, Freesurfer, NeuroImage 62 (2) (2012) 774 – 781. doi: 10.1016/j.neuroimage.2012.01.021.

[45] B. T. Thomas Yeo, F. M. Krienen, J. Sepulcre, M. R. Sabuncu, D. Lashkari, M. Hollinshead, J. L. Roffman, J. W. Smoller, L. Zöllei, J. R. Polimeni, B. Fischl, H. Liu, R. L. Buckner, The organization of the human cerebral cortex estimated by intrinsic functional connectivity, Journal of Neurophysiology 106 (3) (2011) 1125–1165. doi:10.1152/jn.00338.2011.

[46] P. W. Holland, R. E. Welsch, Robust regression using iteratively reweighted least-squares, Communications in Statistics - Theory and Methods 6 (9) (1977) 813–827. doi:10.1080/03610927708827533.

[47] K. Friston, A. Mechelli, R. Turner, C. Price, Nonlinear responses in fmri: The balloon model, volterra kernels, and other hemodynamics, NeuroImage 12 (4) (2000) 466 – 477. doi:10.1006/nimg.2000.0630.

[48] E. C. Hansen, D. Battaglia, A. Spiegler, G. Deco, V. K. Jirsa, Functional connectivity dynamics: Modeling the switching behavior of the resting state, NeuroImage 105 (2015) 525 – 535. doi: 10.1016/j.neuroimage.2014.11.001.

[49] C. Zhang, S. A. Baum, V. R. Adduru, B. B. Biswal, A. M. Michael, Test-retest reliability of dynamic functional connectivity in resting state fmri, NeuroImage 183 (2018) 907 – 918. doi:10.1016/j.neuroimage.2018.08.021.

[50] R. Engle, Dynamic conditional correlation: A simple class of multivariate generalized autoregressive conditional heteroskedasticity models, Journal of Business & Economic Statistics 20 (3) (2002) 339–350. doi:10.1198/073500102288618487.

[51] P. Shrout, J. Fleiss, Intraclass correlations: uses in assessing rater reliability, Psychological bulletin 86 (2) (1979) 420–428.

[52] H. Shou, A. Eloyan, S. Lee, V. Zipunnikov, A. N. Crainiceanu, M. B. Nebel, B. Caffo, M. A. Lindquist, C. M. Crainiceanu, Quantifying the reliability of image replication studies: The image intraclass correlation coefficient (i2c2), Cognitive, Affective, & Behavioral Neuroscience 13 (4) (2013) 714–724. doi:10.3758/s13415-013-0196-0.

[53] T. O. Laumann, E. M. Gordon, B. Adeyemo, A. Z. Snyder, S. J. Joo, M.-Y. Chen, A. W. Gilmore, K. B. McDermott, S. M. Nelson, N. U. Dosenbach, B. L. Schlaggar, J. A. Mumford, R. A. Poldrack, S. E. Petersen, Functional system and areal organization of a highly sampled individual human brain, Neuron 87 (3) (2015) 657 – 670. doi:10.1016/j.neuron.2015.06.037.

[54] E. M. Gordon, T. O. Laumann, A. W. Gilmore, D. J. Newbold, D. J. Greene, J. J. Berg, M. Ortega, C. Hoyt-Drazen, C. Gratton, H. Sun, J. M. Hampton, R. S. Coalson, A. L. Nguyen, K. B. McDermott, J. S. Shimony, A. Z. Snyder, B. L. Schlaggar, S. E. Petersen, S. M. Nelson, N. U. Dosenbach, Precision functional mapping of individual human brains, Neuron 95 (4) (2017) 791 – 807.e7. doi:10.1016/j.neuron.2017.07.011.

[55] E. S. Finn, X. Shen, D. Scheinost, M. D. Rosenberg, J. Huang, M. M. Chun, X. Papademetris, R. T. Constable, Functional connectome fingerprinting: identifying individuals using patterns of brain connectivity, Nature Neureoscience 18 (2015) 1664–1671. doi:10.1038/nn.4135.

[56] A. D. Friederici, S. M. Gierhan, The language network, Current Opinion in Neurobiology 23 (2) (2013) 250 – 254. doi:10.1016/j.conb.2012.10.002.

[57] R. M. Hutchison, T. Womelsdorf, E. A. Allen, P. A. Bandettini, V. D. Calhoun, M. Corbetta, S. D. Penna, J. H. Duyn, G. H. Glover, J. Gonzalez-Castillo, D. A. Handwerker, S. Keilholz, V. Kiviniemi, D. A. Leopold, F. de Pasquale, O. Sporns, M. Walter, C. Chang, Dynamic functional connectivity: Promise, issues, and interpretations, NeuroImage 80 (2013) 360 – 378. doi:10.1016/j.neuroimage.2013.05.079.

[58] E. A. Allen, E. Damaraju, S. M. Plis, E. B. Erhardt, T. Eichele, V. D. Calhoun, Tracking whole-brain connectivity dynamics in the resting state, Cerebral Cortex 24 (3) (2014) 663–676. doi:10.1093/cercor/bhs352.

[59] R. F. Betzel, M. Fukushima, Y. He, X.-N. Zuo, O. Sporns, Dynamic fluctuations coincide with periods of high and low modularity in restingstate functional brain networks, NeuroImage 127 (2016) 287–297. doi: 10.1016/j.neuroimage.2015.12.001.

[60] M. G. Preti, T. A. Bolton, D. V. D. Ville, The dynamic functional connectome: State-of-the-art and perspectives, NeuroImage 160 (2017) 41 – 54. doi:10.1016/j.neuroimage.2016.12.061.

[61] A. S. Choe, M. B. Nebel, A. D. Barber, J. R. Cohen, Y. Xu, J. J. Pekar, B. Caffo, M. A. Lindquist, Comparing test-retest reliability of dynamic functional connectivity methods, NeuroImage 158 (2017) 155 – 175. doi:10.1016/j.neuroimage.2017.07.005.

[62] G. Aguirre, E. Zarahn, M. D’Esposito, The inferential impact of global signal covariates in functional neuroimaging analyses, NeuroImage 8 (3) (1998) 302 – 306. doi:doi.org/10.1006/nimg.1998.0367.

[63] M. F. Singh, A. Wang, T. S. Braver, S. Ching, Scalable surrogate deconvolution for identification of partially-observable systems and brain modeling, bioRxiv (2020). doi:10.1101/2020.03.20.000661. URL https://www.biorxiv.org/content/early/2020/03/23/2020.03.20.000661

[64] F.-H. Lin, J. R. Polimeni, J.-F. L. Lin, K. W.-K. Tsai, Y.-H. Chu, P.-Y. Wu, Y.-T. Li, Y.-C. Hsu, S.-Y. Tsai, W.-J. Kuo, Relative latency and temporal variability of hemodynamic responses at the human primary visual cortex, NeuroImage 164 (2018) 194 – 201. doi:10.1016/j.neuroimage.2017.01.041.

[65] K. E. Stephan, L. Kasper, L. M. Harrison, J. Daunizeau, H. E. [den Ouden], M. Breakspear, K. J. Friston, Nonlinear dynamic causal models for fmri, NeuroImage 42 (2) (2008) 649 – 662. doi:10.1016/j.neuroimage.2008.04.262.

[66] K. Friston, K. H. Preller, C. Mathys, H. Cagnan, J. Heinzle, A. Razi, P. Zeidman, Dynamic causal modelling revisited, NeuroImage 199 (2019) 730 – 744. doi:10.1016/j.neuroimage.2017.02.045.

[67] T. O. Laumann, A. Z. Snyder, A. Mitra, E. M. Gordon, C. Gratton, B. Adeyemo, A. W. Gilmore, S. M. Nelson, J. J. Berg, D. J. Greene, J. E. McCarthy, E. Tagliazucchi, H. Laufs, B. L. Schlaggar, N. U. F. Dosenbach, S. E. Petersen, On the stability of bold fmri correlations, Cerebral Cortex 27 (10) (2017) 4719–4732. doi:10.1093/cercor/bhw265.

[68] M. Kafashan, B. J. A. Palanca, S. Ching, Dimensionality reduction impedes the extraction of dynamic functional connectivity states from fmri recordings of resting wakefulness, Journal of Neuroscience Methods 293 (2018) 151 – 161. doi:10.1016/j.jneumeth.2017.09.013.

[69] C. J. Honey, R. Kötter, M. Breakspear, O. Sporns, Network structure of cerebral cortex shapes functional connectivity on multiple time scales, Proceedings of the National Academy of Sciences 104 (24) (2007) 10240–10245. doi:10.1073/pnas.0701519104.

[70] B. J. He, Spontaneous and task-evoked brain activity negatively interact, Journal of Neuroscience 33 (11) (2013) 4672–4682. doi: 10.1523/JNEUROSCI.2922-12.2013.

[71] M. W. Cole, T. Ito, D. S. Bassett, D. H. Schultz, Activity flow over resting-state networks shapes cognitive task activations, Nature Neuroscience 19 (12) (2016) 1718–1726. doi:10.1038/nn.4406.

[72] A. Wang, M. F. Singh, J. Etzel, T. Braver, Enhancing task fmri preprocessing via whole-brain neural modeling of intrinsic activity dynamics, in: Organization for Human Brain Mapping (Virtual). 2020. 2020.

[73] A. Mitra, A. Z. Snyder, E. Tagliazucchi, H. Laufs, M. E. Raichle, Propagated infra-slow intrinsic brain activity reorganizes across wake and slow wave sleep, eLife 4 (2015) e10781. doi:10.7554/eLife.10781.

[74] A. Mitra, A. Kraft, P. Wright, B. Acland, A. Z. Snyder, Z. Rosenthal, L. Czerniewski, A. Bauer, L. Snyder, J. Culver, J.-M. Lee, M. E. Raichle, Spontaneous infra-slow brain activity has unique spatiotemporal dynamics and laminar structure, Neuron 98 (2) (2018) 297 – 305.e6. doi:doi.org/10.1016/j.neuron.2018.03.015.

[75] A. Kucyi, K. D. Davis, Dynamic functional connectivity of the default mode network tracks daydreaming, NeuroImage 100 (2014) 471 – 480. doi:10.1016/j.neuroimage.2014.06.044.

[76] M. D. Fox, A. Z. Snyder, J. L. Vincent, M. E. Raichle, Intrinsic fluctuations within cortical systems account for intertrial variability in human behavior, Neuron 56 (1) (2007) 171 – 184. doi:10.1016/j.neuron.2007.08.023.

[77] M. Schölvinck, K. Friston, G. Rees, The influence of spontaneous activity on stimulus processing in primary visual cortex, NeuroImage 59 (3) (2012) 2700 – 2708. doi:10.1016/j.neuroimage.2011.10.066.

[78] S. Sadaghiani, J.-B. Poline, A. Kleinschmidt, M. D’Esposito, Ongoing dynamics in large-scale functional connectivity predict perception, Proceedings of the National Academy of Sciences 112 (27) (2015) 8463–8468. doi:10.1073/pnas.1420687112.

[79] T. Schreiber, Measuring information transfer, Phys. Rev. Lett. 85 (2000) 461–464. doi:10.1103/PhysRevLett.85.461.

[80] A. Schroeter, F. Schlegel, A. Seuwen, J. Grandjean, M. Rudin, Specificity of stimulus-evoked fmri responses in the mouse: The influence of systemic physiological changes associated with innocuous stimulation under four different anesthetics, NeuroImage 94 (2014) 372 – 384. doi:10.1016/j.neuroimage.2014.01.046.

[81] F. Schlegel, A. Schroeter, M. Rudin, The hemodynamic response to somatosensory stimulation in mice depends on the anesthetic used: Implications on analysis of mouse fmri data, NeuroImage 116 (2015) 40 – 49. doi:10.1016/j.neuroimage.2015.05.013.

[82] N. Todd, S. Moeller, E. J. Auerbach, E. Yacoub, G. Flandin, N. Weiskopf, Evaluation of 2d multiband epi imaging for high-resolution, whole-brain, task-based fmri studies at 3t: Sensitivity and slice leakage artifacts, NeuroImage 124 (2016) 32 – 42. doi: 10.1016/j.neuroimage.2015.08.056.

[83] G. Deco, V. K. Jirsa, P. A. Robinson, M. Breakspear, K. Friston, The dynamic brain: From spiking neurons to neural masses and cortical fields, PLOS Computational Biology 4 (8) (2008) 1–35. doi:10.1371/journal.pcbi.1000092.

[84] M. Singh, D. Zald, A simple transfer function for nonlinear dendritic integration, Frontiers in Computational Neuroscience 9 (2015) 98. doi:10.3389/fncom.2015.00098.

[85] D. P. Kingma, J. Ba, Adam: A method for stochastic optimization, CoRR abs/1412.6980 (2014). arXiv:1412.6980. URL http://arxiv.org/abs/1412.6980

[86] A. R. Aron, T. W. Robbins, R. A. Poldrack, Inhibition and the right inferior frontal cortex, Trends in Cognitive Sciences 8 (4) (2004) 170 – 177. doi:10.1016/j.tics.2004.02.010.

[87] D. Swick, V. Ashley, A. U. Turken, Left inferior frontal gyrus is critical for response inhibition, BMC Neuroscience 9 (1) (2008) 102. doi:10.1186/1471-2202-9-102.

